# Kinetic assays of DNA polymerase fidelity: a new theoretical perspective beyond Michaelis-Menten kinetics

**DOI:** 10.1101/2020.07.10.196808

**Authors:** Qiu-Shi Li, Yao-Gen Shu, Zhong-Can Ou-Yang, Ming Li

## Abstract

The high fidelity of DNA polymerase (DNAP) is critical for the faithful replication of DNA. There are several quantitative approaches to measure DNAP fidelity. Directly counting the error frequency in the replication products gives the true fidelity but it turns out very hard to implement in practice. Two biochemical kinetic approaches, the steady-state assay and the transient-state assay, were then suggested and widely adopted. In these assays, the error frequency is indirectly estimated by using kinetic theories combined with the measured apparent kinetic rates. However, whether it is equivalent to the true fidelity has never been clarified theoretically, and in particular there are different strategies using these assays to quantify the proofreading efficiency of DNAP but often lead to inconsistent results. In this paper, we make a comprehensive examination on the theoretical foundation of the two kinetic assays, based on the theory of DNAP fidelity recently proposed by us. Our studies show that while the conventional kinetic assays are generally valid to quantify the discrimination efficiency of DNAP, they are valid to quantify the proofreading efficiency of DNAP only when the kinetic parameters satisfy some constraints which will be given explicitly in this paper.These results may inspire more carefully-designed experiments to quantify DNAP fidelity.

## I. INTRODUCTION

The high fidelity of DNA polymerase (DNAP) is critical for faithful replication of genomic DNA. Quantitative studies on DNAP fidelity began in 1960s’ and became an important issue in biochemistry and molecular biology. Intuitively, the DNAP fidelity can be roughly understood as the reciprocal of the overall mismatch (error) frequency when a given DNA template is replicated with both the matched dNTPs (denoted as dRTP or R) and the mismatched dNTPs (denoted as dWTP or W). For instance, by using artificial simple template and conducting the replication reaction with both dRTP and dWTP, one can measure the ratio of the incorporated dRTPs to dWTPs in the final products so as to quantify the overall error frequency[1, 2]. Beyond such overall fidelity, the site-specific fidelity was defined as the reciprocal of the error frequency at individual template sites. In principle, the error frequency at any template site can be directly counted if a sufficient amount of full-length replication products can be collected and sequenced(this will be denoted as true fidelity ℱ in this paper), *e.g.* by using deep sequencing techniques [3, 4]. However, this type of sequencing-based approach always requires a huge workload and was rarely adopted in fidelity assay. It is also hard to specify the sequence-context influences on the fidelity. A much simpler strategy is to only investigate the error frequency at the assigned template site by single-nucleotide incorporation assays. Such assays are conducted for *exo*^−^-DNAP (exonuclease-deficient DNAP), in which dRTP and dWTP compete for the same assigned template site and the amount of the final products containing the incorporated dRTP or dWTP are then determined by gel analysis to give the error frequency, *e.g.* [5, 6]. By designing various template sequences, one can further dissect the sequence-context dependence of the site-specific error frequency. Although the direct measurements seem simple and intuitive, they are actually very challenging since mismatches occur with too low probability to be detected even when heavily-biased dNTP pools are used. Besides, the single-nucleotide incorporation assays do not apply to *exo*^+^-DNAP (exonuclease-efficient DNAP) because the coexistence of the polymerase activity and the exonuclease activity makes the reaction products very complicated and hard to interpret. Hence two alternative kinetic approaches were proposed, inspired by the kinetic proofreading theory of biosynthetic processes proposed by Hopfield [7] and Ninio[8].

The steady-state assay was developed by A. Fersht for *exo*^−^-DNAP, which is based on the Michaelis-Menten kinetics of the incorporation of a single dRTP or dWTP at the same assigned template site[9, 10]. The two incorporation reactions are conducted separately under steady-state conditions to obtain the so-called specificity constant (the quasi-first order rate constant) (*k*_*cat*_/*K*_*m*_)_*R*_ or (*k*_*cat*_/*K*_*m*_)_*W*_ respectively, *k*_*cat*_ is the maximal steady-state turnover rate of dNTP incorporation and *K*_*m*_ is the Michaelis constant. The site-specific fidelity is then characterized as the ratio between the two incorporation velocities, *i.e.* (*k*_*cat*_/*K*_*m*_)_*R*_[dRTP] (*k*_*cat*_/*K*_*m*_)_*W*_ [dWTP] (denoted as steady-state fidelity *f*_*s·s*_). However, this is only an operational definition of DNAP fidelity which is not necessarily equal to the true fidelity (see discussions in Sec.III B). So far as we know, there was only one experiment work which did the comparison between *f*_*s·s*_ and ℱ and indicated their possible equivalence for *exo*^−^-Klenow fragment (KF^−^)[6], but no theoretical works have ever been published to examine the equivalence in general.

Besides the steady-state method, the transient-state kinetic analysis was also proposed to obtain the specificity constant[11, 12]. Under the pre-steady-state condition or the single-turnover condition, one can obtain the parameter *k*_*pol*_/*K*_*d*_ (a substitute for *k*_*cat*_/*K*_*m*_) for the single-nucleotide incorporation reactions with *exo*^−^-DNAP, and define the site-specific fidelity as (*k*_*pol*_/*K*_*d*_)_*R*_[dRTP]/(*k*_*pol*_/*K*_*d*_)_*W*_ [dWTP] (denoted as transient-state fidelity *f*_*t·s*_). But again the relation between *f*_*t·s*_ and ℱ is not yet clarified. Although the experiment has indicated the possible equivalence of *f*_*t·s*_ to ℱ for KF^−^[6], a general theoretical examination is still needed.

Further, these methods fail to unambiguously measure the site-specific fidelity of *exo*^+^-DNAP. For *exo*^+^-DNAP, the total fidelity is assumed to consist of two multiplier factors. The first is the initial discrimination *f*_*ini*_ contributed solely by the polymerase domain, which is often characterized by *f*_*s·s*_ or *f*_*t·s*_. The second factor is the additional proofreading efficiency *f*_*pro*_ contributed by the exonuclease domain, which is defined by the ratio of the elongation probability of the terminal R (*P*_*el*_,*R*) to that of the terminal W (*P*_*el*_,*W*). Here the elongation probability is defined as *P*_*el*_ = *k*_*el*_/(*k*_*el*_ + *k*_*ex*_), *k*_*el*_ is the rate of elongation to the next site, and *k*_*ex*_ is the rate of excising the primer terminal nucleotide (*e.g.* Eq.(A1-A6) in Ref.[13]). *P*_*el*_,*R* is usually assumed close to 100%, so *f*_*pro*_ equals approximately to 1 + *k*_*ex*_,*W* /*k*_*el*_,*W*. Although these expressions seem reasonable, there are some problems that were not clarified. First, the mathematical definition of *f*_*pro*_ is subjective though intuitive, so a rigorous theoretical foundation is needed. Second, the apparent rate parameters *k*_*el*_ and *k*_*ex*_ are not well defined since both the elongation and the excision are multi-step processes. *k*_*el*_ and *k*_*ex*_ are unknown functions of the involved rate constants but there is not a unique way to define them. They could be theoretically defined under steady-state assumptions (Eq.(6) in Ref.[14]) or operationally defined by experiment assays (*e.g.* steady-state assays[15, 16] or transient-state assays [17]), but different ways often lead to different estimates of *f*_*pro*_ (see Sec.III). Additionally, *k*_*el*_ should be properly understood as the ultimate elongation rate in the sense that the elongated terminal (the added nucleotide) is no longer excised. This condition is not met if the *exo*^+^-DNAP can proofread the buried mismatches (*e.g.* the penultimate or antepenultimate mismatches, etc.). In these cases, *k*_*el*_ is affected not only by the next template site but also by further sites. Such far-neighbor effects were not seriously considered in previous studies. For these reasons, the widely cited initial discrimination *f*_*ini*_ ≈ 10^4~5^ and the proofreading efficiency *f*_*pro*_ ≈ 10^2~3^ [18] are questionable.

Recently two equivalent rigorous theories were proposed to investigate the true fidelity of either *exo*^−^-DNAP or *exo*^+^-DNAP, *i.e.* the iterated function systems by P.Gaspard [19] and the first-passage (FP) method by us [20]. In particular, we can numerically rigorously compute the true fidelity (ℱ) of *exo*^+^-DNAP by the FP method and can also derive the approximate analytical expressions (*F*) of the true fidelity. With these firmly established results, we can address all the above questions systematically. In the following sections, we will first give a brief review of the FP method and the major conclusions already obtained for the minimal kinetic model of DNA replication. Then we will generalize these conclusions to more realistic kinetic models for *exo*^−^-DNAP and *exo*^+^-DNAP, and carefully examine the relations between *f*_*s·s*_, *f*_*t·s*_ and ℱ.

## II. METHODS

### A. Basics of the FP method

The first-passage (FP) method was proposed to study the replication of the entire template by *exo*^+^-DNAP [20].

Here the minimal reaction scheme Fig.1 is taken as an example to illustrate the basic logic of this method. 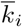 is the rate of incorporating dNTP to the primer at the template site *i* − 1 (the dNTP-concentration dependence of 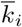 is not explicitly shown here), 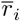 is the rate of excising the primer terminal at the template site *i*. Intuitively, 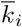 and 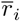 depend on the identity (A, G, T or C) and the state (matched or mismatched) not only of the base pair at site *i* but also of the one or more preceding base pairs. If there is only the nearest-neighbor (first-order) effect, 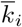 and 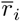 can be written as 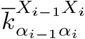and 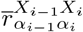, *X*_*i*–1_ (or *X*_*i*_) represents the nucleotide at site *i* – 1 (or *i*) on the template, *α*_*i*-1_ represents the nucleotide at site *i* – 1 on the primer, *α*_*i*_ represents the next nucleotide to be incorporated to the primer terminal at site *i* (for 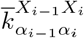) or the terminal nucleotide of the primer at site *i* to be excised (for 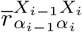). *X* and *α* can be any of the four types of nucleotides. Similarly, there are 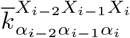 etc. for the second-order neighbor effects, and so on for farther-neighbor (higher-order) effects.

**FIG. 1:**
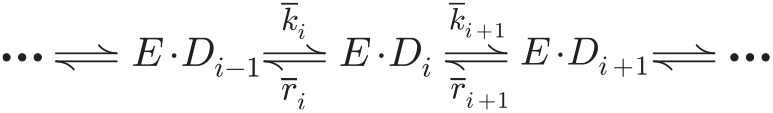
The minimal reaction scheme of DNA replication. *E*: the enzyme DNAP. *D*_*i*_: the primer-template duplex with primer terminal at the template site *i*.

To calculate the true fidelity ℱ, we consider the replication of any given template *X*_1_*X*_2_…*X*_*L*_ of length *L*. The primer grows from the starting unit *α*_1_ and reaches the last site *L* to generate the final product of the sequence *α*_1_*α*_2_…*α*_*L*_. Except the first unit *α*_1_ and the last unit *α*_*L*_, any incorporated dNTP can be excised during this replication. In other words, this is a first-passage process from the reflecting boundary at the first site to the absorbing boundary at the last site. Since there are four types of dNTP competing for the incorporation reaction at each site, 4^*L*^ kinds of final products with different sequences should be generated. By solving the corresponding equations numerically, one can obtain the probability distribution of the final product 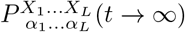 from which the true fidelity ℱ can be precisely counted. One can also derive approximate analytical expressions for the true fidelity under some conditions of the kinetic parameters. A brief introduction of these calculations is given in Appendix A (see also Supplemental Material (SM) Sec.I A.). It is worth noting that the FP method does not need any extra assumptions like the steady-state or the quasi-equilibrium assumptions. Below we list the major results in terms of 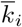 and 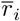.

### B. The fidelity calculated by the FP method

For *exo*^+^-DNAP which has first-order neighbor effects, we have derived the approximate analytical expression of the overall fidelity at site *i* [20],

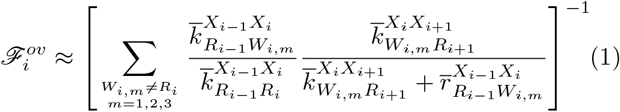

*R* represents the matched nucleotide, and *W*_*i,m*_ represents one of the three types of mismatched nucleotides. This formula provides a quite good approximation of ℱ: the relative deviation from the precise numerical result is less than 10% in a wide range of the kinetic parameters (in this paper, the relative deviation between any two quantities *a* and *b* is defined as |a–b|/min(*a, b*), min(*a, b*) means the smaller one of *a* and *b*. Details can be found in SM Sec.I A). This formula shows that the true fidelity at site *i* is overwhelmingly determined by the nearest neighbor sites. For simplicity, we omit all the superscripts below unless it causes misunderstanding, and also use *W*_*i*_ to represent any type of mismatch. Each term in the sum represents the error frequency of a particular type of mismatch, whose reciprocal (denoted as *F*_*i*_) corresponds to the mismatch-specific fidelity discussed in the conventional steady-state assay or transient-state assay.Here the mismatch-specific fidelity is denoted as ℱ_*i*_ ≡ ℱ_*ini,i*_ · ℱ_*pro,i*_, in which ℱ_*ini,i*_ is the initial discrimination and ℱ_*pro,i*_ is the proofreading efficiency. Similarly, we denote 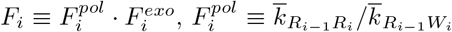, 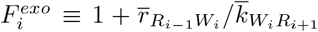. According to Eq.(1), we have

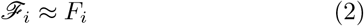

Since ℱ_*ini,i*_ is actually the fidelity for *exo*^−^-DNAP, we can set 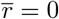 in Eq.(1) to obtain

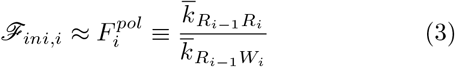

Hence, ℱ_*pro,i*_ ≡ ℱ_*i*_/ℱ_*ini,i*_ can be estimated by 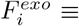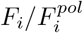, as below

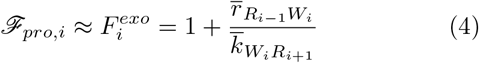

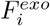 is similar to *f*_*pro*_ defined in Sec.I, if 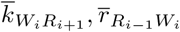 are regarded as *k*_*el,W*_, *k*_*ex,W*_ respectively. In these equations, ≈ means that the relative deviations of *F*_*i*_, 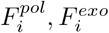 from ℱ_*i*_, ℱ_*ini,i*_, ℱ_*pro,i*_ are less than 10% (see details in SM Sec.I A).
For *exo*^+^-DNAP which has second-order neighbor effects, we have also derived the approximate analytical expressions for the overall fidelity at site *i* [20],

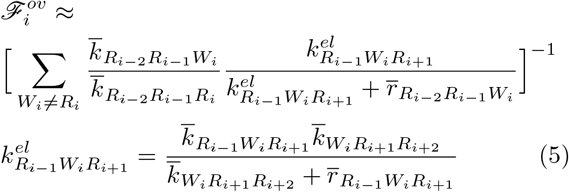

Each term in the sum represents the mismatch-specific error frequency at site *i*. Its reciprocal defines the mismatch-specific fidelity which again consists of the initial discrimination and the proofreading efficiency, but the latter differs significantly from *f*_*pro*_ defined in Sec.I, since the effective elongation rate is not 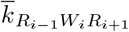 but instead 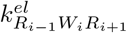 which includes the next-nearest neighbor effects. The same logic can be readily generalized to higher-order neighbor effects where the proof-reading efficiency will be more complicated [20, 21].

In real DNA replication, either the dNTP incorporation or the dNMP excision is a multi-step process. By using the FP method, the complex reaction scheme can be reduced (or mapped) to the minimal scheme Fig.1, and the fidelity can still be calculated by Eq.(2) or Eq.(5), with only one modification: 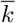 and 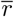 are now the effective incorporation rates and the effective excision rates respectively which are functions of the involved rate constants. In the following sections, we will derive these effective rates by FP method for different multi-step reaction models, and compare them with the effective rates given by the steady-state assays or the transient-state assays. For simplicity, we only discuss the first-order neighbor effects of DNAP in details, since almost all the existing literature focused on nearest-neighbor effect-Higher-order neighbor effects will also be mentioned in later sections.

Below shows a simple example of the effective rates calculated by the FP method for the single-nucleotide incorporation reaction in the direct competition assays.

Fig.2 shows a three-step kinetic model of the competitive incorporation of a single dRTP or dWTP to site *i* + 1, catalyzed by *exo*^−^-DNAP. The direct competition assay estimates the true fidelity by the ratio of the final product [*D*_*i*_*R*] to [*D*_*i*_*W*], which can be interpreted by the FP method as below

**FIG. 2:**
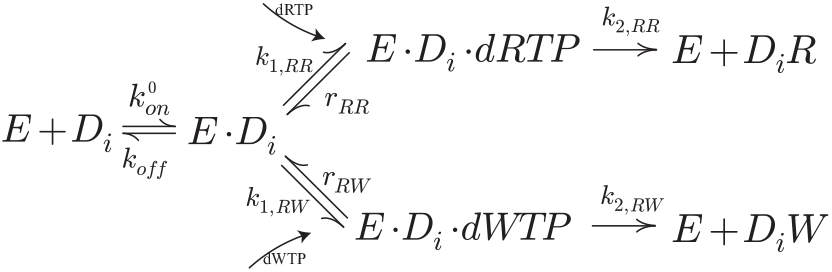
The three-step reaction scheme of the competitive incorporation of dRTP and dWTP. E: *exo*^−^-DNAP. *D*_*i*_: the primer-template duplex with the matched(R) terminal at site *i*. For brevity, the subscript *i* in each rate constant is omitted.

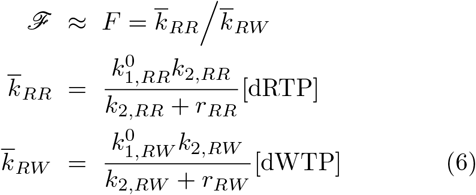

*F* is exactly the initial discrimination defined by Eq.(3) with the two effective incorporation rates 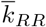 and 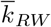. These equations predict correctly the dNTP concentration-dependence of the fidelity which is consistent of the experimental observations in the direct competition assays [6]. Detailed discussion can be found in SM Sec.I B.

## III. RESULTS AND DISCUSSION

As mentioned above, the total fidelity of *exo*^+^-DNAP is assumed to consist of the initial discrimination *f*_*ini*_ and the proofreading efficiency *f*_*pro*_. In this section we present a detailed analysis to show the availability and limitations of the conventional kinetic assays to characterize *f*_*ini*_ and *f*_*pro*_. The reaction scheme under discussion is shown in Fig.3.

**FIG. 3:**
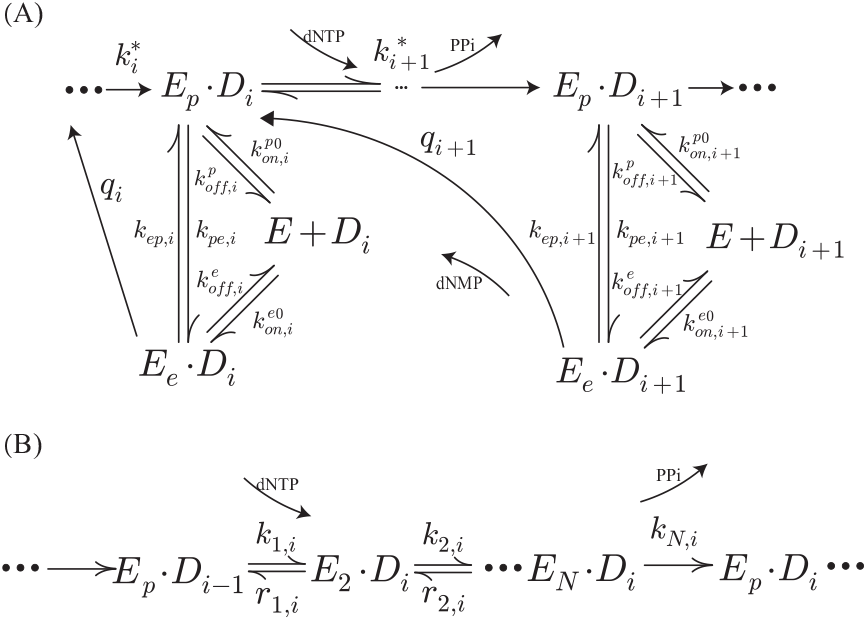
(A): The multi-step reaction scheme of *exo*^+^-DNAP including the multi-step incorporation process (indicated by *k*^*^ and details are shown in (B)), the intramolecular/intermolecular transfer (*i.e.* without/with DNAP dissociation) of the primer terminal between *Pol* and *Exo* and the excision of the primer terminal nucleotide. (B): The multi-step incorporation scheme. The enzyme-substrate complex (*E·D*) goes through *N* states (indicated by subscripts 1, …, *N*) to successfully incorporate a single dNTP (indicated by subscript *i*). To simplify the notation, the superscripts indicating the template nucleotide *X*_*i*_ and the subscripts indicating the primer nucleotide *α*_*i*_ are omitted. This rule also applies to other figures in this paper, unless otherwise specified.

In this reaction scheme, the primer terminal can transfer between *Pol* and *Exo* in two different ways, *i.e.* the intramolecular transfer without DNAP dissociation (the transfer rates are denoted as *k*_*pe*_ and *k*_*ep*_), and the inter-molecular transfer in which DNAP can dissociate from and rebind to either *Pol* or *Exo* (the rates are denoted as *k*_*on*_ and *k*_*off*_). These two modes have been revealed by single-turnover experiments[17] and directly observed by smFRET[22]. Here the quasi-first order rate *k*_*on*_ is proportional to the concentration of DNAP or DNA, *i.e.* 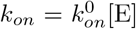 or 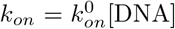 (see later sections). *k*^*^ is the effective incorporation rate, as explained below.

### A. The FP method

Applying the FP method to complex reaction schemes like Fig.3(A), one can always reduce them to the simplified version Fig.1 with uniquely determined effective rates. For instance, for the multi-step incorporation schemes in Fig.3(B), the effective incorporation rate *k*^*^ is given by (see details in Appendix B or SM Sec.I C)

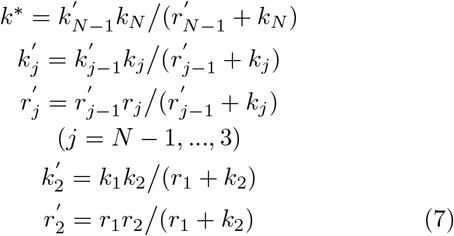

Here 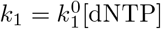, so *k*^*^ = *k*^*0^[dNTP].

For the complete scheme Fig.3(A), the effective rates are

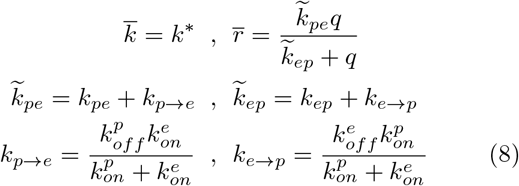

Details can be found in SM Sec.I D. These rates can be written more explicitly such as 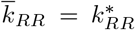, if the states of the base pairs at site *i*, *i* – 1, *etc*. are explicitly indicated. All the rates in the same equation have the same state-subscript. *k*_*p*→*e*_ and *k*_*e*→*p*_ define the effective intermolecular transfer rates between *Pol* and *Exo*. So 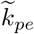 and 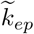 represent the total transfer rates *via* both the intramolecular and the intermolecular ways. Here 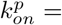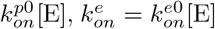 so *k*_*p*→*e*_, *k*_*e*→*p*_ do not depend on [E]. With these effective rates, the initial discrimination and the proofreading efficiency can be estimated by 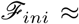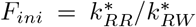 and 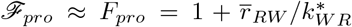 respectively. Here we use the symbols *F*_*ini*_ and *F*_*pro*_ to replace *F*^*pol*^ and *F*^*exo*^ (see Sec.II) which are defined only for the minimal scheme. We keep these notations for any multi-step scheme in the following sections.

### B. The steady-state assay

#### 1. The initial discrimination

The steady-state assay is a standard method to analyze the catalytical capability of enzymes in biochemistry. The steady-state condition in experiment is usually established by two requirements, *i.e.* the substrate is in large excess to the enzyme, and the enzyme can dissociate quickly from the product once a single turnover is finished to resume its catalysis. The last dissociation step is reasonably assumed irreversible, since the enzyme will much unlikely rebind to the same substrate molecule after dissociation because the substrate is in large excess to the enzyme. Under such conditions, the amount of any intermediate product can be regarded approximately as constant (*i.e.* in steady state) in the initial stage of the reaction and thus the amount of the final product increases linearly with time. This initial growth velocity is defined as the steady-state turnover rate which is often used to characterize the catalytical capability of the enzyme.

It should be noted that, however, the steady-state assay is only valid for enzymes with single catalytical activity such as *exo*^−^-DNAP. So one should properly modify the *exo*^+^-DNAP to obtain mutant *exo*^−^-DNAP (*e.g.* by deactivating the exonuclease activity or even eliminate the exonuclease domain by genetic mutations, without changing the polymerase activity of the DNAP), and then employ the steady-state assay to measure the initial discrimination of the DNAP. By measuring the initial velocity of the final product generation (*i.e.* single dNTP incorporation) under the steady-state condition, one can calculate the normalized velocity per enzyme which is in general given by the Michaelis-Menten equation

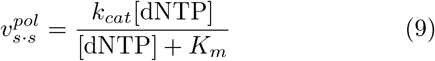

Here *k*_*cat*_ is the maximal steady-state turnover rate of dNTP incorporation and *K*_*m*_ is the Michaelis constant. The superscript *pol* indicates the polymerase activity, the subscript *s · s* indicates the steady state. Fitting the experimental data by this equation, one can get the specificity constant *k*_*cat*_/*K*_*m*_ either for dRT-P incorporation or dWTP incorporation, from which the initial discrimination was defined as *f*_*s·s,ini*_ = (*k*_*cat*_/*K*_*m*_)_*R*_[dRTP]/(*k*_*cat*_/*K*_*m*_)_*W*_[dWTP].

However, there is an apparent difference between the above-defined fidelity and the true fidelity: the former is measured under steady-state conditions, whereas the latter is defined without pre-assumptions like the steady-state approximation. So what is the relation between them?

To understand the exact meaning of *k*_*cat*_/*K*_*m*_, we have to consider the multi-step reaction scheme of dNTP incorporation (Fig.4) which explicitly includes the DNAP binding to DNA and dissociation from DNA. In general, we consider cases where the substrate DNA can bind either to the polymerase domain or to the deficient exonuclease domain (if the domain is only mutated but not eliminated). For the mutant *exo*^−^-DNAP, the transfer rate 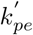 and 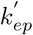 are different from *k*_*pe*_ and *k*_*ep*_ of the wild type *exo*^+^-DNAP. Under the steady-state condition, it can be easily shown

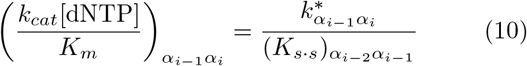

**FIG. 4:**
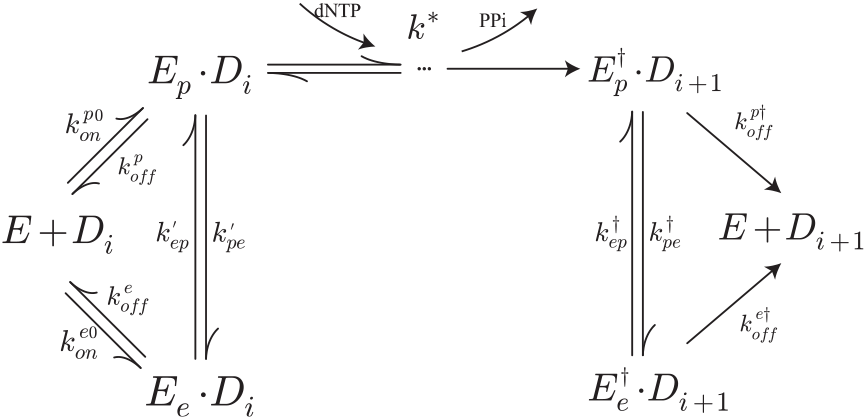
The reaction scheme for the steady-state assay to measure the specificity constant of the nucleotide incorporation reaction of DNAP with deficient exonuclease domain. The omitted multi-step incorporation process is shown in Fig.3(B).

Here *k*^*^ is defined in Eq.(7), *α* = *R, W*. 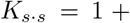 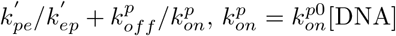

Theoretically, the real reaction scheme may be more complicated than Fig.4, *i.e.* there could be more pathways or more steps in DNAP binding/dissociation or DNA transfer. No matter how complicated the scheme is, it can be proven that the form of Eq.(10) is universal: *k*^*^ is exactly the effective incorporation rate defined by Eq.(7), and *K*_*s·s*_ is a simple function of the equilibrium constants of the steps just before dNTP binding(see details in SM Sec.II A,B).

Since *K*_*s·s*_ is independent on the incoming dNTP(*α*_*i*_), the fidelity can be given generally by

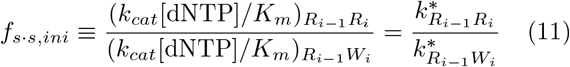

which is equal to *F*_*ini*_. Since *F*_*ini*_ is a good approximation to ℱ_*ini*_ with less than 10% deviation, the steady-state assay provides a quite good measure of the initial discrimination.

Below we give an example of using the measured specificity constants to estimate the site-specificity fidelity. In the steady-state assay of KF^−^ [6], *k*_*cat*_ and *K*_*m*_ of the incorporation of 4 types of dNTP onto the template <u>T</u> (the underline indicates the target site under investigation) has been measured separately, as listed in Table I. The initial discrimination *f*_*s·s,ini*_ of each mismatch can then be calculated by Eq.(11). The result is shown in the last column, indicating the significant mismatch-specific variations of the *f*_*s·s,ini*_. One can also calculate the site-specific initial discrimination, ≈ 7.6 × 10^3^, simply by using Eq.(1).

**Table I:**
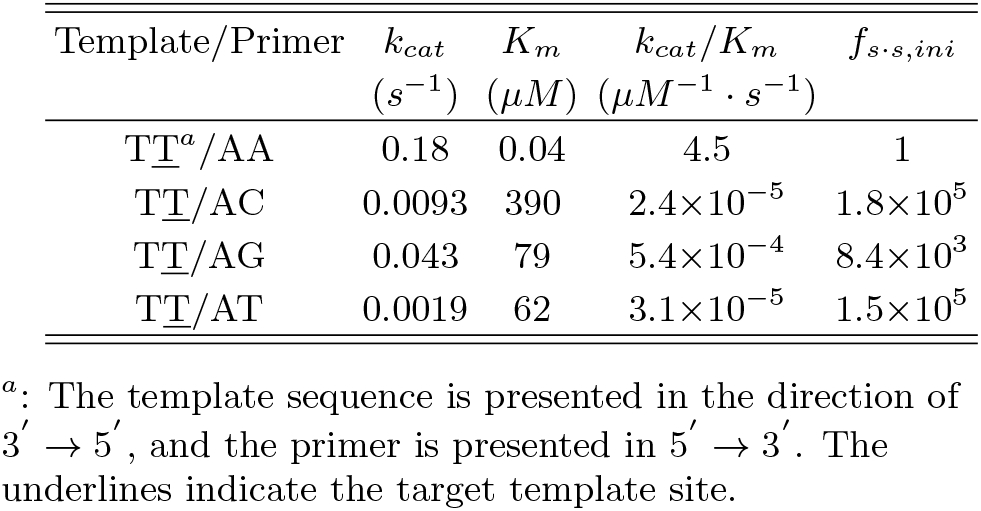
The steady-state assay of KF^−^[6]

#### 2. The proofreading efficiency

There were also some works using the steady-state assay to define the effective elongation rate *k*_*el,W*_ and the effective excision rate *k*_*ex,W*_ in order to characterize the proofreading efficiency *f*_*s.s,pro*_ = 1+*k*_*ex,W*_/*k*_*el,W*_ of *exo*^+^-DNAP. If *k*_*ex,W*_ equals to 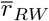 and *k*_*el,W*_ equals to 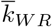, we then have 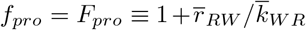. Unfortunately, however, neither equation holds in general.

For instance, some works used the turnover velocity 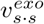 [17, 23], the specificity constant[24, 25] or the maximal turn-over rate *k*_*cat*_ [15] as *k*_*el,W*_. As shown by Eq.(10), however, 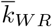 is not equal to any of the three quantities. So, the steady-state assay fails to measure 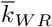, unless *K*_*s·s*_ ≈ 1. For the reaction scheme Fig.4, this condition may be met, since the mutant *exo*^−^-DNAP has no exonuclease domain 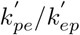 is absent) or it binds the DNA preferentially at the polymerase domain 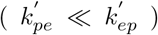 and the DNA concentration in the experimental assay can be set large enough to ensure 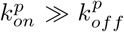. In such cases, the specificity constant, but not *k*_*cat*_ or 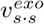, can be regarded as 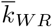.

Of course *K*_*s·s*_ ≈ 1 may not hold if one considers reaction schemes more complicated than Fig.4. For instance, the primer terminal transfer may be a multi-step process rather than the one-step process shown in Fig.4. In this paper we do not discuss such schemes, since so far there are no experimental supporting evidences (except that there could be an additional DNAP translocation step before dNTP binding, as discussed in later sections). So we assume that 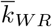 can be reasonably measured by the specificity constant for cases discussed in this paper.

The steady-state assay was also employed to study the excision reaction of DNAP (Fig.5). There was some studies which measured the initial velocity 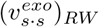 and interpreted it as *k*_*ex,W*_ [15]. Whereas 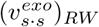 is determined by all the rate constants in Fig.5, some rate constants like 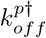 and 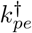 are absent from the effective excision rate 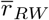. So in principle, 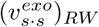 is not equal to 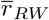.

**FIG. 5:**
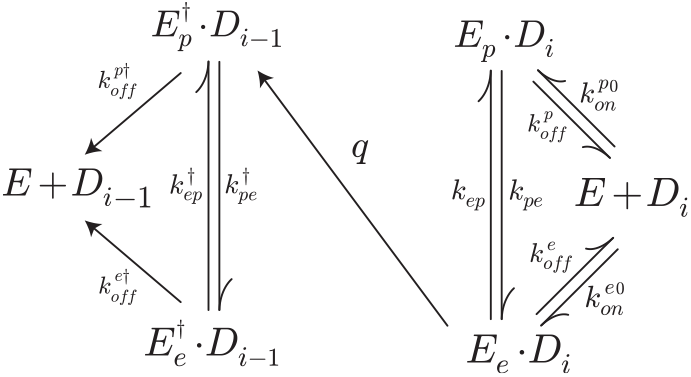
The reaction scheme for the steady-state assay to measure the effective excision rate of *exo*^+^-DNAP.

In summary, since 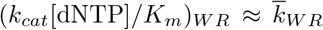 but 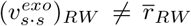, *f*_*s·s,pro*_ may differ significantly from *F*_*pro*_. A numerical example is shown in Fig.6, where 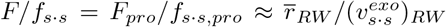 (here we have assumed *k*_*ex,W*_ ≫ *k*_*el,W*_ and 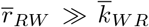 for efficient proofreading).

**FIG. 6:**
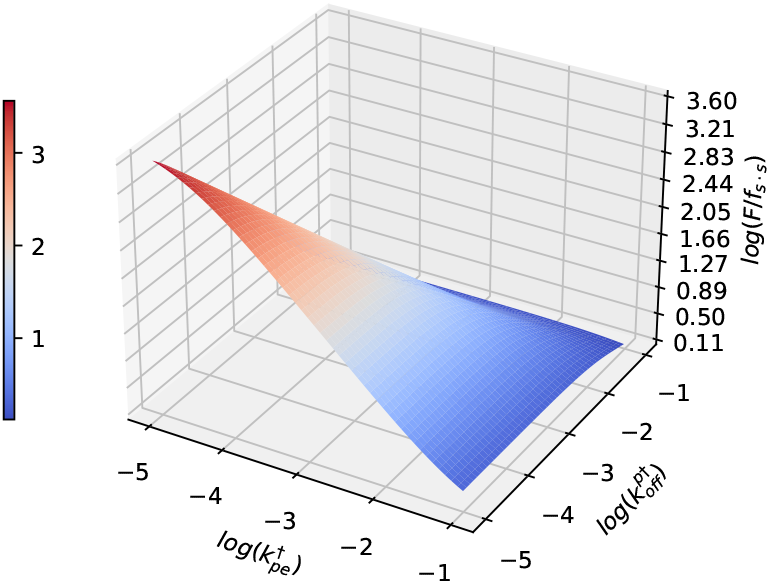
The ratio *F*/*f*_*s·s*_ may become larger than 10 when 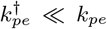 and 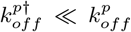. The kinetic rates are set as *q* = 1, 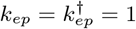, *k*_*pe*_ = 0.1, 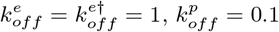, 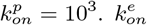 is calculated according to the thermodynamic constraint 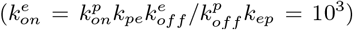. The binding rates are large enough, *i.e.* 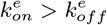 and 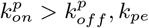.

Because the fidelity is a dimensionless number, the absolute values of the kinetic rates do not matter for our calculation. So we set *q* = 1 and other kinetic rates in unit of *q* with the magnitudes inspired by the measured values for T7 DNAP [17] (Table II), *e.g. k_ep_* and 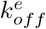 are set equal to *q*, and *k*_*pe*_ and 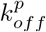 are smaller than *q* which are set to be 0.1. 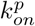 and 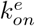 are determined by DNA concentration which is often taken as 1*μM* in experiments(*e.g.* [15]). So these two values can be very large and we set them to be 10^3^ in the calculations, which ensures 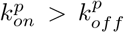 and 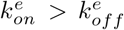. Larger values do not change the results. The dissociation rate from *Exo* are assumed irrelevant to the identity of the terminal, so 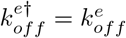.

**Table II:** Kinetic rates of the excision of a mismatched T of A<u>G</u>/TT of T7 DNAP[17]

| kinetic parameter | rate |
| --- | --- |
| $q$ | $896s^{-1}$ |
| $k_{ep}$ | $714s^{-1}$ |
| $k_{pe}$ | $2.3s^{-1}$ |
| $k_{off}^p$ | $0.4s^{-1}$ |
| $k_{off}^e$ | $1022s^{-1}$ |
| $k_{on}^{p0}$ | $4 \times 10^7 M^{-1}s^{-1}$ |
| $k_{on}^{e0}$ | $(4 - 6) \times 10^8 M^{-1}s^{-1}$ |

Fig.6 shows that the ratio *F*/*f*_*s·s*_ can even become larger than 10 when 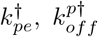 are much smaller than *k*_*pe*_, 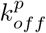. This may unfortunately be true, since 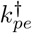 is the transfer rate of the newly formed matched terminal (after excision), which should be much smaller than the corresponding rate *k*_*pe*_ of the original mismatched terminal. 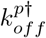 is the dissociation rate of DNAP from the matched terminal, which should be much smaller than that from the mismatched terminal 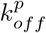. So the steady-state assay are invalid to measure the proofreading efficiency of *exo*^+^-DNAP. In the experiment to estimate ℱ_*pro*_ for ap-polymerse[15], the authors wrongly interpreted *k*_*cat*_ and 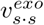 as 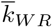 and 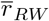 respectively, and gave that 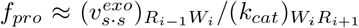. This measure is even totally irrelevant to ℱ_*pro*_.

One can also change the concentration of the substrate DNA to obtain the specificity constant of the excision reaction in experiments[16], as can be shown theoretically

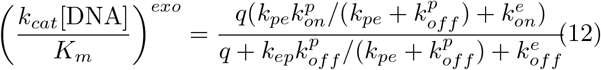

This expression includes explicitly the DNA concentration, whereas the effective rate 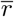 is independent of DNA concentration. So, it can not be used to estimate the effective excision rate. Details can be found in SM Sec.II C.

### C. The transient-state assay

#### 1. The initial discrimination

Similar to the steady-state assay, the transient-state assay can also be employed to study the polymerase and exonuclease of DNAP separately. The transient-state assay often refers to two different methods, the pre-steady-state assay or the single-turnover assay. Since the theoretical foundations of these two methods are the same, we only discuss the latter below for simplicity.

In single-turnover assays, the enzyme is in large excess to the substrate, and so the dissociation of the enzyme from the product is neglected. The time course of the product accumulation or the substrate consumption is monitored. The data is then fitted by exponential functions (single-exponential or multi-exponential) to give one or more exponents (*i.e.* the characteristic rates). In the initial discrimination assay, these rates are complex functions of all the involved rate constants and dNTP concentration, which in principle can be analytically derived for any given kinetic model. For instance, for the commonly-used two-step model including only substrate binding and the subsequent irreversible chemical step, one can directly solve the kinetic equations to get two rates. On the other hand, the time course of the product accumulation can often be well fitted by single-exponential function at any given dNTP concentration, which implies that the two rates differ by more than one order of magnitude and only the smaller one is recorded in experiments. It was proved by K. Johnson that the smaller one obeys approximately the Michaelis-Menten-like equations [26](see also SM Sec.III A).

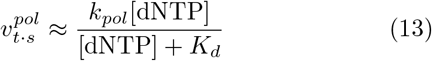

The subscript *t · s* indicates the transient state. Similar to the steady-state assays, *k*_*pol*_/*K*_*d*_ is regarded as the specificity constant and thus the initial discrimination is defined as *f*_*t·s,ini*_ = (*k*_*pol*_/*K*_*d*_)_*R*_[dRTP]/(*k*_*pol*_/*K*_*d*_)_*W*_ [dWTP].

To get a better understanding of *k*_*pol*_/*K*_*d*_, we consider the much more realistic multi-step scheme in Fig.7, in which the DNA can bind to either the polymerase domain or the deficient exonuclease domain and the dNTP incorporation is a multi-step process. One can prove that Eq.(13) still holds for such schemes and the specificity constant can be approximated as

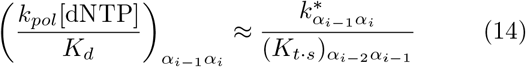

**FIG. 7:**
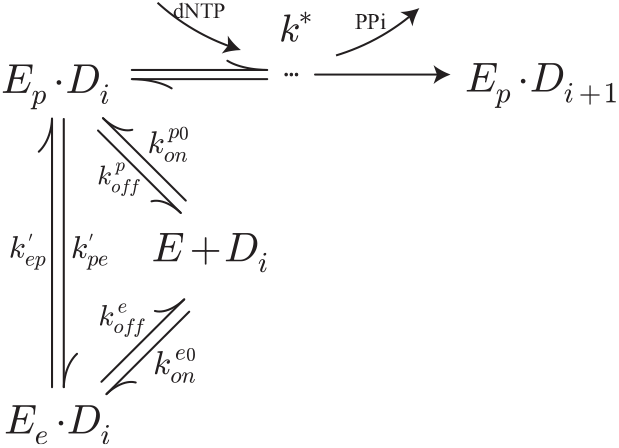
The reaction scheme for the transient-state assay to measure the specificity constant of the nucleotide incorporation reaction of DNAP with deficient exonuclease domain. The omitted multi-step incorporation process is shown in Fig.3.(B).

Eq.(14) provides a rough estimate of the real specificity constant with less than 200% relative deviation. The rigorous analysis is too lengthy to be presented here (for details see SM Sec.III B). Here *α* = *R, W*. 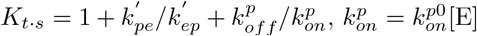. Hence, the relative deviation of *f*_*t·s,ini*_ from 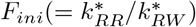 is less than 200%. Since *F*_*ini*_ ≈ ℱ_*ini*_ (with less than 10% deviation), the transient-state assay can give a rough estimate on ℱ_*ini*_. In practice, both the steady-state assay and the transient-state assay are always used (say, Ref.[6]) to guarantee that the specificity constants obtained by both methods agree with one another (no order of magnitude difference) in order to make a good estimate on *k*^*^ and thus the true initial discrimination. Additionally, the specificity constant, but not *k*_*pol*_, can be used to estimate 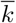 when *K*_*t·s*_ ≈ 1.

#### 2. The proofreading efficiency

The transient-state assay of the exonuclease activity is often done under single-turnover conditions. The time course of product accumulation or substrate consumption is fitted by a single exponential or a double exponential to give one or two characteristic rates[17, 24, 25]. In the following, we show that the smallest one of the fitted exponents may probably be equal to 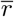 under some conditions.

The simplest model for the transient-state assay of the excision reaction is depicted in Fig.8. By solving the corresponding kinetic equations, one can get three characteristic rates and the smallest one is given by

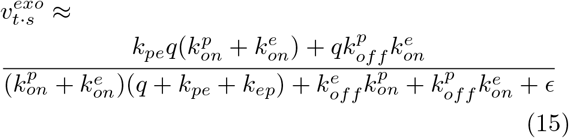

**FIG. 8:**
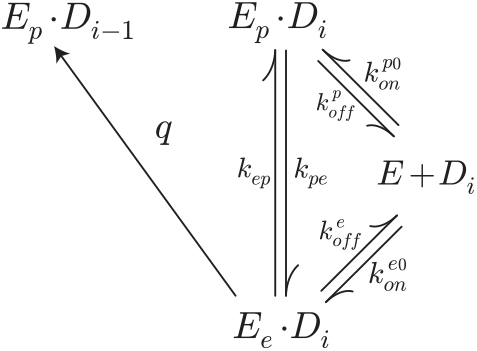
The reaction scheme for the transient-state assay to measure the effective excision rate of *exo*^+^-DNAP.

Eq.(15) gives a rough estimate on the precise 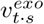 with less than than 200% deviation (details can be found in SM Sec.III C.). Here 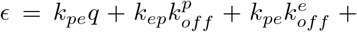 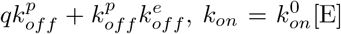. The DNAP concentration in the experiments can be set large enough, which ensures 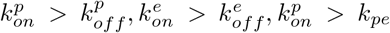 and ϵ ≈ 0 (compared to other terms in the denominator), Eq.(15) can be simplified as

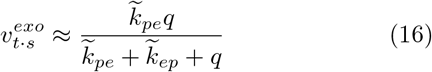

If 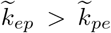 (when DNA binds preferentially to the polymerase domain) or 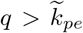 (when the excision is a very fast process), one can get 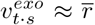, with 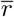 defined by Eq.(8). So, if the real excision reaction follows the simplest scheme in Fig.8, 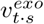 may be interpreted as 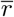. However, if *q*, 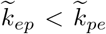, there could be large difference between 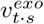and 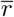. Fig.9 gives an example showing difference tihnea wide range of some key parameters.

**FIG. 9:**
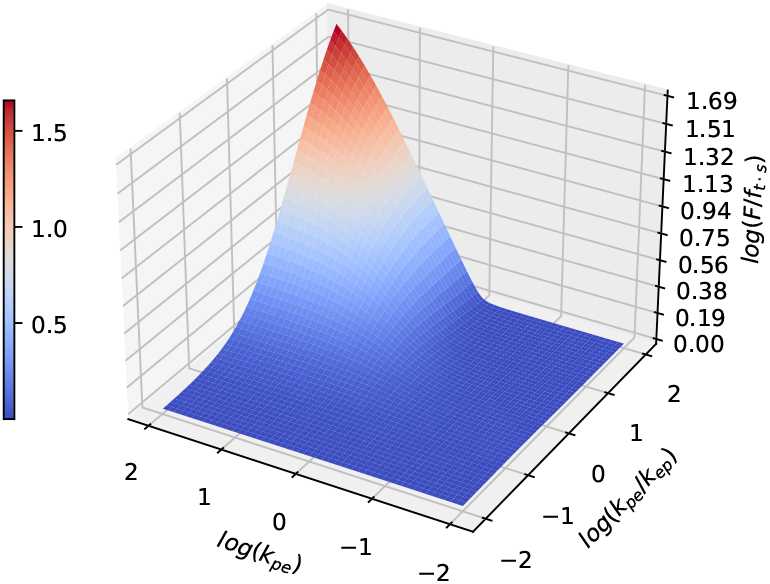
The ratio *F*/*f*_*t* · *s*_ may become lager than 10 when q, 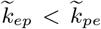. The kinetic rates are set as *q* = 1, 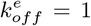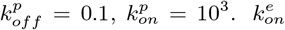 is calculated by the thermodynamic constraint 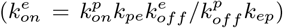. The binding rates are always large enough when adjusting *k*_*pe*_ and *k*_*ep*_, *i.e.* 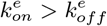 and 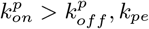, *k*_*pe*_.

In Fig.9 the proofreading efficiency is defined as *f*_*t*·*s*_ = 1 + *k*_*ex,W*_ / *k*_*el,W*_, *k*_*ex,W*_ is taken as 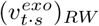 and *k*_*el,W*_ is taken as (*k*_*pol*_[dNTP]/*K*_*d*_)_*W R*_. Since 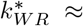 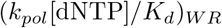, we have 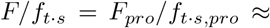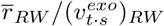. In this example, we show the precise value of 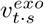 by numerically solving the original kinetic equations, rather than using the approximate expression Eq.(15). The kinetic rates are set as explained in Sec.III B2. The typical free energy difference between DNA binding to *Pol* and *Exo* may be only a few *k*_*B*_*T*, so we set 10^−2^ < *k*_*pe*_/*k*_*ep*_ < 10^2^. As shown, the ratio *F*/*f*_*t*·*s*_ may become even larger than 10 when *q*, 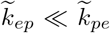 (the red area). So the transient-state assay can be used to roughly measure the proofreading efficiency only when 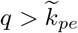 or 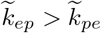 (the blue region in Fig.9) .

In Table II we lists all the kinetic rates of the excision reaction of T7 DNAP determined by the experiment [17], which shows *q*, 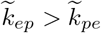, so we can use 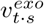 to estimate 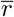 and calculate the *F*_*pro*_ by Eq.(4). In the original paper [17], however, the authors defined the proofreading efficiency as 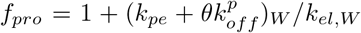, with the ambiguous quantity *θ* which was supposed between 0 and 1 (depending on the fate of the DNA after dissociation), and regarded *k*_*el,W*_ as 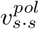 (Eq.(9)). This is essentially different from Eq.(4). Moreover, the authors ignored the template sequence-dependence of the proofreading efficiency in their calculations. In fact, Table II lists the rates of excising a mismatched T of A<u>G</u>/TT, the corresponding elongation rate (*k*_*el,W*_) should be the rate of incorporating the matched dNTP over the mismatched terminal T, *i.e.* 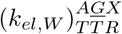 (*X* can be one of *A, T, G, C*. *R* means the dNTP matched to *X*), according to Eq.(4).

But the authors used 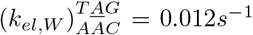 (calculated by Eq.(9) with the measured *k*_*cat*_ = 0.025*s*^−1^, *K*_*m*_ = 87*μ*M, [dNTP]=100*μ*M in Ref.[23]) to estimate f_pro_ and finally gave *f*_*pro*_ = 1 + (2.3 + 0.5 × 0.4)/0.012 ≈ 210 (*θ* = 0.5). So this estimate is highly questionable. Even if the template sequence-dependence of k*_el_*, *W* can be ignored, one should estimate *F*_*pro*_ by Eq.(4). The elongation rate *k*_*el,W*_ (or *k*_*W R*_) is approximately determined as *k*_*cat*_[dNTP]/*K*_*m*_ = 0.03*s*^−1^. The effective excision rate is determined by Eq.(8), 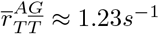 So we get a rough estimate of the proofreading efficiency for this template sequence, *ℱ*_*pro,i*_ ≈ *F*_*pro*_ ≈ 1 + 1.23/0.03 ≈ 40 which is much smaller than that given by the authors.

There is another example possibly showing the second-order neighbor effects in proofreading of Human Mitochondrial DNAP. The excision velocities and the elongation rates were measured for a particular template sequence GT<u>T</u>GG/CAT by the transient-state assays[24, 25], as listed in Table III. Since there is no additional information on *q,* 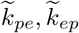, we can not use these values to estimate the proofreading efficiency. Here we just assume that the excision velocities can be taken as the effective excision rates to make a rough estimate. *F*_*pro*_ of a certain type of mismatch (Eq.(5)) can be rewritten as

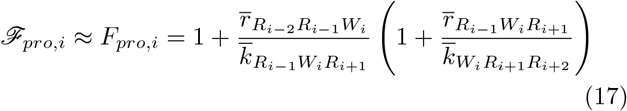

**Table III:** Kinetic rates of human mitochondrial DNAP[24, 25]

| the effective rate | the experiment measure | the measured rate |
| --- | --- | --- |
| $\bar{r}_{CAT}^{GTT}{}^a$ | $v_{t.s}^{exo}$ | $0.4s^{-1}$ |
| $\bar{k}_{ATC}^{TTG}$ | $k_{pol}[dNTP]/K_d{}^b$ | $0.1s^{-1}$ |
| $\bar{r}_{ATC}^{TTG}$ | $v_{t.s}^{exo}{}^c$ | $3.0s^{-1}$ |
| $\bar{r}_{ATC}^{TTG}$ | $\hat{v}_{t.s}^{exo}{}^d$ | $21.5s^{-1}$ |
| $\bar{k}_{TCC}^{TGG}$ | $k_{pol}[dNTP]/K_d$ | $2.7s^{-1}$ |

<sup>a</sup>: The template sequence around the target site $\underline{T}$ is
presented in the direction of $3' \rightarrow 5'$ : GTTGG
<sup>b</sup>: $[dNTP] = 100\mu M$
<sup>c</sup>: without dNTP in the solution
<sup>d</sup>: in the presence of dNTP

With the kinetic rates in Table III, we get

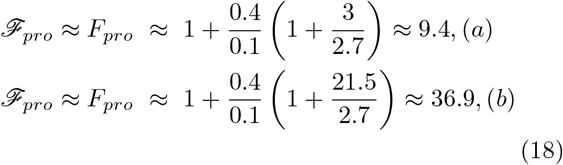

These two values correspond to two different experimental conditions: (a) without dNTP in the solution; (b) in the presence of dNTP. In the latter case, the second-order proofreading (21.5/2.7) contributes to *ℱ*_*pro*_ even more than the first-order proofreading (0.4/0.1).

It is worth emphasizing that the interpretation of 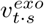 is model-dependent. Theoretically, the reaction scheme could be more complicated than the simplest model in Fig.8, *e.g.* there may be multiple substeps in the intramolecular transfer process since the two domains are far apart (2-4 nm[18]). For any complex scheme, one can analytically calculate 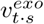 and 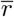. These two functions always differ greatly (examples can be found in SM Sec.III D). So the single-turnover assay *per se* is not a generally reliable method to measure the effective excision rate.

In total, neither the steady-state assay nor the transient-state assay can reliably measure the effective elongation rates and the effective excision rates, so it’s invalid in principle to use these assays to estimate the proofreading efficiency of DNAP, unless one can do more detailed study to confirm that the required conditions on the key rate parameters are actually met.

### D. More realistic models including DNAP translocation

So far we have not considered the important step, DNAP translocation, in the above kinetic models.DNAP should translocate forward along the template to the next site after dNTP incorporation, which empties the active pocket to accept the next dNTP for incorporation. Goodman *et al.* had discussed the possible effect of such a translocation on the transient-state gel assay very early[27], and recently DNAP translocation has been directly observed for phi29 DNAP by using nanopore techniques[28–31] or optical tweezers[32]. However, so far there is no any theory or experiment to seriously study the effect of translocation on the replication fidelity.

By using optical tweezers, Morin *et al.* had shown that DNAP translocation is not powered by PPi release or dNTP binding[32] and it’s indeed a thermal ratchet process. So the simplest reaction scheme accounting for DNAP translocation can be depicted as Fig.10. *k*_*t*_ and *r*_*t*_ are the forward and the backward translocation rate respectively. *E*_*pre*_ · *D*_*i*_ and *E*_*post*_ · *D*_*i*_ indicate the pre-translocation and the post-translocation state of DNAP respectively. Here, the primer terminal can only switch intramolecularly between *E*_*e*_ and *E*_*pre*_ (but not *E*_*post*_), according to the experimental observation [31]. We also assume DNAP can bind DNA either in state *E*_*pre*_ · *D*_*i*_ or in state *E*_*post*_ · *D*_*i*_ with possibly different binding rates and dissociation rates.

**FIG. 10:**
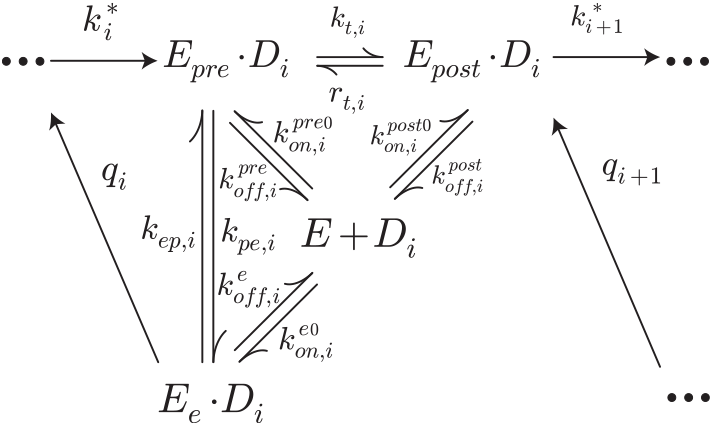
The multi-step reaction scheme of *exo*^+^-DNAP including the translocation step.

This complex scheme can be reduced to the minimal scheme Fig.1 by using the FP analysis. The obtained effective rates are given by (details see SM Sec.IV A)

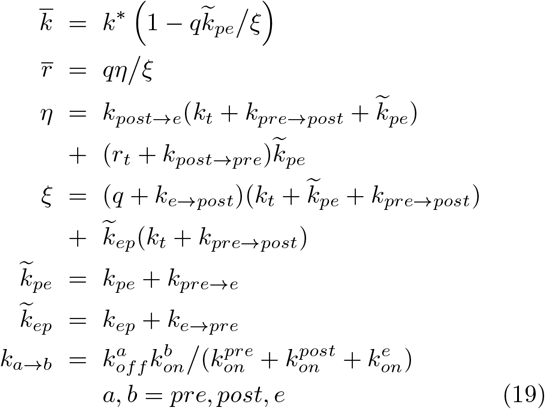

here *k*\* is defined by Eq.(7), 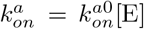. The total true fidelity *ℱ*_*i*_(≡ *ℱ*_*ini,i*_ · *ℱ*_*pro,i*_) can be approximated by *F*_*i*_(≡ *F*_*ini,i*_ · *F*_*pro,i*_), as given below

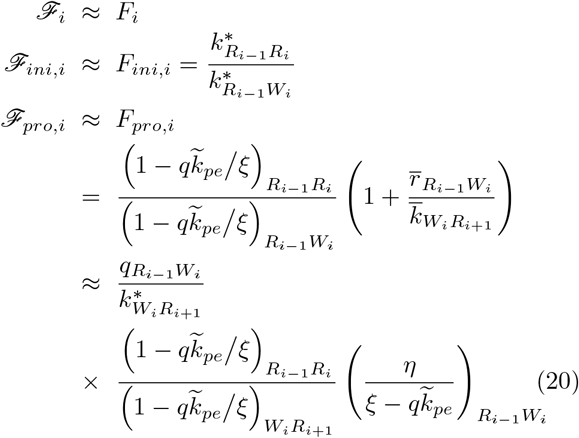

Here *ℱ*_*ini,i*_ is the initial discrimination and *ℱ*_*pro,i*_ is the proofreading efficiency, which are esitmated by *F*_*ini,i*_ and *F_pro,i_* respectively. *F*_*ini,i*_ is obtained by applying Eq.(3) with the effective rates 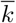 in Eq.(19) while setting *q* = 0. The relative deviations of *F*_*i*_, *F*_*ini,i*_, *F*_*pro,i*_ from *ℱ*_*i*_, *ℱ*_*ini,i*_, *ℱ*_*pro,i*_ are less than 10% (for details see SM Sec.IV A).

On the other hand, we can calculate the fidelity defined by the kinetic assays. Following the same logic in Sec.III C 1, we obtain the specificity constant by the transient-state assay of the mutant *exo*^−^-DNAP (for details see SM Sec.IV B.)

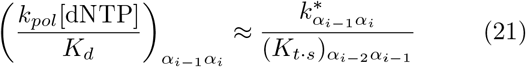

Here *α* = *R, W*. 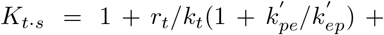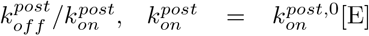. Eq.(21) gives a rough estimate of the specificity constant with less than 200% relative deviation. Similarly, the steady-state assay also define another specificity constant, 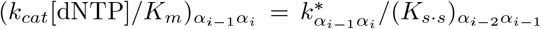, with 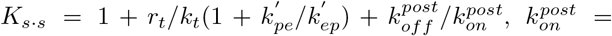 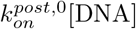. Since *K*_*t*·*s*_ (or *K*_*s* · *s*_) is independent on the incoming dNTP *α*_*i*_, the initial discrimination can be measured by the steady-state assay with high precision or roughly estimated by the transient-state assay, *i.e*. *F*_*ini*_ = *f*_*s·s,ini*_ and *F*_*ini*_ ≈ *f*_*t* · *s*,*ini*_ (with no order of magnitude difference).

The transient-state excision velocity 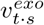 can be calculated as follows

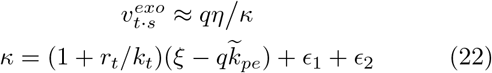

*∈*_1_ and *∈*_2_ are very complex functions of the kinetic rates which are too lengthy to give here. Eq.(22) offers a rough estimate of the real 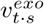 with less than 300% deviation (see details in SM Sec.IV B).

With Eq.(21)(22), the proofreading efficiency *f*_*pro*_ is defined by the transient-state assay as

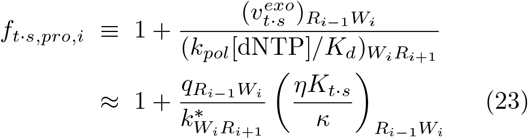

here ≈ means less than 300% deviation. We compare this rough estimate on *f*_*t* · *s,pro*_ to *F*_*pro*_ defined by Eq.(20). They might be approximately equal only under some conditions. For instance, if the kinetic rates satisfy the following conditions

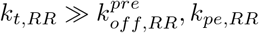, *k*_*pe,RR*_
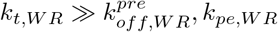, *k*_*pe,W R*_
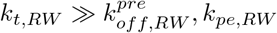, *k*_*pe,RW*_ and 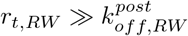.
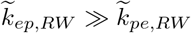 and 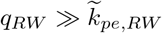.

Here ≫ means more than one order of magnitude higher. Under such conditions, one can estimate that 0.69 < *F*_*pro*_/*f*_*t* · *s,pro*_ < 1.20, the relative deviation of *f*_*t* ·*s,pro*_ from *F*_*pro*_ is less than 31% (details see SM Sec.IV B). Here we have assumed *K*_*t* · *s*_ ≈ 1 + *r*_*t*_/*k*_*t*_, since the DNAP concentration [E] can be set large enough in the transient-state assays to ensure 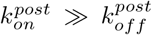 and the transfer rates 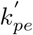 and 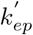 of the mutant DNAP often satisfies 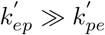.

In general, *f*_*t* · *s,pro*_ can be much different from *F*_*pro*_ in a wide range of kinetic parameters, *e.g.* the translocation in the presence of a terminal mismatch may be very slow [33] so that the above condition (3) is violated. Fig.11 shows an example. Since some kinetic rates for terminal mismatch *RW* (for simplicity we omit the subscript in this example), *e.g.* 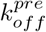 and 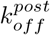, are unknown from the experiments, we just assume that 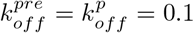 (see 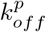 in Table II) and 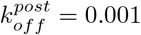. Other rates is set as explained in Sec.III B2. The ratio *r_t_/k_t_* ranges from 10^−2^ to 10^2^, corresponding to a free energy difference (between the *Pre* and *Post* states) of a few *k*_*B*_*T*. *q* = 1, 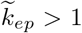 and 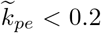, so the condition (4) still holds. As shown in the figure, the ratio 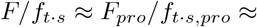 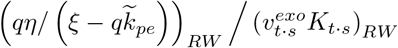 can become larger than 10 when *k*_*t*_ and *r*_*t*_ get small enough. Here 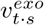 is precisely computed by numerically solving the original kinetic equations, and *K*_*t* · *s*_ ≈ 1+*r*_*t*_/*k*_*t*_ as discussed above.

**FIG. 11:**
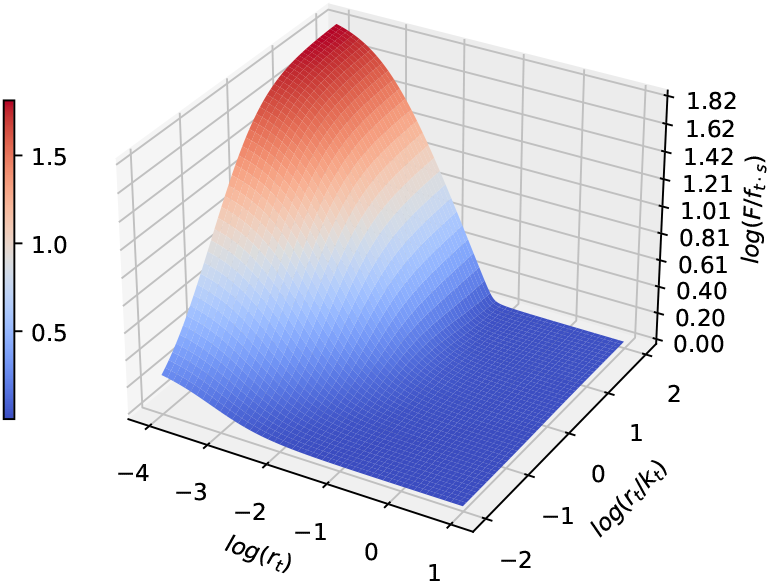
The ratio *F/f*_*t* · *s*_ becomes larger than 10 when *k*_*t*_ and *r*_*t*_ get small enough. The kinetic rates (for terminal mismatch *RW*): *q* = 1, *k*_*pe*_ = 0.1, *k*_*ep*_ = 1, 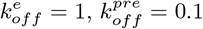, 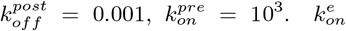 and 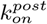 are calculated by thermodynamic constraints 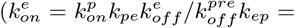 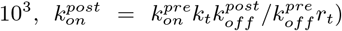. The binding rates are always large enough when adjusting *k*_*t*_ and *r*_*t*_ to ensure 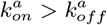, *a* = *pre, post, e* and 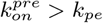.

Fig.12 shows another example where the condition (1)(2)(3) hold but the condition (4) is violated. We assume for the terminal mismatch *RW*, 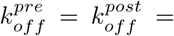 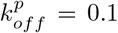 (see 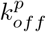 in Table II). 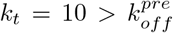 and 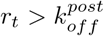 which ensures the condition (1). Other kinetic rates are set as explained in Sec.III B 2. As shown, the ratio *F*/*f*_*t* · *s*_ may become larger than 10 when *k*_*pe*_ gets large enough.

**FIG. 12:**
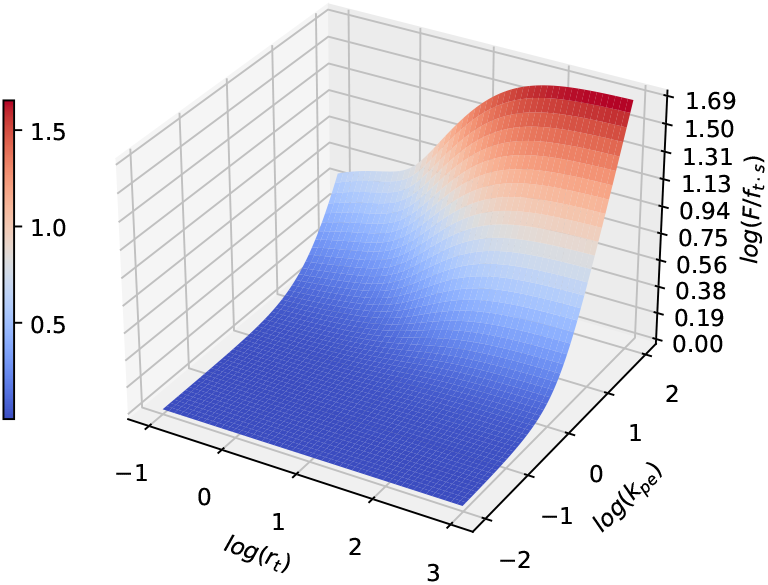
The ratio *F*/*f*_*t* · *s*_ may become larger than 10 when *k*_*pe*_ gets large enough. The kinetic rates (for terminal mismatch *RW*): *q* = 1, *k*_*ep*_ = 1, 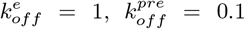 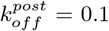, *k*_*t*_ = 10, 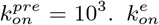 and 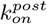 are calculated by thermodynamic constraints 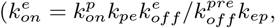 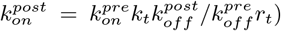. The binding rates are always large enough when adjusting *k*_*pe*_ and *r*_*t*_ to ensure 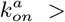 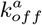, *a* = *pre, post, e* and 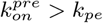.

However, the condition (2) may also not hold for real DNAPs. Since the buried mismatch may slow down DNAP translocation, 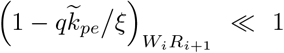 may hold if 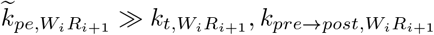 and 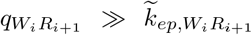. On the other hand, kinetic rates for terminal *WR* are present in *F*_*pro*_ (in the factor 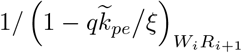, see Eq.(20)) but totally absent from *f*_*t* · *s,pro*_ (see Eq.(23)). So, *F*_*pro*_ can become much larger if 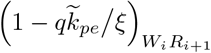 gets much smaller than 1, while other kinetic rates for terminal *RW* are fixed. That is to say, *F/f*_*t* · *s*_ may become even more than one order of magnitude larger than that shown by the red region in Fig.11 and Fig.12.

In total, if the terminal mismatch or the buried mismatch severely slows down the DNAP translocation, the transient-state assay is likely to underestimate the fidelity even by one or more orders of magnitude. Unfortunately, so far as we know, there have been no experimental studies on the translocation kinetics of any DNAP in the presence of the terminal mismatch or the buried mismatch. So, in this paper we can not further evaluate the reliability of the reported transient-state assays on specific DNAPs in the literature. Future experimental studies should offer more information on this issue.

## IV. SUMMARY

The conventional kinetic assays of DNAP fidelity, *i.e.* the steady-state assay or the transient-state assay, have indicated that the initial discrimination *f*_*ini*_ is about 10^4∼5^ and the proofreading efficiency *f*_*pro*_ is about 10^2∼3^ [18]. Although these assays have been widely used for decades and these estimates of *f*_*ini*_ and *f*_*pro*_ have been widely cited in the literature, they are not unquestionable since the logic underlying these methods are not well founded. No rigorous theories about the true fidelity *ℱ* have ever been proposed, and its relation to the operationally defined *f*_*s* · *s*_ or *f*_*t* · *s*_ has never been clarified.

In this paper, we examined carefully the relations between *f*_*s* · *s*_, *f*_*t* · *s*_ and *F*, based on the FP method recently proposed by us to investigate the true fidelity of *exo*^−^-DNAP or *exo*^+^-DNAP. We conclude that for *exo*^−^-DNAP, the steady-state assay can measure *ℱ* with very high precision, while the transient-state assay offers a rough estimate of *ℱ* (with no order of magnitude difference), just by measuring the specificity constant (*k*_*cat*_/*K*_*m*_ or *k*_*pol*_/*K*_*d*_).

For *exo*^+^-DNAP, however, the situation is more complicated. The steady-state assay or the transient-state assay can still be used to measure the initial discrimination *ℱ*_*ini*_, as done for *exo*^−^-DNAP (so the above cited estimates of *f*_*ini*_ are reliable). But either method fails to directly measure the effective excision rate and the effective elongation rate, and thus in principle they can not characterize the proofreading efficiency *ℱ*_*pro*_. So the widely cited estimates *f*_*pro*_ ∼ 10^2∼3^ are very suspicious. Our analysis shows that these kinetic assays may be valid to measure *ℱ*_*pro*_ only if the involved rate constants satisfy some conditions. If there are no further evidences to support these conditions, the assays *per se* may largely underestimate *ℱ*_*pro*_. Even when all these conditions are met, there were still quite different strategies in using these assays which often give inconsistent results. In this paper, we have shown definitely that there is only one proper way to estimate *F*_*pro*_ by using the transient-state assay.

How can one estimate the true fidelity unambiguously? A possible way is to dissect the reaction mechanism, *i.e.* measuring the rate constants of each step by transient-state experiments [17, 23, 34–38], and then calculate the effective rate according to Eq.(7) and Eq.(8). This is a perfect approach but needs heavy work. Theoretically, there is another approach, a single-molecule assay based on the FP analysis, to directly measure the effective rates. Simply put, in the framework of the first-passage theory, each effective rate required to calculate the true fidelity can be interpreted as the reciprocal of the residence time at the corresponding state in the stochastic dNTP incorporation process or the terminal excision process at the single-DNA level. This might be done, at least in principle, by analyzing the stochastic trajectories obtained in the single-molecule experiments. We do not discuss it here and leave the details in SM Sec.V. We hope that this may inspire future single-molecule assays on DNAP fidelity.

Last but not least, we have focused on the first-order (nearest) neighbor effect in this paper and just mentioned the higher-order neighbor effects which may also be important to DNAP fidelity. As shown in Sec.III C 2, the second-order (next-nearest) neighbor effect may even be more significant than the first-order effect in the proof-reading of human mitochondrial DNAP. Although this conclusion is based on the rough estimates given there, we believe that it’s somewhat universal. The proofreading efficiency *ℱ*_*i*_ of *exo*^+^-DNAP consists of two factors, 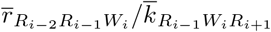 and 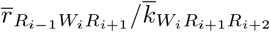, representing the first-order and the second-order proof-reading efficiency respectively. These two factors are both dependent on the stability of the primer-template duplex. For naked dsDNA duplex, numerous experiments have shown that a penultimate mismatch leads to much lower stability than a terminal mismatch [39]. This implies that a penultimate mismatch may more significantly disturb the base stacking of the primer-template conjunction in the polymerase domain and thus the forward translocation of DNAP will be slower and the *Pol*-to-*Exo* transfer of the primer terminal will be faster, if compared with the terminal mismatch. More explicitly, it’s very likely that 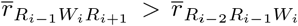 and 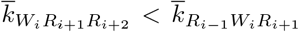. In such cases, the second-order factor may be larger than the first-order factor.

We hope that the analysis and the suggestions presented in this paper will urge serious experimental reexaminations on the conventional kinetic assays of DNAP fidelity and offer some inspirations to single-molecule experimentalists to conduct more accurate fidelity analysis.

## V ACKNOWLEDGMENTS

The authors thank the financial support by National Natural Science Foundation of China (No.11675180,11774358), the CAS Strategic Priority Research Program (No.XDA17010504), Key Research Program of Frontier Sciences of CAS (No.Y7Y1472Y61), Research Fund of Wenzhou Institute CAS (No.WIUCASYJ2020004,WIUCASQD2020009).

## Appendix A: BASICS OF THE FP METHOD

For the minimal reaction scheme (Fig.1), the master equations for replicating a given template *X*_1_*X*_2_…*X*_*L*_ (in the direction 1 → *L*) are shown below (here we only discuss the first-order neighbor effects as an example),

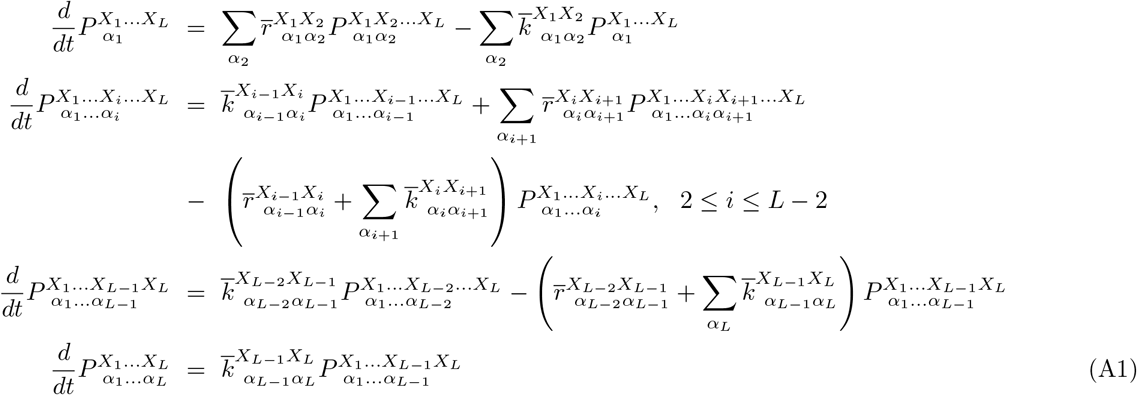

Here 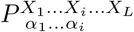 is the probability of the primer with the sequence *α*_1_…*α*_*i*_ at time *t*. 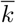 and 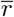 are the nucleotide incorporation rate and the excision rate respectively. In these equations, we have assumed that the first unit *α*_1_ and the last unit *α_L_* of the primer chain cannot be excised (the boundary condition), and *α*_1_ may be R or W with the given probability 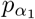 (the initial condition).

We are concerned only about the sequence distribution of the final products 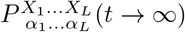, which can be given by integrating Eq.(A1)

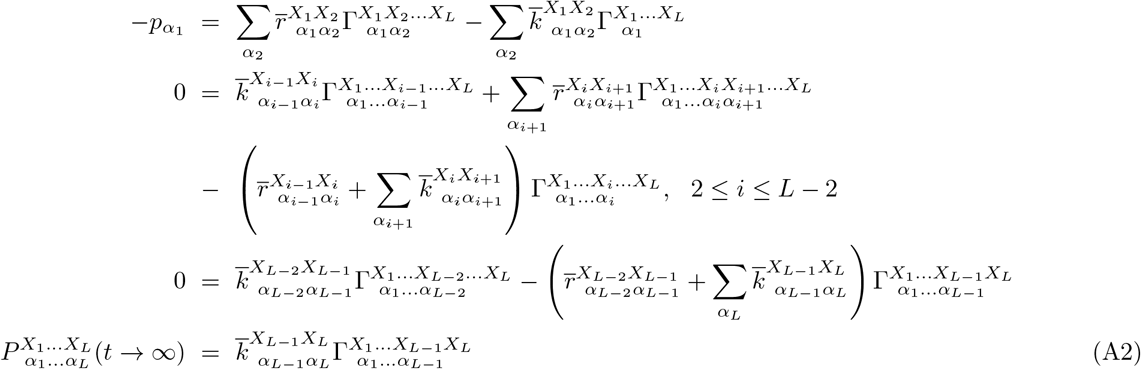

Here 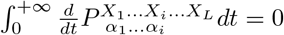 *dt* = 0 for any 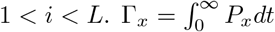 is precisely the average residence time at the state *x* (see Appendix A of Ref.[20]). One can solve these equations numerically to obtain 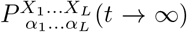 and directly give the site-specific fidelity *ℱ*_*i*_.

*ℱ*_*i*_ can be numerically computed for any given kinetic parameters. It can also be calculated analytically, though approximately, under some restrictive conditions on the parameters which seem quite reasonable for real DNA replications. That is, 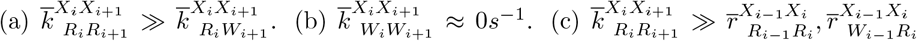 Under such conditions, one can get the approximate expressions of the true fidelity, as given by Eq.(1)-(5). Details can be found in SM Sec.I A.

## APPENDIX B: THE UNIQUE REDUCTION OF THE MULTI-STEP SCHEME TO THE MINIMAL SCHEME

The FP method can also be applied to more complex reaction schemes such as the multi-step scheme in Fig.3. These schemes can be reduced uniquely to the minimal scheme Fig.1 and hence the true fidelity *F* can be calculated by Eq.(1)-(5). Here we take the scheme Fig.3(B) as an example to illustrate the reduction procedure.

Similar to Appendix A, one can write the master equations for all the states in the reaction scheme of replicating a template *X*_1_…*X_L_* and integrate these equations to give the sequence distribution of the final products 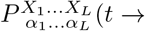 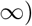. Here we only show the integrated equations for the variables 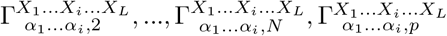 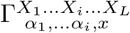corresponds to *E*_*x*_ · *D*_*i*_ in Fig.3(B), *x* = 2…*N*, *p*)

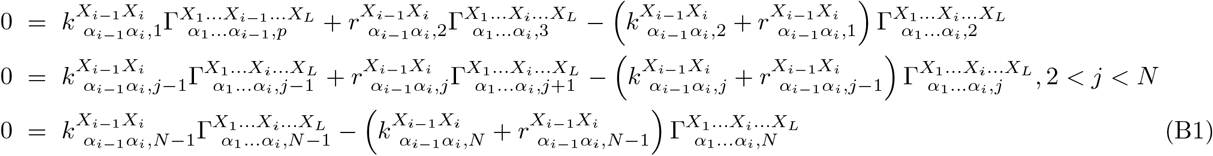

and

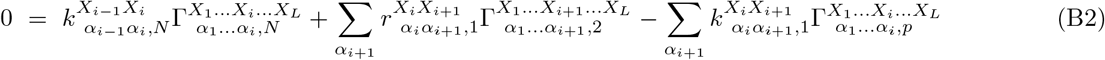

Here 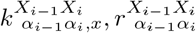 correspond to *k*_*x,i*_, *r*_*x,i*_ in Fig.3(B). According to Eq.(B1), the variables 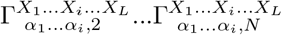 can be solved as functions of 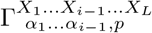. There is a similar system of equations for 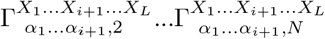, and 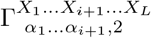 can also be solved as a function of 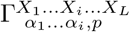. So Eq.(B2) can be rewritten as

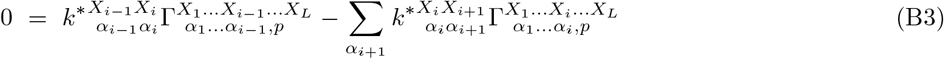

with *k*\* given by Eq.(7). Comparing Eq.(B3) with Eq.(A2), one can identify *k*\* as the effective incorporation rate 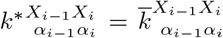 if 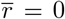. No approximations such as the steady-state assumption or the quasi-equilibrium assumption are needed in the reduction procedure. It should be noted that the above reduction is unique in the sense that other reduction procedure eliminating 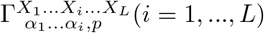 will lead to equations much different from Eq.(A2). For example, the reduced integrated equation with retained 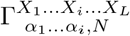 is

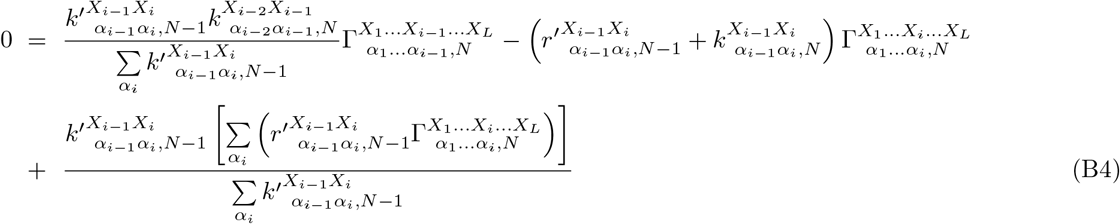

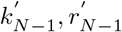 is given by Eq.(7). The form of Eq.(B4) is totally different from Eq.(B3) and Eq.(A2), so no effective incorporation rates can be properly defined and Eq.(2) and Eq.(A2) cannot be used to calculate the fidelity.

Similarly, the complex scheme Fig.3(A) can also be mapped to Fig.1 by the same logic. One can write the complete system of the integrated equations, and eliminate 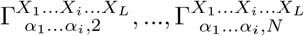 (correspoding to *E*_2_ · *D*_*i*_, …, *E_N_* · *D*_*i*_) while retain 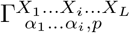 (corresponding to *E*_*p*_ · *D*_*i*_). This procedure leads to exactly the same equations as Eq.(A2) (with only 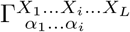 replaced by 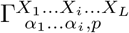), which defines the effective incorporation rate and the effective excision rate rigorously (Eq.(7)(8)) and thus the probability distribution of the final products 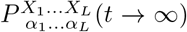 can be calculated in terms of these effective rates. Other reduction procedures which eliminates 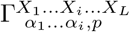 will result in equations totally different from Eq.(A2) and thus no effective rates can be properly defined.

## References

[1] T. A. Trautner, M. N. Swartz, and A. Kornberg, Proc. Natl. Acad. Sci. U.S.A. 48, 449 (1962).

[2] Z. W. Hall and I. R. Lehman, J. Mol. Biol. 36, 321 (1968).

[3] D. F. Lee, J. Lu, S. Chang, J. J. Loparo, and X. S. Xie, Nucleic Acids Res. 44, e118 (2016).

[4] A. M. de Paz, T. R. Cybulski, A. H. Marblestone, B. M. Zamft, G. M. Church, E. S. Boyden, K. P. Kording, and K. E. Tyo, Nucleic Acids Res. 46, e78 (2018).

[5] K. Clayton and W. Branscomb, J. Biol. Chem. 254, 1902 (1979).

[6] J. G. Bertram, K. Oertell, J. Petruska, and M. F. Goodman, Biochemistry 49, 20 (2010).

[7] J. Hopfield, Proc. Natl. Acad. Sci. U.S.A. 71, 4135 (1974).

[8] J. Ninio, Biochimie 57, 587 (1975).

[9] A. R. Fersht, Enzyme Structure and Mechanism (W.H.Freeman & Co Ltd., 1985), 2nd ed.

[10] A. R. Fersht, Proc. R. Soc. London, Ser. B 187, 397 (1974).

[11] W. M. Kati, K. A. Johnson, L. F. Jerva, and K. S. Anderson, J. Biol. Chem. 267, 25988 (1992).

[12] K. A. Johnson, Annu. Rev. Biochem. 62, 685 (1993).

[13] A. R. Fersht, J. W. Knill-Jones, and W. C. Tsui, J. Mol. Biol. 156, 37 (1982).

[14] A. R. Fersht, Proc. Natl. Acad. Sci. U.S.A. 76, 4946 (1979).

[15] B. M. Wingert, E. E. Parrott, and S. W. Nelson, Biochemistry 52, 7723 (2013).

[16] A. K. Vashishtha and R. D. Kuchta, Biochemistry 54, 240 (2015).

[17] M. J. Donlin, S. S. Patel, and K. A. Johnson, Biochemistry 30, 538 (1991).

[18] A. Bȩbenek and I. Ziuzia-Graczyk, Curr. Genet. 64, 985 (2018).

[19] P. Gaspard, Phys. Rev. E 96, 042403 (2017).

[20] Q. S. Li, P. D. Zheng, Y. G. Shu, Z. C. Ou-Yang, and M. Li, Phys. Rev. E 100, 012131 (2019).

[21] Y. S. Song, Y. G. Shu, X. Zhou, Z. C. Ou-Yang, and M. Li, J. Phys.: Condens. Matter 29, 025101 (2017).

[22] R. Lamichhane, S. Y. Berezhna, J. P. Gill, E. Van Der Schans, and D. P. Millar, J. Am. Chem. Soc. 135, 4735 (2013).

[23] I. Wong, S. S. Patel, and K. A. Johnson, Biochemistry 30, 526 (1991).

[24] A. A. Johnson and K. A. Johnson, J. Biol. Chem. 276, 38097 (2001).

[25] A. A. Johnson and K. A. Johnson, J. Biol. Chem. 276, 38090 (2001).

[26] K. A. Johnson, The Enzymes 20, 1 (1992).

[27] M. F. Goodman, S. Creighton, L. B. Bloom, J. Petruska, and T. A. Kunkel, Crit. Rev. Biochem. Mol. Biol. 28, 83 (1993).

[28] J. M. Dahl, A. H. Mai, G. M. Cherf, N. N. Jetha, D. R. Garalde, A. Marziali, M. Akeson, H. Wang, and K. R. Lieberman, J. Biol. Chem. 287, 13407 (2012).

[29] K. R. Lieberman, J. M. Dahl, A. H. Mai, M. Akeson, and H. Wang, J. Am. Chem. Soc. 134, 18816 (2012).

[30] K. R. Lieberman, J. M. Dahl, A. H. Mai, A. Cox, M. Akeson, and H. Wang, J. Am. Chem. Soc. 135, 9149 (2013).

[31] K. R. Lieberman, J. M. Dahl, and H. Wang, J. Am. Chem. Soc. 136, 7117 (2014).

[32] J. A. Morin, F. J. Cao, J. M. Lázaro, J. R. Arias-Gonzalez, J. M. Valpuesta, J. L. Carrascosa, M. Salas, and B. Ibarra, Nucleic Acids Res. 43, 3643 (2015).

[33] Z. Ren, Nucleic Acids Res. 44, 7457 (2016).

[34] Y. C. Tsai and K. A. Johnson, Biochemistry 45, 9675 (2006).

[35] V. Purohit, N. D. Grindley, and C. M. Joyce, Biochemistry 42, 10200 (2003).

[36] C. M. Joyce, O. Potapova, A. M. DeLucia, X. Huang, V. P. Basu, and N. D. Grindley, Biochemistry 47, 6103 (2008).

[37] Y. Santoso, C. M. Joyce, O. Potapova, L. Le Reste, J. Hohlbein, J. P. Torella, N. D. Grindley, and A. N. Kapanidis, Proc. Natl. Acad. Sci. U.S.A. 107, 715 (2010).

[38] J. Hohlbein, L. Aigrain, T. D. Craggs, O. Bermek, O. Potapova, P. Shoolizadeh, N. D. Grindley, C. M. Joyce, and A. N. Kapanidis, Nat. Commun. 4, 1 (2013).

[39] J. SantaLucia and D. Hicks, Annu. Rev. Biophys. Biomol. Struct. 33, 415 (2004).

