## Supplemental material for "Kinetic assays of DNA polymerase fidelity: a new theoretical perspective beyond Michaelis-Menten kinetics"

### I. THE FIRST-PASSAGE METHOD

In this section we first review the first-passage (FP) method applied to the minimal model of DNA replication as shown in Fig.S1, then show how to reduce various multi-step reaction schemes of $exo^{-}$-DNAP and $exo^{+}$-DNAP to the minimal scheme in order to calculate the fidelity.

#### A. The FP method for the minimal model

$$\cdots \rightleftharpoons E\cdot D_{i-1} \underset{\overline{r}_i}{\overset{\overline{k}_i}{\rightleftharpoons}} E\cdot D_i \underset{\overline{r}_{i+1}}{\overset{\overline{k}_{i+1}}{\rightleftharpoons}} E\cdot D_{i+1} \rightleftharpoons \cdots$$

Fig.S1: The minimal reaction scheme.

The FP method can be employed to directly calculate the sequence distribution of the final products when the replication of a given template is finished, from which the site-specific fidelity can be counted. Here we give the details. For the minimal reaction scheme (Fig.S1), the master equations for replicating a given template $X_1X_2...X_L$ (in the direction $1 \rightarrow L$) are shown below. For simplicity and without loss of generality, we only discuss the first-order neighbor effects.

---

*Electronic address:

$$
\begin{aligned}
\frac{d}{dt}P^{X_1\ldots X_L}_{\alpha_1} &= \sum_{\alpha_2}\overline{r}^{X_1X_2}_{\alpha_1\alpha_2}P^{X_1X_2\ldots X_L}_{\alpha_1\alpha_2} - \sum_{\alpha_2}\overline{k}^{X_1X_2}_{\alpha_1\alpha_2}P^{X_1\ldots X_L}_{\alpha_1} \\
\frac{d}{dt}P^{X_1\ldots X_i\ldots X_L}_{\alpha_1\ldots\alpha_i} &= \overline{k}^{X_{i-1}X_i}_{\alpha_{i-1}\alpha_i}P^{X_1\ldots X_{i-1}\ldots X_L}_{\alpha_1\ldots\alpha_{i-1}} + \sum_{\alpha_{i+1}}\overline{r}^{X_iX_{i+1}}_{\alpha_i\alpha_{i+1}}P^{X_1\ldots X_iX_{i+1}\ldots X_L}_{\alpha_1\ldots\alpha_i\alpha_{i+1}} \\
&- \left(\overline{r}^{X_{i-1}X_i}_{\alpha_{i-1}\alpha_i} + \sum_{\alpha_{i+1}}\overline{k}^{X_iX_{i+1}}_{\alpha_i\alpha_{i+1}}\right)P^{X_1\ldots X_i\ldots X_L}_{\alpha_1\ldots\alpha_i}, \quad 2\le i\le L-2 \\
\frac{d}{dt}P^{X_1\ldots X_{L-1}X_L}_{\alpha_1\ldots\alpha_{L-1}} &= \overline{k}^{X_{L-2}X_{L-1}}_{\alpha_{L-2}\alpha_{L-1}}P^{X_1\ldots X_{L-2}\ldots X_L}_{\alpha_1\ldots\alpha_{L-2}} - \left(\overline{r}^{X_{L-2}X_{L-1}}_{\alpha_{L-2}\alpha_{L-1}} + \sum_{\alpha_L}\overline{k}^{X_{L-1}X_L}_{\alpha_{L-1}\alpha_L}\right)P^{X_1\ldots X_{L-1}X_L}_{\alpha_1\ldots\alpha_{L-1}} \\
\frac{d}{dt}P^{X_1\ldots X_L}_{\alpha_1\ldots\alpha_L} &= \overline{k}^{X_{L-1}X_L}_{\alpha_{L-1}\alpha_L}P^{X_1\ldots X_{L-1}X_L}_{\alpha_1\ldots\alpha_{L-1}} \qquad (1)
\end{aligned}
$$

Here $P^{X_1\ldots X_i\ldots X_L}_{\alpha_1\ldots\alpha_i}$ is the probability of the primer chain with the sequence $\alpha_1\ldots\alpha_i$ at time $t$, indicated by the state symbol $E\cdot D_i$ in Fig.S1. $\overline{k}$ and $\overline{r}$ are the nucleotide incorporation rate and the excision rate respectively. In these equations, we have assumed that the first unit $\alpha_1$ and the last unit $\alpha_L$ of the primer chain cannot be excised (the reflecting and the absorbing boundary conditions respectively), and $\alpha_1$ may be any one of the four nucleotides (A, G, T, C) with the given probability $p_{\alpha_1}$ (the initial condition).

We are concerned only about the sequence distribution of the final products $P^{X_1\ldots X_L}_{\alpha_1\ldots\alpha_L}(t\to\infty)$, which can be given by integrating Eq.(1)

$$
\begin{aligned}
-p_{\alpha_1} &= \sum_{\alpha_2}\overline{r}^{X_1X_2}_{\alpha_1\alpha_2}\Gamma^{X_1X_2\ldots X_L}_{\alpha_1\alpha_2} - \sum_{\alpha_2}\overline{k}^{X_1X_2}_{\alpha_1\alpha_2}\Gamma^{X_1\ldots X_L}_{\alpha_1} \\
0 &= \overline{k}^{X_{i-1}X_i}_{\alpha_{i-1}\alpha_i}\Gamma^{X_1\ldots X_{i-1}\ldots X_L}_{\alpha_1\ldots\alpha_{i-1}} + \sum_{\alpha_{i+1}}\overline{r}^{X_iX_{i+1}}_{\alpha_i\alpha_{i+1}}\Gamma^{X_1\ldots X_iX_{i+1}\ldots X_L}_{\alpha_1\ldots\alpha_i\alpha_{i+1}} \\
&- \left(\overline{r}^{X_{i-1}X_i}_{\alpha_{i-1}\alpha_i} + \sum_{\alpha_{i+1}}\overline{k}^{X_iX_{i+1}}_{\alpha_i\alpha_{i+1}}\right)\Gamma^{X_1\ldots X_i\ldots X_L}_{\alpha_1\ldots\alpha_i}, \quad 2\le i\le L-2 \\
0 &= \overline{k}^{X_{L-2}X_{L-1}}_{\alpha_{L-2}\alpha_{L-1}}\Gamma^{X_1\ldots X_{L-2}\ldots X_L}_{\alpha_1\ldots\alpha_{L-2}} - \left(\overline{r}^{X_{L-2}X_{L-1}}_{\alpha_{L-2}\alpha_{L-1}} + \sum_{\alpha_L}\overline{k}^{X_{L-1}X_L}_{\alpha_{L-1}\alpha_L}\right)\Gamma^{X_1\ldots X_{L-1}X_L}_{\alpha_1\ldots\alpha_{L-1}} \\
P^{X_1\ldots X_L}_{\alpha_1\ldots\alpha_L}(t\to\infty) &= \overline{k}^{X_{L-1}X_L}_{\alpha_{L-1}\alpha_L}\Gamma^{X_1\ldots X_{L-1}X_L}_{\alpha_1\ldots\alpha_{L-1}} \qquad (2)
\end{aligned}
$$

$\Gamma_x = \int_0^\infty P_x dt$ is precisely the average residence time at the state $x$ (details see also Appendix A of Ref.[1]). By solving Eq.(2), one can obtain

$$\begin{aligned}
P^{X_1...X_L}_{\alpha_1...\alpha_L}(t\to\infty) &= \left(p_{\alpha_1}/g^{X_1...X_L}_{\alpha_1}\right)\Pi^{X_1X_2...X_L}_{\alpha_1\alpha_2}\Pi^{X_2X_3...X_L}_{\alpha_2\alpha_3}...\Pi^{X_iX_{i+1}...X_L}_{\alpha_i\alpha_{i+1}}...\Pi^{X_{L-2}X_{L-1}X_L}_{\alpha_{L-2}\alpha_{L-1}}\\
\Pi^{X_iX_{i+1}...X_L}_{\alpha_i\alpha_{i+1}} &= \overline{k}^{X_iX_{i+1}}_{\alpha_i\alpha_{i+1}}/\left(\overline{r}^{X_iX_{i+1}}_{\alpha_i\alpha_{i+1}}+g^{X_{i+1}...X_L}_{\alpha_{i+1}}\right) \quad ,i=1,2,...,L-2\\
g^{X_{i+1}...X_L}_{\alpha_{i+1}} &= \sum_{\alpha_{i+2}}\left(\Pi^{X_{i+1}X_{i+2}...X_L}_{\alpha_{i+1}\alpha_{i+2}}g^{X_{i+2}...X_L}_{\alpha_{i+2}}\right)\\
g^{X_{L-1}X_L}_{\alpha_{L-1}} &= \sum_{\alpha_L}\overline{k}^{X_{L-1}X_L}_{\alpha_{L-1}\alpha_L}
\end{aligned} \tag{3}$$

Eq.(3) can be transformed into a more intuitive form, a forward inhomogeneous Markov chain, as below

$$\begin{aligned}
P^{X_1...X_L}_{\alpha_1...\alpha_L}(t\to\infty) &= p_{\alpha_1}M^{X_1X_2...X_L}_{\alpha_1\alpha_2}M^{X_2X_3...X_L}_{\alpha_2\alpha_3}...M^{X_iX_{i+1}...X_L}_{\alpha_i\alpha_{i+1}}...M^{X_{L-1}X_L}_{\alpha_{L-1}\alpha_L}\\
M^{X_iX_{i+1}...X_L}_{\alpha_i\alpha_{i+1}} &= \Pi^{X_iX_{i+1}...X_L}_{\alpha_i\alpha_{i+1}}g^{X_{i+1}...X_L}_{\alpha_{i+1}}/g^{X_i...X_L}_{\alpha_i}\\
M^{X_{L-1}X_L}_{\alpha_{L-1}\alpha_L} &= \overline{k}^{X_{L-1}X_L}_{\alpha_{L-1}\alpha_L}/g^{X_{L-1}X_L}_{\alpha_{L-1}}
\end{aligned} \tag{4}$$

$M$ is a stochastic matrix in which the sum of each row equals to 1. Noting there is one match ($R_i$) but three mismatches ($W_i, i=1,2,3$) at each site $i$, one can write $M$ as

$$M_{i-1} = \begin{pmatrix}
M^{X_{i-1}X_i...X_L}_{R_{i-1}R_i} & M^{X_{i-1}X_i...X_L}_{R_{i-1}W_{i,1}} & M^{X_{i-1}X_i...X_L}_{R_{i-1}W_{i,2}} & M^{X_{i-1}X_i...X_L}_{R_{i-1}W_{i,3}}\\
M^{X_{i-1}X_i...X_L}_{W_{i-1,1}R_i} & M^{X_{i-1}X_i...X_L}_{W_{i-1,1}W_{i,1}} & M^{X_{i-1}X_i...X_L}_{W_{i-1,1}W_{i,2}} & M^{X_{i-1}X_i...X_L}_{W_{i-1,1}W_{i,3}}\\
M^{X_{i-1}X_i...X_L}_{W_{i-1,2}R_i} & M^{X_{i-1}X_i...X_L}_{W_{i-1,2}W_{i,1}} & M^{X_{i-1}X_i...X_L}_{W_{i-1,2}W_{i,2}} & M^{X_{i-1}X_i...X_L}_{W_{i-1,2}W_{i,3}}\\
M^{X_{i-1}X_i...X_L}_{W_{i-1,3}R_i} & M^{X_{i-1}X_i...X_L}_{W_{i-1,3}W_{i,1}} & M^{X_{i-1}X_i...X_L}_{W_{i-1,3}W_{i,2}} & M^{X_{i-1}X_i...X_L}_{W_{i-1,3}W_{i,3}}
\end{pmatrix} \tag{5}$$

So the probability distribution at site $i$ $P^{X_i}_{\alpha_i} = \sum_{\{\alpha_j\},j\neq i} P^{X_1...X_L}_{\alpha_1...\alpha_L}(t\to\infty)$ is given by

$$\left(P^{X_i}_{R_i}\ P^{X_i}_{W_{i,1}}\ P^{X_i}_{W_{i,2}}\ P^{X_i}_{W_{i,3}}\right) = \left(p_{R_1}\ p_{W_{1,1}}\ p_{W_{1,2}}\ p_{W_{1,3}}\right)M_1M_2...M_{i-1} \tag{6}$$

According to Eq.(3), we define

$$\mathcal{U}^{X_{i-1}X_i}_{\alpha_{i-1}\alpha_i} \equiv \Pi^{X_{i-1}X_i...X_L}_{\alpha_{i-1}\alpha_i}g^{X_i...X_L}_{\alpha_i} = \overline{k}^{X_{i-1}X_i}_{\alpha_{i-1}\alpha_i}\Bigg/\left(1+\frac{\overline{r}^{X_{i-1}X_i}_{\alpha_{i-1}\alpha_i}}{\mathcal{U}^{X_iX_{i+1}}_{\alpha_iR_{i+1}}+\sum\limits_{W_{i+1}}\mathcal{U}^{X_iX_{i+1}}_{\alpha_iW_{i+1}}}\right) \tag{7}$$

$\sum\limits_{W_i}$ means summing over all the three mismatches at site $i$. The transfer matrix $M$ can now be rewritten as

$$M^{X_{i-1}X_i...X_L}_{\alpha_{i-1}\alpha_i} = \frac{\mathcal{U}^{X_{i-1}X_i}_{\alpha_{i-1}\alpha_i}}{\mathcal{U}^{X_{i-1}X_i}_{\alpha_{i-1}R_i}+\sum\limits_{W_i}\mathcal{U}^{X_{i-1}X_i}_{\alpha_{i-1}W_i}} \tag{8}$$

The overall fidelity at site $i$, considering all the three mismatches, is defined by

$$\mathscr{F}_i^{ov} \equiv \frac{P_{R_i}}{\sum\limits_{W_i} P_{W_i}} = \left[\sum_{W_i} \frac{P_{W_i}}{P_{R_i}}\right]^{-1} \tag{9}$$

Below we discuss the specific fidelity for any one mismatch $W_i$, since the kinetic assays are always concerned about the mismatch-specific fidelity. According to Eq.(5)(6), it can be written as

$$\begin{aligned}
\mathscr{F}_i \equiv \frac{P_{R_i}}{P_{W_i}} &= \frac{P_{R_{i-1}} M^{X_{i-1}X_i\ldots X_L}_{R_{i-1}R_i} + \sum\limits_{W_{i-1}} \left(P_{W_{i-1}} M^{X_{i-1}X_i\ldots X_L}_{W_{i-1}R_i}\right)}{P_{R_{i-1}} M^{X_{i-1}X_i\ldots X_L}_{R_{i-1}W_i} + \sum\limits_{W_{i-1}} \left(P_{W_{i-1}} M^{X_{i-1}X_i\ldots X_L}_{W_{i-1}W_i}\right)} \\
&= \frac{P_{R_{i-1}}\left[\mathcal{U}^{X_{i-1}X_i}_{R_{i-1}R_i} \Big/ \left(\mathcal{U}^{X_{i-1}X_i}_{R_{i-1}R_i} + \sum\limits_{W_i}\mathcal{U}^{X_{i-1}X_i}_{R_{i-1}W_i}\right)\right] + \sum\limits_{W_{i-1}}\left(P_{W_{i-1}}\left[\mathcal{U}^{X_{i-1}X_i}_{W_{i-1}R_i} \Big/ \left(\mathcal{U}^{X_{i-1}X_i}_{W_{i-1}R_i} + \sum\limits_{W_i}\mathcal{U}^{X_{i-1}X_i}_{W_{i-1}W_i}\right)\right]\right)}{P_{R_{i-1}}\left[\mathcal{U}^{X_{i-1}X_i}_{R_{i-1}W_i} \Big/ \left(\mathcal{U}^{X_{i-1}X_i}_{R_{i-1}R_i} + \sum\limits_{W_i}\mathcal{U}^{X_{i-1}X_i}_{R_{i-1}W_i}\right)\right] + \sum\limits_{W_{i-1}}\left(P_{W_{i-1}}\left[\mathcal{U}^{X_{i-1}X_i}_{W_{i-1}W_i} \Big/ \left(\mathcal{U}^{X_{i-1}X_i}_{W_{i-1}R_i} + \sum\limits_{W_i}\mathcal{U}^{X_{i-1}X_i}_{W_{i-1}W_i}\right)\right]\right)} \\
&= \left(\frac{\mathcal{U}^{X_{i-1}X_i}_{R_{i-1}R_i}}{\mathcal{U}^{X_{i-1}X_i}_{R_{i-1}W_i}}\right)\left(\frac{1 + \sum\limits_{W_{i-1}}\left(\mathcal{H}_{W_{i-1}}\mathcal{U}^{X_{i-1}X_i}_{W_{i-1}R_i} \Big/ \mathcal{U}^{X_{i-1}X_i}_{R_{i-1}R_i}\right)}{1 + \sum\limits_{W_{i-1}}\left(\mathcal{H}_{W_{i-1}}\mathcal{U}^{X_{i-1}X_i}_{W_{i-1}W_i} \Big/ \mathcal{U}^{X_{i-1}X_i}_{R_{i-1}W_i}\right)}\right)
\end{aligned} \tag{10}$$

in which

$$\mathcal{H}_{W_{i-1}} \equiv \frac{P_{W_{i-1}}}{P_{R_{i-1}}} \frac{\mathcal{U}^{X_{i-1}X_i}_{R_{i-1}R_i} + \sum\limits_{W_i}\mathcal{U}^{X_{i-1}X_i}_{R_{i-1}W_i}}{\mathcal{U}^{X_{i-1}X_i}_{W_{i-1}R_i} + \sum\limits_{W_i}\mathcal{U}^{X_{i-1}X_i}_{W_{i-1}W_i}} \tag{11}$$

$\mathscr{F}_i$ can be numerically computed for any given kinetic parameters. It can also be calculated analytically, though approximately, under the following conditions (biologically-relevant conditions) which seem quite reasonable for real DNA replications

(a) $\overline{k}^{X_iX_{i+1}}_{R_iR_{i+1}} \gg \overline{k}^{X_iX_{i+1}}_{R_iW_{i+1}}$, which means that the addition of $R$ is always much faster than that of $W$.

(b) $\overline{k}^{X_iX_{i+1}}_{W_iW_{i+1}} \approx 0s^{-1}$. In biochemical or biophysical experiments, $\overline{k}_{WW}$ has never been successfully measured, so $\overline{k}_{WW} \simeq 0s^{-1}$ is always assumed in literatures, while $\overline{k}_{RR}$, $\overline{k}_{RW}$ and $\overline{k}_{WR}$ have finite values in experimental measurements. Hence, the condition $\overline{k}_{WW} \simeq 0s^{-1}$ in this paper just means that $\overline{k}_{WW}$ is arbitrarily smaller than $\overline{k}_{RR}$, $\overline{k}_{RW}$ and $\overline{k}_{WR}$.

(c) $\overline{k}^{X_iX_{i+1}}_{R_iR_{i+1}} \gg \overline{r}^{X_{i-1}X_i}_{R_{i-1}R_i}, \overline{r}^{X_{i-1}X_i}_{W_{i-1}R_i}$, which means that the successive additions of R always dominate the replication process in order to guarantee the high replication velocity.

Here $\gg$ means more than one order of magnitude higher. Under such conditions, we can estimate the upper bound and lower bound of $\mathscr{F}_i$ and give the approximate analytical expression $F_i$ . For simplicity, we omit the superscript $X$ in all the notations.

First we estimate the bounds of $\mathcal{U}_{R_{i-1}R_i}/\mathcal{U}_{R_{i-1}W_i}$ in Eq.(10). From Eq.(7), we have

$$\frac{\mathcal{U}_{R_{i-1}R_i}}{\mathcal{U}_{R_{i-1}W_i}}=\frac{\overline{k}_{R_{i-1}R_i}}{\overline{k}_{R_{i-1}W_i}}\left(1+\frac{\overline{r}_{R_{i-1}W_i}}{\mathcal{U}_{W_iR_{i+1}}+\sum\limits_{W_{i+1}}\mathcal{U}_{W_iW_{i+1}}}\right)\Bigg/\left(1+\frac{\overline{r}_{R_{i-1}R_i}}{\mathcal{U}_{R_iR_{i+1}}+\sum\limits_{W_{i+1}}\mathcal{U}_{R_iW_{i+1}}}\right) \tag{12}$$

The upper bound of $1+\overline{r}_{R_{i-1}W_i}/(\mathcal{U}_{W_iR_{i+1}}+\sum\limits_{W_{i+1}}\mathcal{U}_{W_iW_{i+1}})$ can be estimated by

$$\begin{aligned}1+\frac{\overline{r}_{R_{i-1}W_i}}{\mathcal{U}_{W_iR_{i+1}}+\sum\limits_{W_{i+1}}\mathcal{U}_{W_iW_{i+1}}} &< 1+\frac{\overline{r}_{R_{i-1}W_i}}{\mathcal{U}_{W_iR_{i+1}}}\\ &= 1+\frac{\overline{r}_{R_{i-1}W_i}}{\overline{k}_{W_iR_{i+1}}}\left(1+\frac{\overline{r}_{W_iR_{i+1}}}{\mathcal{U}_{R_{i+1}R_{i+2}}+\sum\limits_{W_{i+2}}\mathcal{U}_{R_{i+1}W_{i+2}}}\right)\\ &< 1+\frac{\overline{r}_{R_{i-1}W_i}}{\overline{k}_{W_iR_{i+1}}}\left(1+\frac{\overline{r}_{W_iR_{i+1}}}{\mathcal{U}_{R_{i+1}R_{i+2}}}\right)\\ &< 1+\frac{\overline{r}_{R_{i-1}W_i}}{\overline{k}_{W_iR_{i+1}}}\left(1+\frac{\overline{r}_{W_iR_{i+1}}}{\overline{k}_{R_{i+1}R_{i+2}}}\left(1+\frac{\overline{r}_{R_{i+1}R_{i+2}}}{\overline{k}_{R_{i+2}R_{i+3}}}\Big(1+...\Big)\right)\right)\end{aligned} \tag{13}$$

Considering the condition (c) $\overline{k}_{R_iR_{i+1}} > 10\ \overline{r}_{R_{i-1}R_i}, 10\ \overline{r}_{W_{i-1}R_i}$, one can get

$$\begin{aligned}1+\frac{\overline{r}_{R_{i-1}W_i}}{\mathcal{U}_{W_iR_{i+1}}+\sum\limits_{W_{i+1}}\mathcal{U}_{W_iW_{i+1}}} &< 1+\frac{\overline{r}_{R_{i-1}W_i}}{\overline{k}_{W_iR_{i+1}}}\left(1+0.1\Big(1+0.1\Big(1+...\Big)\Big)\right)\\ &< 1+\frac{\overline{r}_{R_{i-1}W_i}}{\overline{k}_{W_iR_{i+1}}}\frac{1}{0.9}\\ &< \left(1+\frac{\overline{r}_{R_{i-1}W_i}}{\overline{k}_{W_iR_{i+1}}}\right)\frac{1}{0.9}\end{aligned} \tag{14}$$

The lower bound can be given as below, by noticing that $\mathcal{U}_{\alpha_{i-1}\alpha_i} < \overline{k}_{\alpha_{i-1}\alpha_i}$ and condition (b) $\overline{k}_{WW} \approx 0$,

$$1+\frac{\overline{r}\ _{R_{i-1}W_i}}{\mathcal{U}_{W_iR_{i+1}}+\sum\limits_{W_{i+1}}\mathcal{U}_{W_iW_{i+1}}} > 1+\frac{\overline{r}_{R_{i-1}W_i}}{\overline{k}_{W_iR_{i+1}}} \tag{15}$$

So we finally obtain

$$\left(1+\frac{\overline{r}_{R_{i-1}W_i}}{\overline{k}_{W_iR_{i+1}}}\right)<1+\frac{\overline{r}_{R_{i-1}W_i}}{\mathcal{U}_{W_iR_{i+1}}+\sum\limits_{W_{i+1}}\mathcal{U}_{W_iW_{i+1}}}<\frac{1}{0.9}\left(1+\frac{\overline{r}_{R_{i-1}W_i}}{\overline{k}_{W_iR_{i+1}}}\right) \tag{16}$$

Following the same logic from Eq.(13) to Eq.(16), and according to the condition (c) $\overline{k}_{R_iR_{i+1}} > 10\ \overline{r}_{R_{i-1}R_i}$, one can get the upper bound and lower bound of $1+\overline{r}_{R_{i-1}R_i}/(\mathcal{U}_{R_iR_{i+1}}+\sum\limits_{W_{i+1}}\mathcal{U}_{R_iW_{i+1}})$ in Eq.(12)

$$1<1+\frac{\overline{r}_{R_{i-1}R_i}}{\mathcal{U}_{R_iR_{i+1}}+\sum\limits_{W_{i+1}}\mathcal{U}_{R_iW_{i+1}}}<\frac{1}{0.9} \tag{17}$$

Combining Eq.(12)(16)(17) , we have

$$0.9\frac{\overline{k}_{R_{i-1}R_i}}{\overline{k}_{R_{i-1}W_i}}\left(1+\frac{\overline{r}_{R_{i-1}W_i}}{\overline{k}_{W_iR_{i+1}}}\right) < \frac{\mathcal{U}_{R_{i-1}R_i}}{\mathcal{U}_{R_{i-1}W_i}} < \frac{1}{0.9}\frac{\overline{k}_{R_{i-1}R_i}}{\overline{k}_{R_{i-1}W_i}}\left(1+\frac{\overline{r}_{R_{i-1}W_i}}{\overline{k}_{W_iR_{i+1}}}\right) \tag{18}$$

On the other hand, $\mathcal{U}_{W_{i-1}W_i} < \overline{k}_{W_{i-1}W_i} \approx 0s^{-1}$ in Eq.(10), so Eq.(10)(18) give the lower bound of $\mathscr{F}_i$,

$$\mathscr{F}_i > \frac{\mathcal{U}_{R_{i-1}R_i}}{\mathcal{U}_{R_{i-1}W_i}} > 0.9\frac{\overline{k}_{R_{i-1}R_i}}{\overline{k}_{R_{i-1}W_i}}\left(1+\frac{\overline{r}_{R_{i-1}W_i}}{\overline{k}_{W_iR_{i+1}}}\right) \tag{19}$$

From Eq.(10) one can get the upper bound of $\mathscr{F}_i$,

$$\begin{aligned}\mathscr{F}_i &= \frac{\mathcal{U}_{R_{i-1}R_i}}{\mathcal{U}_{R_{i-1}W_i}}\left(\frac{1+\sum_{W_{i-1}}\left(\mathcal{H}_{W_{i-1}}\mathcal{U}_{W_{i-1}R_i}\Big/\mathcal{U}_{R_{i-1}R_i}\right)}{1+\sum_{W_{i-1}}\left(\mathcal{H}_{W_{i-1}}\mathcal{U}_{W_{i-1}W_i}\Big/\mathcal{U}_{R_{i-1}W_i}\right)}\right)\\ &< \frac{\mathcal{U}_{R_{i-1}R_i}}{\mathcal{U}_{R_{i-1}W_i}}\left(1+\sum_{W_{i-1}}\left(\mathcal{H}_{W_{i-1}}\frac{\mathcal{U}_{W_{i-1}R_i}}{\mathcal{U}_{R_{i-1}R_i}}\right)\right)\\ &< \frac{\mathcal{U}_{R_{i-1}R_i}}{\mathcal{U}_{R_{i-1}W_i}}\left(1+\sum_{W_{i-1}}\left(\frac{P_{W_{i-1}}}{P_{R_{i-1}}}\left(1+\sum_{W_i}\frac{\mathcal{U}_{R_{i-1}W_i}}{\mathcal{U}_{R_{i-1}R_i}}\right)\right)\right)\end{aligned} \tag{20}$$

Because Eq.(19) holds for any site $i$, and according to the condition (a) $\overline{k}_{R_iR_{i+1}} > 10\ \overline{k}_{R_iW_{i+1}}$ for any site $i$, we have

$$\begin{aligned}\frac{P_{W_{i-1}}}{P_{R_{i-1}}} = \frac{1}{\mathscr{F}_{i-1}} &< \frac{1}{0.9}\times 0.1 \approx 0.11\\ \frac{\mathcal{U}_{R_{i-1}W_i}}{\mathcal{U}_{R_{i-1}R_i}} &< \frac{1}{0.9}\times 0.1 \approx 0.11\end{aligned} \tag{21}$$

So we obtain

$$\begin{aligned}\mathscr{F}_i &< \frac{\mathcal{U}_{R_{i-1}R_i}}{\mathcal{U}_{R_{i-1}W_i}}\left(1+3\times 0.11\left(1+3\times 0.11\right)\right)\\ &< \frac{1}{0.9}\frac{\overline{k}_{R_{i-1}R_i}}{\overline{k}_{R_{i-1}W_i}}\left(1+\frac{\overline{r}_{R_{i-1}W_i}}{\overline{k}_{W_iR_{i+1}}}\right)\times 1.44\\ &\approx 1.60\frac{\overline{k}_{R_{i-1}R_i}}{\overline{k}_{R_{i-1}W_i}}\left(1+\frac{\overline{r}_{R_{i-1}W_i}}{\overline{k}_{W_iR_{i+1}}}\right)\end{aligned} \tag{22}$$

Eq.(19)(22) give the lower bound and upper bound of $\mathscr{F}_i$

$$\begin{aligned}0.9F_i < \mathscr{F}_i &< 1.60F_i\\ F_i \equiv \frac{\overline{k}_{R_{i-1}R_i}}{\overline{k}_{R_{i-1}W_i}}&\left(1+\frac{\overline{r}_{R_{i-1}W_i}}{\overline{k}_{W_iR_{i+1}}}\right)\end{aligned} \tag{23}$$

We define the relative deviation $|F_i - \mathscr{F}_i|/\min(F_i, \mathscr{F}_i)$ to characterize the accuracy of the above approximations. Here $\min(a, b)$ means the smaller one of $a, b$. We use similar defination to characterize the accuracy of any approximate expression throughout this paper. It can be seen that $F_i$ deviates less than 60% from the precise $\mathscr{F}_i$, meaning it's a good approximation.

For exonuclease-deficient DNAP, we denote $\mathscr{F}_i$ as the initial discrimination $\mathscr{F}_{ini,i}$ (*i.e.*, $\mathscr{F}_i$ given by Eq.(10), taking $\overline{r}=0$). In such cases, $\mathcal{U}_{\alpha_{i-1}\alpha_i}=\overline{k}_{\alpha_{i-1}\alpha_i}$. Eq.(19) gives the lower bound of $\mathscr{F}_{ini,i}$

$$\mathscr{F}_{ini,i} > \frac{\mathcal{U}_{R_{i-1}R_i}}{\mathcal{U}_{R_{i-1}W_i}} = \frac{\overline{k}_{R_{i-1}R_i}}{\overline{k}_{R_{i-1}W_i}} \tag{24}$$

Eq.(20) gives the upper bound of $\mathscr{F}_{ini,i}$

$$\mathscr{F}_{ini,i} < \frac{\overline{k}_{R_{i-1}R_i}}{\overline{k}_{R_{i-1}W_i}}\left(1+3\times 0.1\left(1+3\times 0.1\right)\right) = 1.39\frac{\overline{k}_{R_{i-1}R_i}}{\overline{k}_{R_{i-1}W_i}} \tag{25}$$

Combining Eq.(24)(25), we get

$$\begin{aligned} F_i^{pol} &< \mathscr{F}_{ini,i} < 1.39F_i^{pol} \\ F_i^{pol} &\equiv \frac{\overline{k}_{R_{i-1}R_i}}{\overline{k}_{R_{i-1}W_i}} \end{aligned} \tag{26}$$

$F_i^{pol}$ deviates less than 39% from the precise $\mathscr{F}_{ini,i}$.

Based on the above $\mathscr{F}_i$ and $\mathscr{F}_{ini,i}$, the proofreading efficiency can now be defined as $\mathscr{F}_{pro,i}\equiv\mathscr{F}_i/\mathscr{F}_{ini,i}$. Correspondingly, we also define $F_i^{exo}\equiv F_i/F_i^{pol}=1+\overline{r}_{R_{i-1}W_i}/\overline{k}_{W_iR_{i+1}}$, and have

$$\begin{aligned} \mathscr{F}_{pro,i} &= \frac{\mathscr{F}_i/F_i}{\mathscr{F}_{ini,i}/F_i^{pol}}F_i^{exo} \\ 0.65F_i^{exo} &< \mathscr{F}_{pro,i} < 1.60F_i^{exo} \end{aligned} \tag{27}$$

So $F_i^{exo}$ deviates less than 60% from the precise $\mathscr{F}_{pro,i}$, *i.e.*, $F_i^{exo}$ is a good estimate of $\mathscr{F}_{pro,i}$.

It should be noted that $\overline{k}_{RR}$ can be much larger (say, more than 100 times larger) than $\overline{k}_{RW},\overline{r}_{WR},\overline{r}_{RR}$ for real DNA replications. In such cases, one can get

$$\begin{aligned} 0.99F_i &< \mathscr{F}_i < 1.04F_i \\ F_i^{pol} &< \mathscr{F}_{ini,i} < 1.03F_i^{pol} \\ 0.96F_i^{exo} &< \mathscr{F}_{pro,i} < 1.04F_i^{exo} \end{aligned} \tag{28}$$

$F_i, F_i^{pol}, F_i^{exo}$ deviate less than 4% from their precise counterparts respectively. So we claimed in the main text that the relative deviations of these approximate expressions are about 10%. Fig.5(b) in Ref.[1] gives numerical examples to support this conclusion.

In summary, we list the approximate expressions of DNAP fidelity under bio-relevant conditions, as below

$$\mathscr{F}_i\equiv\mathscr{F}_{ini,i}\cdot\mathscr{F}_{pro,i},\qquad \mathscr{F}_{ini,i}\approx F_i^{pol}\equiv\frac{\overline{k}^{X_{i-1}X_i}_{R_{i-1}R_i}}{\overline{k}^{X_{i-1}X_i}_{R_{i-1}W_i}},\qquad \mathscr{F}_{pro,i}\approx F_i^{exo}\equiv 1+\frac{\overline{r}^{X_{i-1}X_i}_{R_{i-1}W_i}}{\overline{k}^{X_iX_{i+1}}_{W_iR_{i+1}}} \tag{29}$$

The bio-relevant conditions (a)(b)(c) deserve more explanations. The conditions (a) and (b) are based on experimental data about real DNAPs, *i.e.*, $k_{RR}$ is always more than three orders of magnitude larger than $k_{RW}$, and $k_{WW}$ is never measured in experiments, as will explained in more details in Sec.II. The condition (c) is consistent with the premise of the conventional kinetic assays (the steady-state assay and the transient-state assay). As mentioned in Introduction of the main text, the total fidelity defined by the conventional kinetic assays is assumed to consist of two factors $f_{ini}$ (initial discrimination) and $f_{pro}$ (proofreading efficiency). $f_{pro}$ is defined as $f_{pro,i} \equiv P_{el,R_i}/P_{el,W_i}$, in which $P_{el,R_i} \equiv k_{el,R_iR_{i+1}}/(k_{el,R_iR_{i+1}} + k_{ex,R_{i-1}R_i})$ and $P_{el,W_i} \equiv k_{el,W_iR_{i+1}}/(k_{el,W_iR_{i+1}} + k_{ex,R_{i-1}W_i})$. $k_{el}$ is the rate of elongation to the next site, $k_{ex}$ is the rate of excision of the terminal nucleotide, and $P_{el}$ is the elongation probability. $k_{el,RR}, k_{ex,RR}, k_{el,WR}, k_{ex,RW}$ are similar to $\overline{k}_{RR}, \overline{r}_{RR}, \overline{k}_{WR}, \overline{r}_{RW}$ in the FP method. It is always assumed in the kinetic assays that $P_{el,R_i} \approx 100\%$, meaning $k_{el,R_iR_{i+1}} \gg k_{ex,R_{i-1}R_i}$ which is consistent with $\overline{k}_{RR} \gg \overline{r}_{RR}$ in the condition (c). So we have $f_{pro,i} \approx 1/P_{el,W_i} = 1 + k_{ex,R_{i-1}W_i}/k_{el,W_iR_{i+1}}$, which is similar to $F_{pro}$. On the other hand, $k_{el,W_iR_{i+1}}$, by definition, should be more properly understood as the ultimate elongation rate in the sense that the elongated terminal ($R_{i+1}$) is no longer excised. In other words, the further elongation probability $P'_{el,R_{i+1}} \equiv k_{el,R_{i+1}R_{i+2}}/(k_{el,R_{i+1}R_{i+2}} + k_{ex,W_iR_{i+1}})$ should be almost 100%, meaning $k_{el,R_{i+1}R_{i+2}} \gg k_{ex,W_iR_{i+1}}$ which is consistent with $\overline{k}_{RR} \gg \overline{r}_{WR}$ in condition (c). Or else, $k_{el,W_i}$ should depend on farther neighors like $R_{i+2}, R_{i+3}, ...$, which is never considered in previous kinetic assays. So in this paper, we restrict our discussions in accord with the logic of conventional kinetic assays, assuming $\overline{r}^{X_iX_{i+1}}_{W_iR_{i+1}} \ll \overline{k}^{X_{i+1}X_{i+2}}_{R_{i+1}R_{i+2}}$ by default. Under the bio-relevant conditions, we can safely use Eq.(29) in this paper and compare it with the results given by kinetic assays.

It should be pointed out that $\overline{k}_{RR} \gg \overline{r}_{WR}$ is not necessary in the above FP analysis. Discarding this condition, one can get

$$\mathscr{F}_i \equiv \mathscr{F}_{ini,i} \cdot \mathscr{F}_{pro,i}, \qquad \mathscr{F}_{ini,i} \approx F_i^{pol} \equiv \frac{\overline{k}^{X_{i-1}X_i}_{R_{i-1}R_i}}{\overline{k}^{X_{i-1}X_i}_{R_{i-1}W_i}}, \qquad \mathscr{F}_{pro,i} \approx F_i^{exo} \equiv 1 + \frac{\overline{r}^{X_{i-1}X_i}_{R_{i-1}W_i}}{\overline{k}^{X_iX_{i+1}}_{W_iR_{i+1}}}\left(1 + \frac{\overline{r}^{X_iX_{i+1}}_{W_iR_{i+1}}}{\overline{k}^{X_{i+1}X_{i+2}}_{R_{i+1}R_{i+2}}}\right) \tag{30}$$

The factor $\overline{r}^{X_iX_{i+1}}_{W_iR_{i+1}}/\overline{k}^{X_{i+1}X_{i+2}}_{R_{i+1}R_{i+2}}$ can also be measured by kinetic assays in principle.

#### B. Direct competition assays: an example of FP method

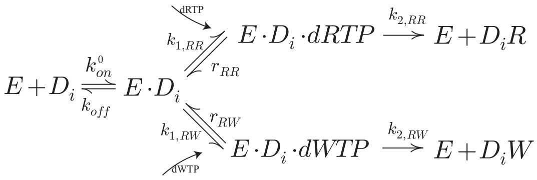

Fig.S 2: The three-step reaction scheme of the competitive incorporation of dRTP and dWTP. $E$: $exo^-$-DNAP. $D_i$: the primer-template duplex with the matched(R) terminal at site $i$. For brevity, the subscript $i$ in each rate constant is omitted.

Fig.S2 shows a three-step kinetic model of the competitive incorporation of a single dRTP or dWTP (at site $i+1$) to the matched (R) terminal (at site $i$). Since W occurs with much smaller probability than R in the primer, only the incorporation of a dRTP or a dWTP to the matched terminal contributes to the fidelity. So the true fidelity can be estimated by the ratio of the final product $D_iR$ ($R_iR$) to $D_iW$ ($R_iW$) when the substrate DNA are totally consumed, *i.e.*,

$$\mathscr{F} \approx F = \frac{[D_i R]}{[D_i W]} \tag{31}$$

Kinetic equations for this model are given below ,

$$\begin{aligned} \frac{d}{dt}[D_i\alpha] &= k_{2,R\alpha}[E \cdot D_i \cdot d\alpha TP] \\ \frac{d}{dt}[E \cdot D_i \cdot d\alpha TP] &= k^0_{1,R\alpha}[E \cdot D_i][d\alpha TP] \\ &- (k_{2,R\alpha} + r_{R\alpha})[E \cdot D_i \cdot d\alpha TP] \end{aligned} \tag{32}$$

here $\alpha$ =R,W. The dNTP binding rate is denoted as $k_{1,R\alpha} = k^0_{1,R\alpha}[d\alpha TP]$. The basic idea of FP method is not to directly solve the kinetic equations rigorously (*e.g.* in the transient-state analysis) or approximately by imposing extra assumptions (*e.g.* the steady-state assumption) . Instead, the two equations are integrated to give the products at time $t$,

$$\begin{aligned} [D_i\alpha](t) &= \frac{k^0_{1,R\alpha}k_{2,R\alpha}}{k_{2,R\alpha} + r_{R\alpha}} \int_0^t ([E \cdot D_i](\tau)[d\alpha TP](\tau))d\tau \\ &- \frac{k_{2,R\alpha}}{k_{2,R\alpha} + r_{R\alpha}}[E \cdot D_i \cdot d\alpha TP](t) \end{aligned} \tag{33}$$

The second term approaches to zero with $t$ increases to infinity. dNTP is usually in large excess to template DNA either *in vivo* or *in vitro*, so [dNTP] remains approximately a constant during the reaction. Then the fidelity is simply given by

$$\begin{aligned} F &= \overline{k}_{RR}\Big/\overline{k}_{RW} \\ \overline{k}_{RR} &= \frac{k^0_{1,RR}k_{2,RR}}{k_{2,RR} + r_{RR}}[\text{dRTP}] \\ \overline{k}_{RW} &= \frac{k^0_{1,RW}k_{2,RW}}{k_{2,RW} + r_{RW}}[\text{dWTP}] \end{aligned} \tag{34}$$

$F$ is exactly the initial discrimination defined by Eq.(29) with the two effective incorporation rates $\overline{k}_{RR}$ and $\overline{k}_{RW}$.

In practice, when the reaction time $t$ is large enough for sufficient product accumulation (*i.e.*, the second term on the right side of Eq.(33) is far smaller than the first term), the measured $[D_iR](t)/[D_iW](t)$ becomes nearly time-invariant, and thus it is a good measure of $F$. In the direct competition assay conducted by Bertram *et.al*[6], the incorporation reaction was terminated when about half of the substrate DNA were reacted. This termination criteria *per se* does not meet the above requirement. Other evidences should be considered. For instance, $[D_iR](t)/[D_iW](t)$ is proportional to [dRTP]/[dWTP] if the reaction time $t$ is large, so one can decide whether $t$ is sufficient large by examining whether $[D_iR](t)$[dWTP]$/[D_iW](t)$[dRTP] becomes nearly a constant when [dWTP] or [dRTP] is changed. Combined with these evidences, Bertram *et.al* were able to show that $[D_iR](t)/[D_iW](t)$ measured under their termination condition is really a good measure of the true fidelity.

#### C. Reduction of the multi-step reaction scheme of DNAP without exonuclease domain

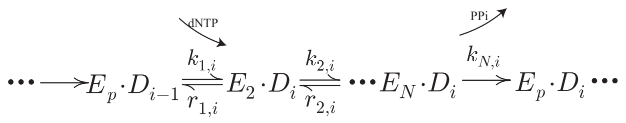


Fig.S3: The multi-step incorporation scheme of DNAP without exonuclease domain. The enzyme-substrate complex $(E \cdot D)$ goes through $N$ states (indicated by subscripts $1, ..., N$) to successfully incorporate a single nucleotide(indicated by subscript $i$). To simplify the notation, the superscripts indicating the template nucleotide $X_i$ and the subscripts indicating the primer nucleotide $\alpha_i$ are omitted. This rule applies to other figures in this paper, unless otherwise specified.

The FP method can also be applied to more complex reaction schemes such as the multi-step scheme in Fig.S3. These schemes can be reduced (mapped) uniquely to the minimal scheme Fig.S1 and hence the true fidelity $\mathscr{F}$ can be calculated by Eq.(29).

Similar to Sec.I A, one can write the master equations for all the states in the reaction scheme of replicating a template $X_1...X_L$ and integrate these equations to give the sequence distribution of the final products $P^{X_1...X_L}_{\alpha_1...\alpha_L}(t \to \infty)$. Here we only show the integrated equations for the variables $\Gamma^{X_1...X_i...X_L}_{\alpha_1...\alpha_i,2}, ..., \Gamma^{X_1...X_i...X_L}_{\alpha_1...\alpha_i,N}, \Gamma^{X_1...X_i...X_L}_{\alpha_1...\alpha_i,p}$ ($\Gamma^{X_1...X_i...X_L}_{\alpha_1,...\alpha_i,x}$ corresponds to $E_x \cdot D_i$ in Fig.S3, $x = 2...N, p$)

$$
\begin{aligned}
0 &= k^{X_{i-1}X_i}_{\alpha_{i-1}\alpha_i,1}\Gamma^{X_1...X_{i-1}...X_L}_{\alpha_1...\alpha_{i-1},p} + r^{X_{i-1}X_i}_{\alpha_{i-1}\alpha_i,2}\Gamma^{X_1...X_i...X_L}_{\alpha_1...\alpha_i,3} - \left(k^{X_{i-1}X_i}_{\alpha_{i-1}\alpha_i,2} + r^{X_{i-1}X_i}_{\alpha_{i-1}\alpha_i,1}\right)\Gamma^{X_1...X_i...X_L}_{\alpha_1...\alpha_i,2} \\
0 &= k^{X_{i-1}X_i}_{\alpha_{i-1}\alpha_i,j-1}\Gamma^{X_1...X_i...X_L}_{\alpha_1...\alpha_i,j-1} + r^{X_{i-1}X_i}_{\alpha_{i-1}\alpha_i,j}\Gamma^{X_1...X_i...X_L}_{\alpha_1...\alpha_i,j+1} - \left(k^{X_{i-1}X_i}_{\alpha_{i-1}\alpha_i,j} + r^{X_{i-1}X_i}_{\alpha_{i-1}\alpha_i,j-1}\right)\Gamma^{X_1...X_i...X_L}_{\alpha_1...\alpha_i,j}, 2 < j < N \\
0 &= k^{X_{i-1}X_i}_{\alpha_{i-1}\alpha_i,N-1}\Gamma^{X_1...X_i...X_L}_{\alpha_1...\alpha_i,N-1} - \left(k^{X_{i-1}X_i}_{\alpha_{i-1}\alpha_i,N} + r^{X_{i-1}X_i}_{\alpha_{i-1}\alpha_i,N-1}\right)\Gamma^{X_1...X_i...X_L}_{\alpha_1...\alpha_i,N}
\end{aligned} \tag{35}
$$

and

$$
0 = k^{X_{i-1}X_i}_{\alpha_{i-1}\alpha_i,N}\Gamma^{X_1...X_i...X_L}_{\alpha_1...\alpha_i,N} + \sum_{\alpha_{i+1}} r^{X_iX_{i+1}}_{\alpha_i\alpha_{i+1},1}\Gamma^{X_1...X_{i+1}...X_L}_{\alpha_1...\alpha_{i+1},2} - \sum_{\alpha_{i+1}} k^{X_iX_{i+1}}_{\alpha_i\alpha_{i+1},1}\Gamma^{X_1...X_i...X_L}_{\alpha_1...\alpha_i,p} \tag{36}
$$

Here $k^{X_{i-1}X_i}_{\alpha_{i-1}\alpha_i,x}, r^{X_{i-1}X_i}_{\alpha_{i-1}\alpha_i,x}$ correspond to $k_{x,i}, r_{x,i}$ in Fig.S3. According to Eq.(35), one can eliminate the variables $\Gamma^{X_1...X_i...X_L}_{\alpha_1...\alpha_i,2}, ..., \Gamma^{X_1...X_i...X_L}_{\alpha_1...\alpha_i,N}$ by expressing them as functions of $\Gamma^{X_1...X_{i-1}...X_L}_{\alpha_1...\alpha_{i-1},p}$. Similarly, $\Gamma^{X_1...X_{i+1}...X_L}_{\alpha_1...\alpha_{i+1},2} ... \Gamma^{X_1...X_{i+1}...X_L}_{\alpha_1...\alpha_{i+1},N}$, and $\Gamma^{X_1...X_{i+1}...X_L}_{\alpha_1...\alpha_{i+1},2}$ can also be solved as a function of $\Gamma^{X_1...X_i...X_L}_{\alpha_1...\alpha_i,p}$. So Eq.(36) can be rewritten as

$$
0 = k^{*X_{i-1}X_i}_{\alpha_{i-1}\alpha_i}\Gamma^{X_1...X_{i-1}...X_L}_{\alpha_1...\alpha_{i-1},p} - \sum_{\alpha_{i+1}} k^{*X_iX_{i+1}}_{\alpha_i\alpha_{i+1}}\Gamma^{X_1...X_i...X_L}_{\alpha_1...\alpha_i,p} \tag{37}
$$

Here $k^*$ is

$$
\begin{aligned}
k^* &= k'_{N-1}k_N/(r'_{N-1}+k_N) \\
k'_j &= k'_{j-1}k_j/(r'_{j-1}+k_j) \\
r'_j &= r'_{j-1}r_j/(r'_{j-1}+k_j) \qquad (j=N-1,...,3) \\
k'_2 &= k_1k_2/(r_1+k_2) \\
r'_2 &= r_1r_2/(r_1+k_2)
\end{aligned} \tag{38}
$$

Note that all the kinetic rates in Eq.(38) have the same superscripts and subscripts, *i.e.*, ${k^*}^{X_{i-1}X_i}_{\alpha_{i-1}\alpha_i} = \left(k'_{N-1}k_N/(r'_{N-1}+k_N)\right)^{X_{i-1}X_i}_{\alpha_{i-1}\alpha_i}$.

Comparing Eq.(37) with Eq.(2), one can identify $k^*$ as the effective incorporation rate (${k^*}^{X_{i-1}X_i}_{\alpha_{i-1}\alpha_i} = \overline{k}^{X_{i-1}X_i}_{\alpha_{i-1}\alpha_i}$, if $\overline{r}=0$). No approximations such as the steady-state assumption or the quasi-equilibrium assumption are needed in the reduction procedure. It should be noted that the above reduction is unique in the sense that other reduction procedure eliminating $\Gamma^{X_1...X_i...X_L}_{\alpha_1...\alpha_i,p}(i=1,...,L)$ will lead to equations much different from Eq.(2). For example, one can eliminate all the variables except $\Gamma^{X_1...X_i...X_L}_{\alpha_1...\alpha_i,N}$. The reduced integrated equation for $\Gamma^{X_1...X_i...X_L}_{\alpha_1...\alpha_i,N}$ is written as

$$
\begin{aligned}
0 \;=\; & \frac{{k'}^{X_{i-1}X_i}_{\alpha_{i-1}\alpha_i,N-1}k^{X_{i-2}X_{i-1}}_{\alpha_{i-2}\alpha_{i-1},N}}{\sum_{\alpha_i}{k'}^{X_{i-1}X_i}_{\alpha_{i-1}\alpha_i,N-1}}\Gamma^{X_1...X_{i-1}...X_L}_{\alpha_1...\alpha_{i-1},N} - \left({r'}^{X_{i-1}X_i}_{\alpha_{i-1}\alpha_i,N-1} + k^{X_{i-1}X_i}_{\alpha_{i-1}\alpha_i,N}\right)\Gamma^{X_1...X_i...X_L}_{\alpha_1...\alpha_i,N} \\
+ \; & \frac{{k'}^{X_{i-1}X_i}_{\alpha_{i-1}\alpha_i,N-1}\left[\sum_{\alpha_i}\left({r'}^{X_{i-1}X_i}_{\alpha_{i-1}\alpha_i,N-1}\Gamma^{X_1...X_i...X_L}_{\alpha_1...\alpha_i,N}\right)\right]}{\sum_{\alpha_i}{k'}^{X_{i-1}X_i}_{\alpha_{i-1}\alpha_i,N-1}}
\end{aligned} \tag{39}
$$

$k'_{N-1}, r'_{N-1}$ is given by Eq.(38). Eq.(39) displays a form totally different from Eq.(37) and Eq.(2), in which no effective incorporation rates can be properly defined. In such cases, Eq.(29) is invalid to calculate the fidelity.

The logic of the reduction procedure applies to all multi-step reaction schemes in this paper.

#### D. Reduction of the multi-step reaction scheme of *exo*$^+$-DNAP

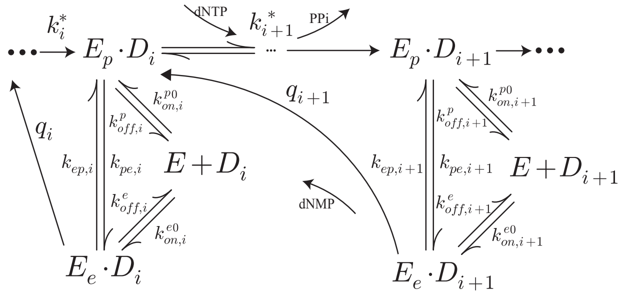


Fig.S4: The multi-step reaction scheme of *exo*$^+$-DNAP.

The complete reaction scheme of $exo^+$ DNAP is shown in Fig.S4, considering additionally the binding and unbinding between the enzyme and the substrate. The integrated master equations are

$$\begin{aligned}
0 &= k_i^*\Gamma_{p,i-1} + k_{ep,i}\Gamma_{e,i} + k_{on,i}^p\Gamma_{0,i} + \sum_{\alpha_{i+1}} q_{i+1}\Gamma_{e,i+1} - \left(k_{pe,i} + k_{off,i}^p + \sum_{\alpha_{i+1}} k_{i+1}^*\right)\Gamma_{p,i} \\
0 &= k_{off,i}^p\Gamma_{p,i} + k_{off,i}^e\Gamma_{e,i} - \left(k_{on,i}^p + k_{on,i}^e\right)\Gamma_{0,i} \\
0 &= k_{pe,i}\Gamma_{p,i} + k_{on,i}^e\Gamma_{0,i} - \left(k_{ep,i} + k_{off,i}^e + q_i\right)\Gamma_{e,i} \ (i = 2, 3, ...L-1)
\end{aligned} \tag{40}$$

Here the notations are simplified, *e.g*, $\Gamma_{x,i} \equiv \Gamma^{X_1...X_i...X_L}_{\alpha_1...\alpha_i,x}$ and $k_{x,i} \equiv k^{X_{i-1}X_i}_{\alpha_{i-1}\alpha_i,x}$. $\Gamma_{0,i}$ corresponds to the unbinding state $E + D_i$. $k_{on}^p = k_{on}^{p0}[E]$,$k_{on}^e = k_{on}^{e0}[E]$.

Eq.(40) is not the original integrated master equations. For brevity, we have omitted the intermediate states ($E_2, ...E_N$ in Fig.S3) and the corresponding integrated master equations. As shown in the above section (Sec.I C), eliminating these intermediate states gives Eq.(37) with $k^*$ which can be regarded as apparent rate constants in the FP analysis. So we can write directly the simplified version Eq.(40). We will use this convention for FP analysis elsewhere in this paper.

By eliminating $\Gamma_e$ and $\Gamma_0$ and retaining $\Gamma_p$ (the state for dNTP binding), one can reduce the above equations to Eq.(2)

$$0 = \overline{k}_i\Gamma_{p,i-1} + \sum_{\alpha_{i+1}} \overline{r}_{i+1}\Gamma_{p,i+1} - \left(\overline{r}_i + \sum_{\alpha_{i+1}} \overline{k}_{i+1}\right)\Gamma_{p,i} \tag{41}$$

with the effective rates

$$\begin{aligned}
\overline{k} &= k^* \quad , \quad \overline{r} = \frac{\widetilde{k}_{pe}q}{\widetilde{k}_{ep} + q} \\
\widetilde{k}_{pe} &= k_{pe} + k_{p\to e} \quad , \quad \widetilde{k}_{ep} = k_{ep} + k_{e\to p} \\
k_{p\to e} &= \frac{k_{off}^p k_{on}^e}{k_{on}^p + k_{on}^e} \quad , \quad k_{e\to p} = \frac{k_{off}^e k_{on}^p}{k_{on}^p + k_{on}^e}
\end{aligned} \tag{42}$$

This reduction is unique in the sense that other reduction procedures eliminating $\Gamma_p$ always lead to much different equations in which no effective rates can be identified.

### II. THE STEADY-STATE ASSAY

The steady-state assay is a standard method to analyze the catalytical capability of enzymes in biochemistry. The steady-state condition in experiment is usually established by two requirements, *i.e.* the substrate is in large excess to the enzyme, and the enzyme can dissociate quickly from the product once a single turnover is finished to resume its catalysis. The last dissociation step is reasonably assumed irreversible, since the enzyme will much unlikely rebind to the same substrate molecule after dissociation because the substrate is in large excess to the enzyme. Under such conditions, the amount of any intermediate product can be regarded approximately as constant (*i.e.* in steady state) in the initial stage of the reaction and thus the amount of the final product increases linearly with time. This initial growth velocity is defined as the steady-state turnover rate which is often used to characterize the catalytical capability of the enzyme. There have been intensive discussions on the theoretical foundation of the steady-state assays [3]. Here we briefly review the steady-state analysis of the reaction schemes discussed in this paper.

#### A. The specificity constant of DNAP without exonuclease domain

The multi-step incorporation process in Fig.S5 is used to illustrate the general idea of the steady-state assays of DNAP fidelity.

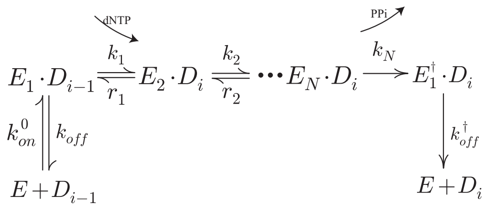


Fig.S 5: The reaction scheme of the steady-state assay to measure the specificity constant of the nucleotide incorporation reaction of DNAP without exonuclease domain.

In the steady-state experiment, the substrate DNA concentration is much higher than the enzyme concentration. During the steady growth phase of the product, the normalized turnover velocity per enzyme can be written as

$$v_{s\cdot s}^{pol} = \frac{k_{off}^{\dagger}[E_1^{\dagger}]}{[E_{tot}]} \tag{43}$$

Here $[E_1^\dagger]$ represents the concentration of $E_1^\dagger \cdot D_i$, and $[E_{tot}]$ represents the total enzyme concentration. Both the experimental results and the theoretical analysis show that the velocity obeys the Michaelis-Menten equation

$$v_{s\cdot s}^{pol} = \frac{k_{cat}[\text{dNTP}]}{K_m + [\text{dNTP}]} \tag{44}$$

The quasi-first order rate constant $k_{cat}/K_m$ is called the specificity constant, which can be obtained by directly solving the following kinetic equations under the steady-state assumptions.

$$\begin{aligned}
\frac{d}{dt}[E_0] = 0 &= k^{\dagger}_{off}[E^{\dagger}_1] + k_{off}[E_1] - k_{on}[E_0] \\
\frac{d}{dt}[E_1] = 0 &= k_{on}[E_0] + r_1[E_2] - (k_1 + k_{off})\,[E_1] \\
\frac{d}{dt}[E_2] = 0 &= k_1[E_1] + r_2[E_3] - (k_2 + r_1)\,[E_2] \\
&\ \ \ldots \\
\frac{d}{dt}[E_j] = 0 &= k_{j-1}[E_{j-1}] + r_j[E_{j+1}] - (k_j + r_{j-1})\,[E_j] \\
&\ \ \ldots \\
\frac{d}{dt}[E_N] = 0 &= k_{N-1}[E_{N-1}] - (k_N + r_{N-1})\,[E_N] \\
\frac{d}{dt}[E^{\dagger}_1] = 0 &= k_N[E_N] - k^{\dagger}_{off}[E^{\dagger}_1]
\end{aligned} \qquad (45)$$

$[E_0]$ represents the concentration of free enzyme. $k_1 = k^0_1[\mathrm{dNTP}]$ represents the dNTP binding rate. $k_{on} = k^0_{on}[\mathrm{DNA}]$ represents the enzyme-substrate binding rate. Solving these equations, one can get

$$\left(\frac{k_{cat}[\mathrm{dNTP}]}{K_m}\right)_{\alpha_{i-1}\alpha_i} = \frac{k^*_{\alpha_{i-1}\alpha_i}}{(K_{s\cdot s})_{\alpha_{i-2}\alpha_{i-1}}}, \quad K_{s\cdot s} = 1 + \frac{k_{off}}{k_{on}} \qquad (46)$$

Here $k^*$ is the effective incorporation rate defined by Eq.(38), $k^* = k^{*0}[\mathrm{dNTP}]$. If the DNA concentration [DNA] is set large enough in the steady-state assay, *i.e.*, $k_{on} \equiv k^0_{on}[\mathrm{DNA}] \gg k_{off}$, we have $K_{s\cdot s} \approx 1$. So the specificity constant is a quite good approximation of $k^*$.

Now we give some detailed explanations on the validity of the bio-relevent conditions given in preceding section. Since $K_{s\cdot s}$ is independent on the incoming dNTP($\alpha_i$), we have $(k_{cat}[\mathrm{dNTP}]/K_m)_{RR}/(k_{cat}[\mathrm{dNTP}]/K_m)_{RW} = k^*_{RR}/k^*_{RW}$. The experimental data shows that $(k_{cat}[\mathrm{dNTP}]/K_m)_{RR}/(k_{cat}[\mathrm{dNTP}]/K_m)_{RW} \gg 1$ (more than three orders of magnitude larger. See the examples in Table I in the main text), so $k^*_{RR}/k^*_{RW} \gg 1$. On the other hand, the FP method gives $\overline{k} = k^*$. So the bio-relevant condition (a) is reasonable. Similarly, the steady-state assays can measure $(k_{cat}[\mathrm{dNTP}]/K_m)_{WR}$ but failed to measure $(k_{cat}[\mathrm{dNTP}]/K_m)_{WW}$, meaning that $(k_{cat}[\mathrm{dNTP}]/K_m)_{WW}$ is too small to be measured. Since $(k_{cat}[\mathrm{dNTP}]/K_m)_{WR}/(k_{cat}[\mathrm{dNTP}]/K_m)_{WW} = k^*_{WR}/k^*_{WW}$, we can assume that $k^*_{WW}$ is arbitrarily smaller than $k^*_{WR}$, *i.e.*, $k^*_{WW} \approx 0s^{-1}$ ($\overline{k}_{WW} \approx 0s^{-1}$). Another evdience to show $\overline{k}_{WW} \approx 0s^{-1}$ is given by the steady-state assay to measure $(k_{cat}[\mathrm{dNTP}]/K_m)_{RW}$. For some designed DNA templates and specifically chosen dNTP, there can be $RW$ and $RWW$ in the product in principle. However, the product $RW$ can be observed, while the product $RWW$ has never been detected. This means the incorporation of a dWTP to the mismatched terminal is extremely slow, *i.e.* $k^*_{WW} \approx 0s^{-1}$. So condition (b) $\overline{k}_{WW} \approx 0s^{-1}$ is reasonable.

#### B. The specificity constant of DNAP with deficient exonuclease domain

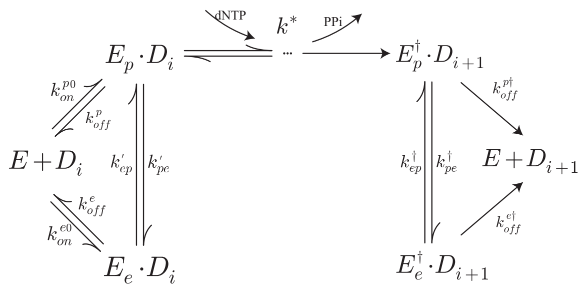


Fig.S 6: The reaction scheme of the steady-state assay to measure the specificity constant of the nucleotide incorporation reaction of DNAP with deficient exonuclease domain.

When the DNAP contains a deficient exonuclease domain, the nucleotide incorporation scheme is more complicated, as shown in Fig.S6. The primer terminal may shuttle between *Pol* and *Exo*, and the transfer rate $k_{pe}^{'}$ and $k_{ep}^{'}$ are different from $k_{pe}$ and $k_{ep}$ of the wild type $exo^+$-DNAP. The analysis in Sec.II A can be directly applied to this scheme. The normalized turnover velocity per enzyme is

$$v_{s \cdot s}^{pol} \; = \; \frac{k_{off}^{p\dagger}[E_p^\dagger] + k_{off}^{e\dagger}[E_e^\dagger]}{[E_{tot}]} \tag{47}$$

The corresponding steady-state equations are

$$\begin{aligned}
\frac{d}{dt}[E_0] = 0 \; &= \; k_{off}^{p\dagger}[E_p^\dagger] + k_{off}^{e\dagger}[E_e^\dagger] + k_{off}^{p}[E_p] + k_{off}^{e}[E_e] - \left(k_{on}^{p} + k_{on}^{e}\right)[E_0] \\
\frac{d}{dt}[E_p] = 0 \; &= \; k_{on}^{p}[E_0] + k_{ep}^{'}[E_e] - \left(k_{off}^{p} + k_{pe}^{'} + k^*\right)[E_p] \\
\frac{d}{dt}[E_e] = 0 \; &= \; k_{on}^{e}[E_0] + k_{pe}^{'}[E_p] - \left(k_{off}^{e} + k_{ep}^{'}\right)[E_e] \\
\frac{d}{dt}[E_p^\dagger] = 0 \; &= \; k^*[E_p] + k_{ep}^{\dagger}[E_e^\dagger] - \left(k_{off}^{p\dagger} + k_{pe}^{\dagger}\right)[E_p^\dagger] \\
\frac{d}{dt}[E_e^\dagger] = 0 \; &= \; k_{pe}^{\dagger}[E_p^\dagger] - \left(k_{off}^{e\dagger} + k_{ep}^{\dagger}\right)[E_e^\dagger]
\end{aligned} \tag{48}$$

These equations are not the original steady-state kinetic equations. For simplicity, we have omitted the intermediate states in the dNTP incorporation process ($E_2, E_3 ... E_N$ in Fig.S5) and the corresponding steady-state kinetic equations which are exactly the same as given by Eq.(45). In fact, after eliminating these variables first by Eq.(45), we obtain the simplified Eq.(48) with $k^*$ defined by Eq.(38). This simplification convention will be used in steady-state analysis elsewhere in this paper.

Solving these equations, one can finally get

$$\left(\frac{k_{cat}[\text{dNTP}]}{K_m}\right)_{\alpha_{i-1}\alpha_i} = \frac{k^*_{\alpha_{i-1}\alpha_i}}{(K_{s\cdot s})_{\alpha_{i-2}\alpha_{i-1}}}, \quad K_{s\cdot s} = 1 + \frac{k_{pe}^{'}}{k_{ep}^{'}} + \frac{k_{off}^{p}}{k_{on}^{p}} \tag{49}$$

Since the DNA concentration [DNA] is always set large enough in the steady-state assay, *i.e.*, $k_{on} \equiv k_{on}^0[\text{DNA}] \gg k_{off}$, and since the mutant $exo^-$-DNAP has no exonuclease domain ( $k_{pe}^{'}/k_{ep}^{'}$ is absent) or it binds the DNA preferentially at the polymerase domain ( $k'_{ep} \gg k'_{pe}$. It can be more than two orders of magnitude larger when the difference of DNA binding free energy between the Pol and Exo is about a few $k_BT$ ), we have $K_{s\cdot s} \approx 1$. So the specificity constant is a quite good approximation of $k^*$.

For this reaction scheme, one can also get $(k_{cat}[\text{dNTP}]/K_m)_{RR}/(k_{cat}[\text{dNTP}]/K_m)_{RW} = k^*_{RR}/k^*_{RW}$ since $K_{s\cdot s}$ is independent on the incoming dNTP. Following the analysis in Sec.II A, one can also get $\overline{k}_{RR}/\overline{k}_{RW} \gg 1$ and $\overline{k}_{WW} \approx 0s^{-1}$ to validate the bio-relevant conditions (a) and (b).

It can be proven that Eq.(49) always holds (except that the explicit function $K_{s.s}$ may differ), no matter how complicated the scheme is: $k^*$ is exactly the effective incorporation rate, and $K_{s\cdot s}$ is a simple function of the equilibrium constants of all the steps before dNTP binding and thus independent on the incoming dNTP. So the equality $(k_{cat}[\text{dNTP}]/K_m)_{RR}/(k_{cat}[\text{dNTP}]/K_m)_{RW} = k^*_{RR}/k^*_{RW}$ holds universally. Following the same logic in Sec.II A, one can also validate the bio-relevant conditions (a) and (b).

#### C. The steady-state assay of the excision reaction of $exo^+$-DNAP

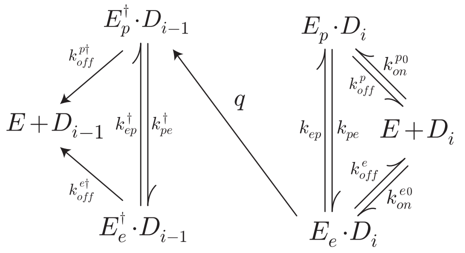


Fig.S7: The reaction scheme of the steady-state assay to measure the effective excision rate of $exo^+$-DNAP.

The steady-state assay has been employed to characterize the effective excision rate of $exo^+$-DNAP, *e.g.* by measuring the excision velocity [4] or the so-called specificity constant of the excision reaction [5]. Here we show it's not a proper way to get the correct excision rate. The excision velocity per enzyme measured by the steady-state assay is

$$v_{s\cdot s}^{exo} = \frac{k_{off}^{p\dagger}[E_p^\dagger] + k_{off}^{e\dagger}[E_e^\dagger]}{[E_{tot}]} \tag{50}$$

The corresponding steady-state equations are

$$\begin{aligned}
\frac{d}{dt}[E_0] = 0 &= k_{off}^{p\dagger}[E_p^\dagger] + k_{off}^{e\dagger}[E_e^\dagger] + k_{off}^{p}[E_p] + k_{off}^{e}[E_e] - (k_{on}^{p} + k_{on}^{e})[E_0] \\
\frac{d}{dt}[E_p] = 0 &= k_{on}^{p}[E_0] + k_{ep}[E_e] - \left(k_{off}^{p} + k_{pe}\right)[E_p] \\
\frac{d}{dt}[E_e] = 0 &= k_{on}^{e}[E_0] + k_{pe}[E_p] - \left(k_{off}^{e} + k_{ep} + q\right)[E_e] \\
\frac{d}{dt}[E_p^\dagger] = 0 &= q[E_e] + k_{ep}^\dagger[E_e^\dagger] - \left(k_{off}^{p\dagger} + k_{pe}^\dagger\right)[E_p^\dagger] \\
\frac{d}{dt}[E_e^\dagger] = 0 &= k_{pe}^\dagger[E_p^\dagger] - \left(k_{off}^{e\dagger} + k_{ep}^\dagger\right)[E_e^\dagger]
\end{aligned} \tag{51}$$

here $k_{on}^{p} = k_{on}^{p0}[\text{DNA}]$ and $k_{on}^{e} = k_{on}^{e0}[\text{DNA}]$ represent the enzyme-DNA binding rates (they are the binding rates per enzyme, noting that the substrate DNA is in large excess to DNAP). By solving these equations, one can obtain the excision velocity per enzyme in the form of Michaelis-Menten equation

$$v_{s\cdot s}^{exo} = \frac{k_{cat}[\text{DNA}]}{K_m + [\text{DNA}]} \tag{52}$$

The mathematic expression of $v_{s\cdot s}^{exo}$ is too complex to be written here, but it can be shown that the specificity constant of the excision reaction is given by

$$\left(\frac{k_{cat}}{K_m}\right)^{exo} = \frac{q(k_{pe}k_{on}^{p0}/(k_{pe} + k_{off}^{p}) + k_{on}^{e0})}{q + k_{ep}k_{off}^{p}/(k_{pe} + k_{off}^{p}) + k_{off}^{e}} \tag{53}$$

Obviously, $v_{s\cdot s}^{exo}$ and $(k_{cat}/K_m)^{exo}[\text{DNA}]$ are much different from the effective excision rate $\overline{r}$ defined in Eq.(42) and thus cannot be used to calculate $f_{pro}$.

### III. THE TRANSIENT-STATE ASSAY

Similar to the steady-state assay, the transient-state assay can also be employed to study the polymerase activity and exonuclease activity of DNAP separately. The transient-state assay often refers to two different methods, the single-turnover assay or the pre-steady-state assay.

In single-turnover assays, the enzyme is in large excess to the substrate, and the dissociation of the enzyme from the product is neglected. The time course of the product accumulation or the substrate consumption is monitored. The data is then fitted by exponential functions (single-exponential or multi-exponential) to give one or more exponents (*i.e.* the characteristic rates). This assay can be applied either to the polymerization process or to the excision process, and the obtained rates are used to define the initial discrimination and the proofreading efficiency.

In pre-steady-state assays, the enzyme concentration is moderately higher than the substrate concentration (say, two or three times larger). The time course of the product accumulation or the substrate consumption is monitored, which always displays two stages. The earlier stage can be fitted by exponential functions which give several characteristic rates, and the later stage can be fitted by linear functions which gives the steady-state velocity. For the earlier stage, it's reasonable to assume that all the substrates remain bound to the enzymes, and the later dissociation completes the catalytic turnover to establish the subsequent steady-state state. So, in the pre-steady-state stage, the system evolves according to the same kinetic equations as that in the single-turnover assay. Hence, the two methods give the same characteristic rates, and we can only discuss the single-turnover assay in this paper.

In single-turnover assays, we are concerned only about the smallest characteristic rate. For instance, in the initial discrimination assay, one can get the smallest rate $v_{t\cdot s}^{pol}$ for each dNTP concentration in a wide range, which is found to be well fitted by Michaelis-Menten-like equation (similar to Eq.(44) in the steady-state assay). In the following subsections, we will show theoretically why this is always observed and how it is used to define the initial discrimination, by using the matrix method to solve the kinetic equations to get the smallest characteristic rate. It will be proven that the true initial discrimination can be approximately estimated by the single-turnover assay. The matrix method is also used to discuss the theoretical foundation of the proofreading efficiency assay. To illustrate the basic logic of the matrix method, we first take the simplest reaction model (the two-step incorporation reaction) as an example.

#### A. Matrix method: applied to the two-step irreversible incorporation reaction

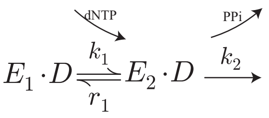


Fig.S8: The two-step reaction scheme of dNTP incorporation.

In the two-step incorporation reaction Fig.S8, we assume that the first step is dNTP binding where $k_1 = k_1^0$[dNTP] and the second step is the chemical step. The kinetic equations of the reaction are

$$\begin{aligned} \frac{d}{dt}[E_1 \cdot D] &= -k_1[E_1 \cdot D] + r_1[E_2 \cdot D] \\ \frac{d}{dt}[E_2 \cdot D] &= k_1[E_1 \cdot D] - (k_2 + r_1)\,[E_2 \cdot D] \end{aligned} \tag{54}$$

The solution of such equations is generally double-exponential

$$[D_{tot}(t)] = C_1 e^{-|\lambda_1|t} + C_2 e^{-|\lambda_2|t} \tag{55}$$

Here $[D_{tot}] = [E_1 \cdot D] + [E_2 \cdot D]$ is the total concentration of the substrate DNA. $\lambda_1, \lambda_2$ are the two characteristic rates which are given by the two eigenvalues (always $\leq 0$) of the coefficient matrix

$$\mathbf{M} = \begin{pmatrix} -k_1 & r_1 \\ k_1 & -(r_1 + k_2) \end{pmatrix} \tag{56}$$

In order to calculate the eigenvalues, one should solve the following equation

$$\begin{aligned} 0 = |\mathbf{M} - \lambda \mathbf{E}| &= \begin{vmatrix} -(k_1 + \lambda) & r_1 \\ k_1 & -(r_1 + k_2 + \lambda) \end{vmatrix} \\ &= A_2^{[2]}\lambda^2 + A_1^{[2]}\lambda + A_0^{[2]} \end{aligned} \tag{57}$$

Here $A_2^{[2]} = 1, A_1^{[2]} = k_1 + k_2 + r_1, A_0^{[2]} = k_1 k_2$. The superscript [2] indicates the reaction is two-stepped (similarly, the superscript $[N]$ for $N$-step reaction in later sections). The roots of Eq.(57) give the two eigenvalues. According to Vieta theorem $|\lambda_1| + |\lambda_2| = A_1^{[2]}/A_2^{[2]}$ and $|\lambda_1| \cdot |\lambda_2| = A_0^{[2]}/A_2^{[2]}$,

$$\frac{1}{|\lambda_1|} + \frac{1}{|\lambda_2|} = \frac{A_1^{[2]}}{A_0^{[2]}} \tag{58}$$

In the biochemical experiments, the time course of the product accumulation or the substrate consumption is monitored. For a given dNTP concentration, the time course is often well fitted by a single-exponential function. For the two-step reaction model, this implies two possibilities, *i.e.*, the two eigenvalues are almost equal ($|\lambda_1| \approx |\lambda_2|$) or the larger one (say, $|\lambda_2|$) may not be observed (*i.e.*, $|\lambda_1| \ll |\lambda_2|$, $\ll$ means at least one order of magnitude smaller). With the dNTP concentration changing in a wide range (*e.g.* from $1\mu M$ to $100\mu M$), the former may hold only in a narrow range, but the time course can always be well fitted by the single-exponential function in the entire range, so the latter is much more probable. With this preassumption, and note that $|\lambda_1| < 0.1\ |\lambda_2|$, one can get

$$\frac{1}{|\lambda_1|} < \frac{A_1^{[2]}}{A_0^{[2]}} < \frac{1.1}{|\lambda_1|} \tag{59}$$

*i.e.*,

$$|\lambda_1| \approx \frac{A_0^{[2]}}{A_1^{[2]}} = \frac{k_1 k_2}{k_1 + k_2 + r_1} = \frac{k_1^0 k_2 [\text{dNTP}]}{k_1^0 [\text{dNTP}] + k_2 + r_1} \tag{60}$$

Eq.(60) provides a good approximation of the precise $|\lambda_1|$, with the relative deviation less than 10%. Eq.(60) is also in the same form as the Michaelis-Menten equation. This explains why the observed $v_{t\cdot s}^{pol}$ (*i.e.*, $|\lambda_1|$) for various [dNTP] can be well fitted by the Michaelis-Menten equation $|\lambda_1| = k_{pol}[\text{dNTP}]/(K_d + [\text{dNTP}])$.

When extrapolating this function to [dNTP] $\approx 0$, we get the slope $k_{pol}/K_d \approx k_1^0 k_2/(k_2 + r_1)$. In Sec.I C we have obtained the effective incorporation rate $k^* = k_1 k_2/(k_2 + r_1)$ by the FP method for this two-step reaction model (see Eq.(38), $N = 2$ and $k_1 = k_1^0[\text{dNTP}]$). So here we obtain $k_{pol}[\text{dNTP}]/K_d \approx k^*$ . Notice that Eq.(60) always underestimates the real $|\lambda_1|$, so $k_{pol}[\text{dNTP}]/K_d$ obtained by data fitting always overestimates $k^*$ but with relative deviation no more than 10%. Hence, there is only less than 10% deviation of the operationally defined fidelity $(k_{pol}[\text{dNTP}]/K_d)_{RR}/(k_{pol}[\text{dNTP}]/K_d)_{RW}$ from the true fidelity $k^*_{RR}/k^*_{RW}$. So, in practice, $k_{pol}[\text{dNTP}]/K_d$ could potentially be used to replace $k_{cat}[\text{dNTP}]/K_m$ (=$k^*$ for the two-step model) to estimate the initial discrimination of DNAP.

Theoretically speaking, one can not preclude the possibility that the time course of dNTP incorporation have to be fitted by double-exponential functions which gives two eigenvalues with close values ($|\lambda_2|$ may be one or two times as much as $|\lambda_1|$), though the single-exponential function performs very well in data fitting in all cases as we know from the literature. Even in such cases, we still have $A_0^{[2]}/A_1^{[2]} < |\lambda_1| < 2A_0^{[2]}/A_1^{[2]}$, which means that the relative deviation of the approximate Eq.(60) from the precise $|\lambda_1|$ is no more than 100%. So one can still conclude $k_{pol}[\text{dNTP}]/K_d \sim k^*$ with less than 100% deviation.

#### B. The specificity constant of the multi-step incorporation reaction

The real DNA replication process contains much more intermediate states. To interpret the observed Michaelis-Menten-like relation $v_{t\cdot s}^{pol} \approx k_{pol}[\text{dNTP}]/(K_d + [\text{dNTP}])$, one can chose more general reaction models such as the N-step model shown in Fig.S9 , rather than the two-step model in the preceding subsection. For the N-step model, the matrix method in Sec.III A can also be applied to give the smallest characteristic rate.

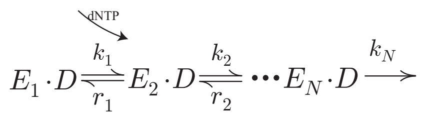


Fig.S9: The N-step reaction scheme of dNTP incorporation.

With similar notations and the same logic as the preceding subsection, we get the following equation for the N characteristic rates

$$\frac{1}{|\lambda_1|}+\frac{1}{|\lambda_2|}+...+\frac{1}{|\lambda_N|}=\frac{A_1^{[N]}}{A_0^{[N]}} \tag{61}$$

$A_1^{[N]}$ and $A_0^{[N]}$ are the first-order and zero-order coefficients respectively of the characteristic polynomials of the matrix.

As assumed in the preceding subsection, the smallest eigenvalue $|\lambda_1|$ is at least an order of magnitude smaller than others ($|\lambda_2|, ..., |\lambda_N|$). On the other hand, there are often not too many intermediate states ($N < 10$) observed in experiments or postulated in any reasonable theoretical model. So one can get

$$\frac{1}{|\lambda_1|}<\frac{A_1^{[N]}}{A_0^{[N]}}<\frac{2}{|\lambda_1|} \tag{62}$$

*i.e.*,

$$|\lambda_1|\sim\frac{A_0^{[N]}}{A_1^{[N]}}=\frac{k_{pol}^{[N]}[\text{dNTP}]}{K_d^{[N]}+[\text{dNTP}]} \tag{63}$$

Eq.(63) is a good approximation with less than 100% deviation from the precise $|\lambda_1|$.

$k_{pol}^{[N]}$ and $K_d^{[N]}$ are too complicated to be given here, but it can be shown that $k_{pol}^{[N]}/K_d^{[N]}=k^{*0}$. $k^*$ is defined by Eq.(38), with $k^*=k^{*0}[\text{dNTP}]$. On the other hand, experimental data fitting gives $|\lambda_1|=k_{pol}[\text{dNTP}]/(K_d+[\text{dNTP}])$, so we have $k_{pol}/K_d\approx k_{pol}^{[N]}/K_d^{[N]}$, leading to

$$\frac{k_{pol}[\text{dNTP}]}{K_d}\sim k^* \tag{64}$$

Similar to the discussion in the preceding subsection, the specificity constant $k_{pol}[\text{dNTP}]/K_d$ obtained by data fitting always overestimate $k^*$ but with relative deviation no more than 100%. So the relative deviation of the operationally defined fidelity $(k_{pol}[\text{dNTP}]/K_d)_{RR}/(k_{pol}[\text{dNTP}]/K_d)_{RW}$ from the true fidelity $k_{RR}^*/k_{RW}^*$ is less than 100%.

Fig.S10 shows the more realistic reaction schemes, considering DNAP binding/unbinding and even DNAP translocation. For generality, we assume there are $N^{'}$ states before dNTP binding.

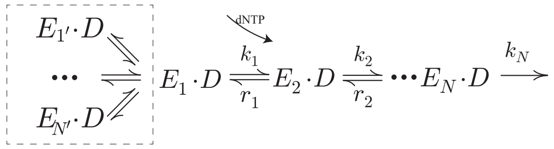


Fig.S10: The realistic reaction scheme of dNTP incorporation. Dashed rectangle represents the possible $N^{'}$ steps before dNTP binding.

Following the same logic above, one can calculate the smallest eigenvalue $|\lambda_1|$ for this complex reaction scheme approximately. For generality, we assume that $N^{'}+N\leq 20$ for any conceivable reaction models and also that $|\lambda_1|$ is at least one order of magnitude smaller than other eigenvalues. We thus have

$$\frac{1}{|\lambda_1|} < \frac{A_1^{[N',N]}}{A_0^{[N',N]}} < \frac{3}{|\lambda_1|}$$
$$\frac{A_0^{[N',N]}}{A_1^{[N',N]}} = \frac{k_{pol}^{[N',N]}[\text{dNTP}]}{K_d^{[N',N]} + [\text{dNTP}]} \tag{65}$$

$k_{pol}^{[N',N]}$ and $K_d^{[N',N]}$ can be rigoriously computed for any reaction model. They are too complicated to be given here, but it can be shown that $A_0^{[N',N]}/A_1^{[N',N]} = k^*/K_{t\cdot s}^{[N',N]}$ when [dNTP] $\to 0$. $K_{t\cdot s}^{[N',N]}$ is a function of the equilibrium constants of all the steps before dNTP binding. For instance, one can obtain $K_{t\cdot s} = 1 + k_{off}/k_{on}$ (here $k_{on} = k_{on}^0[E]$) for Fig.S11 and $K_{t\cdot s} = 1 + k'_{pe}/k'_{ep} + k^p_{off}/k^p_{on}$ (here $k^p_{on} = k^{p0}_{on}[E]$ ) for Fig.S12, which are similar to that obtained by the steady-state analysis. In particular, $K_{t\cdot s} \approx 1$ holds for the schemes in Fig.S11 and Fig.S12 , since $[E]$ can be set large enough in the transient-state assays ($k^p_{off} \ll k^p_{on}$) and $k'_{ep} \gg k'_{pe}$ can be made for the mutant $exo^-$-DNAP.

Since $|\lambda_1|$ can be well fitted by Michaelis-Menten equation $k_{pol}[\text{dNTP}]/(K_d + [\text{dNTP}])$, it leads to

$$\frac{k^*}{K_{t\cdot s}^{[N',N]}} < \frac{k_{pol}[\text{dNTP}]}{K_d} < \frac{3k^*}{K_{t\cdot s}^{[N',N]}} \tag{66}$$

Hence, the relative deviation of the operationally defined fidelity $(k_{pol}[\text{dNTP}]/K_d)_{RR}/(k_{pol}[\text{dNTP}]/K_d)_{RW}$ from the true fidelity $k^*_{RR}/k^*_{RW}$ is less than 200%, noting that $K_{t\cdot s}^{[N',N]}$ is exactly the same for either the terminal R or W.

Summarizing all the results given in this subsection and the preceding subsection, we conclude that since there are some assumptions made in the theoretical foundation of the transient-state assay, the specificity constant defined as $k_{pol}[\text{dNTP}]/K_d$ is not equal to the effective incorporation rate $k^*$, but it still offers a reasonable estimate, which gives an operationally-defined fidelity roughly equal to the true fidelity (with less than 200% relative deviation). In the experimental assays of the initial discrimation of DNAP (*i.e.*, $exo^-$-DNAP), both the steady-state assay and transient-state assay are always used (say, Ref.[6]) to guarantee that the specificity constants obtained by both methods agree with one another (no order of magnitude difference) in order to make a good estimate on $k^*$ and the true initial discrimination.

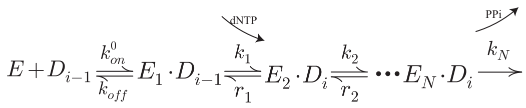

Fig.S11: The reaction scheme of the transient-state assay to measure the specificity constant of the nucleotide incorporation reaction of DNAP without exonuclease domain.

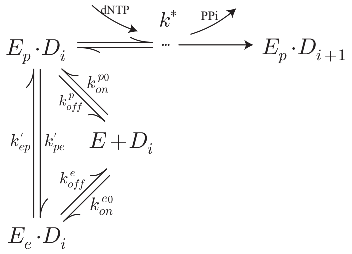


Fig.S12: The reaction scheme of the transient-state assay to measure the specificity constant of the nucleotide incorporation reaction of DNAP with deficient exonuclease domain.

#### C. The effective excision rate of $exo^+$-DNAP

To interpret the so-called excision rate measured in the single turnover assay, one can use the simplest scheme of the excision reaction of $exo^+$-DNAP as shown in Fig.S13.

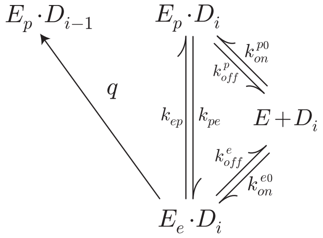


Fig.S13: The reaction scheme of the transient-state assay to measure the effective excision rate of $exo^+$-DNAP.

The matrix analysis (Sec.III A) can also be adopted to get the smallest characteristic rate of the excision reaction. The coefficient matrix is

$$\mathbf{M} = \begin{pmatrix} -(k_{on}^p + k_{on}^e) & k_{off}^p & k_{off}^e \\ k_{on}^p & -(k_{off}^p + k_{pe}) & k_{ep} \\ k_{on}^e & k_{pe} & -(k_{off}^e + k_{ep} + q) \end{pmatrix} \tag{67}$$

Here $k_{on}^p = k_{on}^{p0}[E]$, $k_{on}^e = k_{on}^{e0}[E]$. The matrix method gives the following equation of the three eigenvalues,

$$\frac{1}{|\lambda_1|} + \frac{1}{|\lambda_2|} + \frac{1}{|\lambda_3|} = \frac{A_1^{[3]}}{A_0^{[3]}} \tag{68}$$

in which

$$\begin{aligned} A_1^{[3]} &= (k_{on}^p + k_{on}^e)(q + k_{pe} + k_{ep}) + k_{off}^e k_{on}^p + k_{off}^p k_{on}^e + \epsilon \\ A_0^{[3]} &= k_{pe} q (k_{on}^p + k_{on}^e) + q k_{off}^p k_{on}^e \\ \epsilon &= k_{pe} q + k_{ep} k_{off}^p + k_{pe} k_{off}^e + q k_{off}^p + k_{off}^p k_{off}^e \end{aligned} \tag{69}$$

When $|\lambda_1|$ is more than one order of magnitude smaller than $|\lambda_2|, |\lambda_3|$, the experimentally observed smallest characteristic rate $v_{t\cdot s}^{exo}$ can be approximated as

$$v_{t\cdot s}^{exo} \approx \frac{A_0^{[3]}}{A_1^{[3]}} = \frac{k_{pe}q(k_{on}^p + k_{on}^e) + qk_{off}^p k_{on}^e}{(k_{on}^p + k_{on}^e)(q + k_{pe} + k_{ep}) + k_{off}^e k_{on}^p + k_{off}^p k_{on}^e + \epsilon} \tag{70}$$

This is a good approximation with less than 20% deviation from the precise $v_{t\cdot s}^{exo}$. Even when $|\lambda_1|$ is close to $|\lambda_2|$ and $|\lambda_3|$, we still have$A_0^{[3]}/A_1^{[3]} < |\lambda_1| < 3A_0^{[3]}/A_1^{[3]}$ which always holds. So the above approximate expression can also be used to estimate $v_{t\cdot s}^{exo}$, though it always underestimates the latter with less than 200% deviation. By comparing Eq.(70) and the true effective excision rate $\overline{r}$ defined by Eq.(42), we can show that they are roughly equal under some conditions. For example, the enzyme (DNAP) concentration can be set large enough in the experiments, which ensures $k_{on}^p > k_{off}^p, k_{on}^e > k_{off}^e, k_{on}^p > k_{pe}$ and $\epsilon \approx 0$ (compared to other terms in the denominator). If $\widetilde{k}_{ep} + q > \widetilde{k}_{pe}$ ($\widetilde{k}_{ep}$ and $\widetilde{k}_{pe}$ are given in Eq.(42)), we then have $v_{t\cdot s}^{exo} \sim \overline{r}$.

In this paper, we aim to compare the true fidelity $\mathscr{F} = \mathscr{F}_{ini} \cdot \mathscr{F}_{pro}$ (with $\mathscr{F}_{ini} \approx F_{ini}$ and $\mathscr{F}_{pro} \approx F_{pro}$) and the operationally defined fidelity $f_{t\cdot s} = f_{t\cdot s,ini} \cdot f_{t\cdot s,pro}$. As indicated in preceding sections, we acknowledge that the effective incorporation rate $k^*$ can be measured by the specificity constant in the steady-state assay (or roughly estimated by the specificity constant in the transient-state assay) and thus $F_{ini} \approx f_{t\cdot s,ini}$ is well estabilished. So $F/f_{t\cdot s} \approx F_{pro}/f_{t\cdot s,pro} = (1 + \overline{r}_{RW}/k_{WR}^*)/(1 + (v_{t\cdot s}^{exo})_{RW}/(k_{pol}[\mathrm{dNTP}]/K_d)_{WR}) \approx \overline{r}_{RW}/(v_{t\cdot s}^{exo})_{RW}$ (the last $\approx$ is reasonably assumed for efficient proofreading). $v_{t\cdot s}^{exo}$ can be approximately estimated by Eq.(70) or be precisely numerically computed by solving the eigenvalues of the matrix. Here we choose the latter for the comparison between $F$ and $f_{t\cdot s}$, which gives the Fig.S14 (Fig.9 in the main text). The red region indicates that there can be more than one order of magnitude difference between $\overline{r}$ and $v_{t\cdot s}^{exo}$ (*i.e.*, $F$ and $f_{t\cdot s}$) in some ranges of the key parameters $k_{pe}$ and $k_{ep}$.

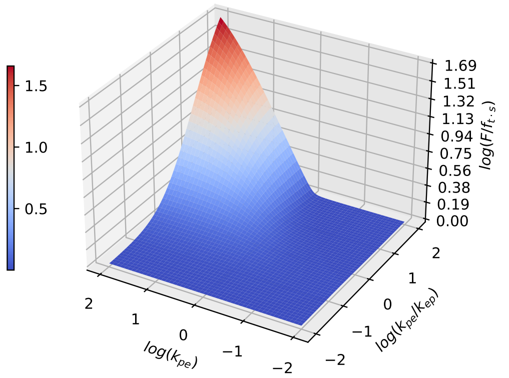


Fig.S14: The ratio $F/f_{t\cdot s} \approx \overline{r}_{RW}/(v_{t\cdot s}^{exo})_{RW}$ may become lager than 10 when $q, \widetilde{k}_{ep} < \widetilde{k}_{pe}$. The kinetic rates are set as $q = 1$, $k_{off}^e = 1$, $k_{off}^p = 0.1$, $k_{on}^p = 10^3$. $k_{on}^e$ is calculated by the thermodynamic constraint ($k_{on}^e = k_{on}^p k_{pe} k_{off}^e / k_{off}^p k_{ep}$). Details explanation see Fig.9 in main text

#### D. The reaction schemes with complex transfer processes

Since the two domains *Pol* and *Exo* of DNAP are often quite far apart, the intramolecular transfer of the primer terminal may include multiple substeps rather than a single step. The example of the more complex reaction scheme including two-step transfer process is shown in Fig.S15.

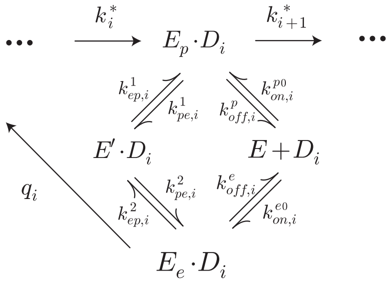


Fig.S15: The reaction scheme of the two-step intramolecular transfer process.

The effective rates can be obtained by using the FP method

$$\begin{aligned}
\overline{k} &= k^* \quad , \quad \overline{r} = \frac{\widetilde{k}_{pe} q}{\widetilde{k}_{ep} + q} \\
\widetilde{k}_{pe} &= k_{pe}^{''} + k_{p\to e} \quad , \quad \widetilde{k}_{ep} = k_{ep}^{''} + k_{e\to p} \\
k_{pe}^{''} &= \frac{k_{pe}^1 k_{pe}^2}{k_{ep}^1 + k_{pe}^2} \quad , \quad k_{ep}^{''} = \frac{k_{ep}^1 k_{ep}^2}{k_{ep}^1 + k_{pe}^2} \\
k_{p\to e} &= \frac{k_{off}^p k_{on}^e}{k_{on}^p + k_{on}^e} \quad , \quad k_{e\to p} = \frac{k_{off}^e k_{on}^p}{k_{on}^p + k_{on}^e}
\end{aligned} \tag{71}$$

$k_{pe}^{''}$ and $k_{ep}^{''}$ represent the effective intramolecular transfer rates. $k_{on}^p = k_{on}^{p0}[E]$ and $k_{on}^e = k_{on}^{e0}[E]$.

Using the transient-state assay, one can get the excision rate,

$$v_{t\cdot s}^{exo} \approx \frac{\widetilde{k}_{pe} q}{\widetilde{k}_{ep} + \widetilde{k}_{pe} + q + \epsilon_3 + \epsilon_4} \tag{72}$$

Where,

$$\begin{aligned}
\epsilon_3 &= \frac{1}{k_{on}^p + k_{on}^e} \left( q k_{pe}^{''} + q k_{off}^p + k_{off}^e k_{off}^p + k_{off}^e k_{pe}^{''} + k_{off}^p k_{ep}^{''} \right) \\
\epsilon_4 &= \frac{1}{k_{ep}^1 + k_{pe}^2} \left( k_{pe}^1 k_{ep}^2 + q k_{pe}^1 + q k_{p\to e} + k_{ep}^2 k_{p\to e} + k_{pe}^1 k_{e\to p} \right)
\end{aligned} \tag{73}$$

Apparently $v_{t\cdot s}^{exo}$ is quite different from $\overline{r}$, so it can not be used as $r_{RW}$ to estimate the proofreading efficiency $f_{pro}$ $(= \overline{r}_{RW}/k_{WR}^*)$. That is to say, the transient-state assay *per se* is not a generally reliable method to measure $f_{pro}$.

### IV. MORE COMPLEX MODELS INCLUDING DNAP TRANSLOCATION

#### A. The FP analysis

The reaction scheme including DNAP translocation is shown in Fig.S16.

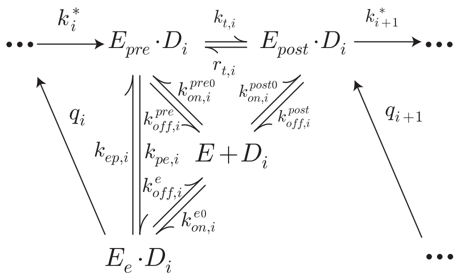

Fig.S16: The multi-step reaction scheme of *exo*$^+$-DNAP including the translocation step.

Here $k^*$ represents the effective rate of the multi-step incorporation process after dNTP binding. The corresponding integrated master equations are

$$\begin{aligned}
0 &= k_i^*\Gamma_{post,i-1} + r_{t,i}\Gamma_{post,i} - k_{t,i}\Gamma_{pre,i} + k_{ep,i}\Gamma_{e,i} - k_{pe,i}\Gamma_{pre,i} + k_{on,i}^{pre}\Gamma_{0,i} - k_{off,i}^{pre}\Gamma_{pre,i} \\
0 &= k_{t,i}\Gamma_{pre,i} - \left(r_{t,i} + \sum_{\alpha_{i+1}} k_{i+1}^*\right)\Gamma_{post,i} + \sum_{\alpha_{i+1}} q_{i+1}\Gamma_{e,i+1} + k_{on,i}^{post}\Gamma_{0,i} - k_{off,i}^{post}\Gamma_{post,i} \\
0 &= -q_i\Gamma_{e,i} - k_{ep,i}\Gamma_{e,i} + k_{pe,i}\Gamma_{pre,i} + k_{on,i}^{e}\Gamma_{0,i} - k_{off,i}^{e}\Gamma_{e,i} \\
0 &= -(k_{on,i}^{pre} + k_{on,i}^{post} + k_{on,i}^{e})\Gamma_{0,i} + k_{off,i}^{pre}\Gamma_{pre,i} + k_{off,i}^{post}\Gamma_{post,i} + k_{off,i}^{e}\Gamma_{e,i}
\end{aligned} \tag{74}$$

Where $\Gamma_0$ corresponds to the unbound state $E + D_i$. $k_{on}^a = k_{on}^{a0}[E]$, $a = pre, post, e$. By eliminating variables except $\Gamma_{post}$, one can obtain

$$0 = \overline{k}_i\Gamma_{post,i-1} + \sum_{\alpha_{i+1}} \overline{r}_{i+1}\Gamma_{post,i+1} - \left(\overline{r}_i + \sum_{\alpha_{i+1}} \overline{k}_{i+1}\right)\Gamma_{post,i} \tag{75}$$

which defines uniquely the effective incorporation rate $\overline{k}$ and the effective excision rate $\overline{r}$ (other reduction procedures eliminating $\Gamma_{post}$ can not give equations like Eq.(75) with properly defined effective rates). Here

$$\begin{gathered}
\overline{k} = k^*\left(1 - q\widetilde{k}_{pe}/\xi\right) \\
\overline{r} = q\eta/\xi \\
\widetilde{k}_{pe} = k_{pe} + k_{pre\to e} \\
\widetilde{k}_{ep} = k_{ep} + k_{e\to pre} \\
\eta = k_{post\to e}(k_t + k_{pre\to post} + \widetilde{k}_{pe}) + (r_t + k_{post\to pre})\widetilde{k}_{pe} \\
\xi = (q + k_{e\to post})(k_t + \widetilde{k}_{pe} + k_{pre\to post}) + \widetilde{k}_{ep}(k_t + k_{pre\to post}) \\
k_{a\to b} = k_{off}^a k_{on}^b/(k_{on}^{pre} + k_{on}^{post} + k_{on}^{e}),\ a, b = pre, post, e
\end{gathered} \tag{76}$$

here $k^*$ is defined by Eq.(38), $k^a_{on} = k^{a0}_{on}[\mathrm{E}]$.

It should be noted that in such cases the effective rates are much more complicated (*e.g.*, the exonuclease can affect the effective incorporation rate, so $\overline{k}$ is no longer $k^*$), but they may still satisfy the bio-relevant conditions ((a)-(c)) in Sec.I A. For example, it has been observed that DNAP translocation in the presence of the terminal $RR$ is very fast (*i.e.* $k_{t,RR} \gg k_{pe,RR}, k^{pre}_{off,RR}$), which leads to $(q\widetilde{k}_{pe}/\xi)_{RR} \ll 1$ and thus $\overline{k}_{RR} \approx k^*_{RR}$. For any terminal $\overline{k} < k^*$ always holds, which leads to $\overline{k}_{RW} < k^*_{RW}$. On the other hand, the initial discrimination $k^*_{RR}/k^*_{RW}$ (defined below) can be measured by the steady-state assay of *exo*$^-$-DNAP which always gives $k^*_{RR}/k^*_{RW} = (k_{cat}[\mathrm{dNTP}]/K_m)_{RR}/(k_{cat}[\mathrm{dNTP}]/K_m)_{RW} \gg 1$ (see Sec.II A). So the condition (a) $\overline{k}_{RR} \gg \overline{k}_{RW}$ still holds. Similarly, the steady-state assays of *exo*$^-$-DNAP also show $k^*_{WW} \approx 0$ (see Sec.II A). Since $\overline{k}_{WW} < k^*_{WW}$, so the condition (b) holds. As mentioned in Sec.I A, we restrict our discussion throughout this paper under the condition (c) which is the premise of the kinetic assays. We still assume $\overline{k}_{RR} \gg \overline{r}_{RW}, \overline{r}_{WR}$ here in order to compare our theoretical results to the results given by the kinetic assays. So one can safely use the approximate expressions in Eq.(29) to estimate the fidelity.

The total true fidelity can now be written as

$$\begin{aligned}
\mathscr{F}_i &\equiv \mathscr{F}_{ini,i} \cdot \mathscr{F}_{pro,i} (\approx F_i \equiv F_{ini,i} \cdot F_{pro,i}) \\
\mathscr{F}_{ini,i} &\approx F_{ini,i} \equiv \frac{k^*_{R_{i-1}R_i}}{k^*_{R_{i-1}W_i}} \\
\mathscr{F}_{pro,i} &\approx F_{pro,i} \equiv \frac{F_i}{F_{ini,i}} \\
&= \frac{\left(1 - q\widetilde{k}_{pe}/\xi\right)_{R_{i-1}R_i}}{\left(1 - q\widetilde{k}_{pe}/\xi\right)_{R_{i-1}W_i}} \left(1 + \frac{\overline{r}_{R_{i-1}W_i}}{\overline{k}_{W_iR_{i+1}}}\right) \\
&\approx \frac{q_{R_{i-1}W_i}}{k^*_{W_iR_{i+1}}} \frac{\left(1 - q\widetilde{k}_{pe}/\xi\right)_{R_{i-1}R_i}}{\left(1 - q\widetilde{k}_{pe}/\xi\right)_{W_iR_{i+1}}} \left(\frac{\eta}{\xi - q\widetilde{k}_{pe}}\right)_{R_{i-1}W_i}
\end{aligned} \tag{77}$$

Here $\mathscr{F}_{ini,i}$ is the initial discrimination of *exo*$^-$-DNAP. It can be esitmated by $F_{ini,i}$ which is calculated by Eq.(29) with the effective rates $\overline{k}$ defined in Eq.(76) while setting $q = 0$. Similarly, the total fidelity $\mathscr{F}_i$ can be estimated by $F_i$ which is given by Eq.(29) with the effective rates $\overline{k}$ defined in Eq.(76). For the $F_{pro,i}$, it is always assumed that $\overline{r}_{R_{i-1}W_i}/\overline{k}_{W_iR_{i+1}} > 1$ for efficient proofreading. Following the same logic in Sec.I A, one can reasonably assume that $k^*_{RR}(\approx \overline{k}_{RR})$ may be more than 100 times larger than $k^*_{RW}(> \overline{k}_{RW}), \overline{r}_{WR}, \overline{r}_{WR}$ to get,

$$\begin{aligned}
0.99F_i &< \mathscr{F}_i < 1.04F_i \\
F_{ini,i} &< \mathscr{F}_{ini,i} < 1.03F_{ini,i}
\end{aligned} \tag{78}$$

And the proofreading efficiency is,

$$\begin{aligned}
\mathscr{F}_{pro,i} &= \frac{\mathscr{F}_i/F_i}{\mathscr{F}_{ini,i}/F_{ini,i}} F_{pro,i} \\
0.96F_{pro,i} &< \mathscr{F}_{pro,i} < 1.04F_{pro,i}
\end{aligned} \tag{79}$$

So the relative deviations of the $F_i, F_{ini,i}, F_{pro,i}$ from the precise fidelity $\mathscr{F}_i, \mathscr{F}_{ini,i}, \mathscr{F}_{pro,i}$ is less than 10%.

#### B. The transient-state assay

The reaction scheme of measuring the specificity constant of the nucleotide incorporation reaction by the transient-state assay is shown in Fig.S17,

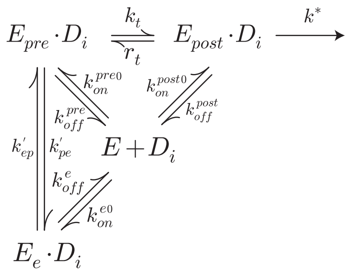

Fig.S17: The reaction scheme of the transient-state assay to measure the specificity constant of the incorporation reaction, considering DNAP translocation.

The specificity constant is given by the matrix analysis method

$$\left(\frac{k_{pol}[\text{dNTP}]}{K_d}\right)_{\alpha_{i-1}\alpha_i} \approx \frac{k^*_{\alpha_{i-1}\alpha_i}}{(K_{t\cdot s})_{\alpha_{i-2}\alpha_{i-1}}} \tag{80}$$

Here $\alpha = R, W$. $K_{t\cdot s} = 1 + r_t/k_t(1 + k'_{pe}/k'_{ep}) + k^{post}_{off}/k^{post}_{on}$, $k^{post}_{on} = k^{post0}_{on}[E]$. Following the same logic in Sec.(III A)(III B), the term on the right side of Eq.(80) always underestimates the measured specificity constant on the left side, with less than 200% relative deviation. Similarly, the steady-state assay also define another specificity constant, $(k_{cat}[\text{dNTP}]/K_m)_{\alpha_{i-1}\alpha_i} = k^*_{\alpha_{i-1}\alpha_i}/(K_{s\cdot s})_{\alpha_{i-2}\alpha_{i-1}}$, with $K_{s\cdot s} = 1 + r_t/k_t(1 + k'_{pe}/k'_{ep}) + k^{post}_{off}/k^{post}_{on}$, $k^{post}_{on} = k^{post,0}_{on}[\text{DNA}]$. Since $K_{t\cdot s}$ (or $K_{s\cdot s}$) is independent on the identity of $\alpha_i$, the initial discrimination can be measured by the steady-state assay with a very high precision or roughly estimated by the transient-state assay, *i.e.* $F_{ini} = f_{s\cdot s,ini}$ and $F_{ini} \approx f_{t\cdot s,ini}$ (with no order of magnitude difference).

In the discussion below, we assume $K_{t\cdot s} \approx 1 + r_t/k_t$, since the DNAP concentration[E] can be set large enough in the transient-state assays to ensure $k^{post}_{on} \gg k^{post}_{off}$ and the transfer rates $k'_{pe}$ and $k'_{ep}$ of the mutant DNAP often satisfies $k'_{ep} \gg k'_{pe}$.

The reaction scheme for the transient-state assay of the excision velocity is shown in Fig.S18.

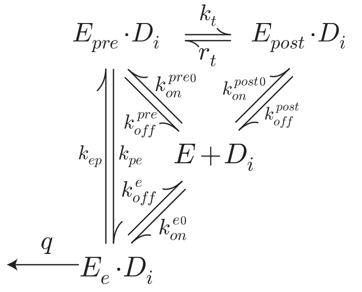

Fig.S18: The reaction scheme of the transient-state assay to measure the excision rate, considering DNAP translocation.

The coefficient matrix is

$$\mathbf{M} = \begin{pmatrix} -(k_{on}^{pre}+k_{on}^{post}+k_{on}^{e}) & k_{off}^{pre} & k_{off}^{post} & k_{off}^{e} \\ k_{on}^{pre} & -(k_{off}^{pre}+k_{pe}+k_t) & r_t & k_{ep} \\ k_{on}^{post} & k_t & -(k_{off}^{post}+r_t) & 0 \\ k_{on}^{e} & k_{pe} & 0 & -(k_{off}^{e}+k_{ep}+q) \end{pmatrix} \tag{81}$$

By using the matrix analysis, one gets the excision velocity

$$v_{t\cdot s}^{exo} \approx q\eta/\kappa \tag{82}$$

Here $\eta$ is defined in Eq.(76).

$$\kappa = (1 + r_t/k_t)(\xi - q\widetilde{k}_{pe}) + \epsilon_1 + \epsilon_2 \tag{83}$$

$\xi, \widetilde{k}_{pe}$ are defined in Eq.(76)

$$\begin{aligned} \epsilon_1 &= k_t k_{post\to e} + r_t(k_{pre\to e} + k_{pe}) + q(k_{pe} + k_{pre\to e} + k_{post\to e}) + k_{pe}(k_{post\to e} + k_{post\to pre}) + k_{off}^{pre} k_{post\to e} \\ \epsilon_2 &= \Big(k_t(q k_{off}^{post} + k_{ep} k_{off}^{post} + k_{off}^{e} k_{off}^{post}) + r_t(q k_{pe} + k_{off}^{e} k_{pe} + q k_{off}^{pre} + k_{ep} k_{off}^{pre} + k_{off}^{e} k_{off}^{pre}) \\ &+ q(k_{pe} k_{off}^{post} + k_{off}^{pre} k_{off}^{post}) + k_{ep} k_{off}^{pre} k_{off}^{post} + k_{pe} k_{off}^{e} k_{off}^{post} + k_{off}^{e} k_{off}^{post} k_{off}^{pre}\Big) \Big/ (k_{on}^{e} + k_{on}^{pre} + k_{on}^{post}) \end{aligned} \tag{84}$$

Follow the same logic in Sec.III C , the approximate Eq.(82) always underestimates the precise $v_{t\cdot s}^{exo}$ but with less than 300% relative deviation.

The proofreading efficiency $f_{pro}$ defined by the transient-state assay is

$$f_{t\cdot s,pro,i} = 1 + \frac{v_{t\cdot s,R_{i-1}W_i}^{exo}}{(k_{pol}[\text{dNTP}]/K_d)_{W_iR_{i+1}}} \approx 1 + \frac{q_{R_{i-1}W_i}}{k_{W_iR_{i+1}}^{*}}\left(\frac{\eta K_{t\cdot s}}{\kappa}\right)_{R_{i-1}W_i} \approx \frac{q_{R_{i-1}W_i}}{k_{W_iR_{i+1}}^{*}}\left(\frac{\eta K_{t\cdot s}}{\kappa}\right)_{R_{i-1}W_i} \tag{85}$$

This is generally different from the $F_{pro}$ defined by Eq.(77). They might be approximately equal only under some conditions. Below we give an example.

Directly comparing the Eq.(77) and Eq.(85), one can find that the $F_{pro,i} \approx f_{t\cdot s,pro,i}$ when

$$\left(1 - q\widetilde{k}_{pe}/\xi\right)_{R_{i-1}R_i} \approx 1 \tag{86}$$

$$\left(1 - q\widetilde{k}_{pe}/\xi\right)_{W_iR_{i+1}} \approx 1 \tag{87}$$

$$\left(\frac{1}{\xi - q\widetilde{k}_{pe}}\right)_{R_{i-1}W_i} \approx \left(\frac{K_{t\cdot s}}{\kappa}\right)_{R_{i-1}W_i} \tag{88}$$

These equations hold if the kinetic rates satisfy the following conditions,

$(1)k_{t,RR} \gg k^{pre}_{off,RR}, k_{pe,RR}$

$(2)k_{t,WR} \gg k^{pre}_{off,WR}, k_{pe,WR}$

$(3)k_{t,RW} \gg k^{pre}_{off,RW}, k_{pe,RW}$ and $r_{t,RW} \gg k^{post}_{off,RW}$.

$(4)\widetilde{k}_{ep,RW} \gg \widetilde{k}_{pe,RW}$ and $q_{RW} \gg \widetilde{k}_{pe,RW}$.

Here $\gg$ means more than one order of magnitude higher.

From the condition (1), one has $0.2k_{t,RR} > k_{pe,RR} + k^{pre}_{off,RR} > \widetilde{k}_{pe,RR}$. Combining $\xi > q(k_t + \widetilde{k}_{pe})$ from Eq.(76), one has $\left(\xi/q\widetilde{k}_{pe}\right)_{RR} > 6$, *i.e.*, $\left(1 - q\widetilde{k}_{pe}/\xi\right)_{RR} > 0.83$, and Eq.(86) holds. From the condition (2), one can similarly get $\left(1 - q\widetilde{k}_{pe}/\xi\right)_{WR} > 0.83$ and thus Eq.(87) holds.

From the condition (3)(4), one can compare each term of $\epsilon_{1,RW}$ and $\left((1 + r_t/k_t)(\xi - q\widetilde{k}_{pe})\right)_{RW}$ in $\kappa_{RW}$ (see Eq.(83)) to get $\epsilon_{1,RW} < 0.1\left((1 + r_t/k_t)(\xi - q\widetilde{k}_{pe})\right)_{RW}$. On the other hand, it has been shown that $k^a_{on} = k^{a0}_{on}[\mathrm{DNA}](a = pre, post, e)$, [DNA] can be set in the experiments to ensure $k^a_{on} \gg k^a_{off}$ and $k^{pre}_{on} + k^{post}_{on} + k^e_{on} \gg k_{pe}$. These conditions leads to $\epsilon_{2,RW} < 0.1\left((1 + r_t/k_t)(\xi - q\widetilde{k}_{pe})\right)_{RW}$. So $\left((1 + r_t/k_t)(\xi - q\widetilde{k}_{pe})\right)_{RW} < \kappa_{RW} < 1.2\left((1 + r_t/k_t)(\xi - q\widetilde{k}_{pe})\right)_{RW}$. And it has been shown that $(K_{t\cdot s})_{R_{i-1}W_i} \approx (1 + r_t/k_t)_{R_{i-1}W_i}$. So $\left(\xi - q\widetilde{k}_{pe}\right)_{RW} < (\kappa/K_{t\cdot s})_{RW} < 1.2\left(\xi - q\widetilde{k}_{pe}\right)_{RW}$, which leads to Eq.(88).

Taking all the above estimates together, one can get $0.69 < F_{pro}/f_{t\cdot s,pro} < 1.20$, and the relative deviation of $f_{t\cdot s,pro}$ from $F_{pro}$ is less than 31%.

It should be noted that some of the above conditions may be unreasonable for real DNAPs. For example, the terminal mismatch or buried mismatch may slow down DNAP translocation, *i.e.*, $k_{t,RW} < k_{pe,RW}$ and $k_{t,WR} < k_{pe,WR}$ (the conditions (2) and (3) are violated). In such cases, $f_{t\cdot s,pro}$ and $F_{pro}$ can be totally different, which is shown by the following numerical examples.

First we consider the cases that the kinetic rates satisfy the condition (1)(2) but violate (3) or (4). This means Eq.(86) and Eq.(87) hold, but Eq.(88) does not hold. In such case, since $F_{ini} \approx f_{t\cdot s,ini}$ is well estabilished, $F/f_{t\cdot s} \approx F_{pro}/f_{t\cdot s,pro} \approx \left(q\eta/\left(\xi - q\widetilde{k}_{pe}\right)\right)_{RW} \Big/ (v^{exo}_{t\cdot s}K_{t\cdot s})_{RW}$. Here $v^{exo}_{t\cdot s}$ is precisely computed by numerically solving the eigenvalues of the matrix, and $K_{t\cdot s} \approx 1 + r_t/k_t$ as discussed above. Here we show two examples, one in Fig.S19 (violating the condition (3). See also Fig.11 in the main text) and the other in Fig.S20 (violating the condition (4). See also Fig.12 in the main text). The red regions indicate that there can be more than one order of magnitude difference between $F$ and $f_{t\cdot s}$.

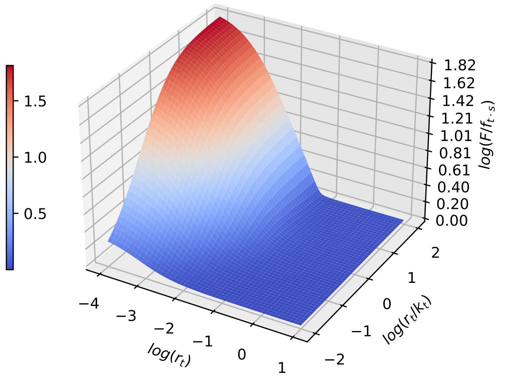


Fig.S19: The ratio $F/f_{t\cdot s}$ becomes larger than 10 when $k_t$ and $r_t$ get small enough. The kinetic rates (for terminal $RW$) : $q = 1$, $k_{pe} = 0.1$, $k_{ep} = 1$, $k^{e}_{off} = 1$, $k^{pre}_{off} = 0.1$, $k^{post}_{off} = 0.001$, $k^{pre}_{on} = 10^3$. $k^{e}_{on}$ and $k^{post}_{on}$ are calculated by thermodynamic constraints ($k^{e}_{on} = k^{p}_{on}k_{pe}k^{e}_{off}/k^{pre}_{off}k_{ep} = 10^3$, $k^{post}_{on} = k^{pre}_{on}k_t k^{post}_{off}/k^{pre}_{off}r_t$). The binding rates are always large enough when adjusting $k_t$ and $r_t$ to ensure $k^{a}_{on} > k^{a}_{off}, a = pre, post, e$ and $k^{pre}_{on} > k_{pe}$.

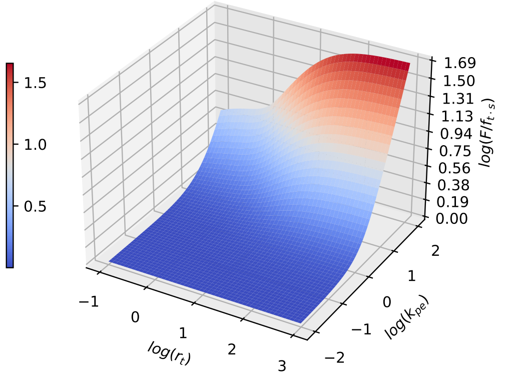


Fig.S20: The ratio $F/f_{t\cdot s}$ may become larger than 10 when $k_{pe}$ gets large enough. The kinetic rates (for terminal $RW$) : $q = 1$, $k_{ep} = 1$, $k^{e}_{off} = 1$, $k^{pre}_{off} = 0.1$, $k^{post}_{off} = 0.1$, $k_t = 10$, $k^{pre}_{on} = 10^3$. $k^{e}_{on}$ and $k^{post}_{on}$ are calculated by thermodynamic constraints($k^{e}_{on} = k^{p}_{on}k_{pe}k^{e}_{off}/k^{pre}_{off}k_{ep}$, $k^{post}_{on} = k^{pre}_{on}k_t k^{post}_{off}/k^{pre}_{off}r_t$). The binding rates are always large enough when adjusting $k_{pe}$ and $r_t$ to ensure $k^{a}_{on} > k^{a}_{off}, a = pre, post, e$ and $k^{pre}_{on} > k_{pe}$.

However, the condition (2) (or equivalently, Eq.(87)) may also not hold for real DNAPs. Since the buried mismatch may slow down DNAP translocation, $\left(1 - q\widetilde{k}_{pe}/\xi\right)_{W_iR_{i+1}} \ll 1$ may hold if $\widetilde{k}_{pe,W_iR_{i+1}} \gg k_{t,W_iR_{i+1}}, k_{pre\to post,W_iR_{i+1}}$ and $q_{W_iR_{i+1}} \gg \widetilde{k}_{ep,W_iR_{i+1}}$. On the other hand, the kinetic rates of the excision pathway for the terminal $W_iR_{i+1}$ are present in $F_{pro}$ (in the factor $1/\left(1 - q\widetilde{k}_{pe}/\xi\right)_{W_iR_{i+1}}$, see Eq.(77)) but totally absent from $f_{t\cdot s,pro}$ (see Eq.(85)). So, $F_{pro}$ can become larger, if $\left(1 - q\widetilde{k}_{pe}/\xi\right)_{W_iR_{i+1}}$ gets smaller than 1 while other kinetic rates for terminal $R_{i-1}W_i$ are fixed. That is to say, $F/f_{t\cdot s}$ may become orders of magnitude larger than that shown by the red regions in Fig.S19 and Fig.S 20.

### V. THE SUGGESTED SINGLE-MOLECULE ASSAY BASED ON FP ANALYSIS

As demonstrated in this paper, neither the steady-state assay nor the transient-state assay can directly measure the effective incorporation rates and the effective excision rates. Here we suggest a possible single-molecule approach, based on the FP analysis, to directly measure the effective rates.

#### A. The single-molecule assay without considering DNAP translocation

In a typical single-molecule experiment, some states of the enzyme-substrate system can be distinguished by techniques such as smFRET[7–10](*e.g.* for KF, the states $E_p \cdot D$, $E_e \cdot D$ and $E + D$ have been resolved by smFRET in Ref.[9]). Supposing that some key states of the enzyme-substrate system can be well resolved by some techniques while the normal activity of the enzyme is not significantly perturbed, we suggest a single-molecule assay as follows.

##### 1. *Measuring the effective incorporation rate of exo$^-$-DNAP*

We first introduce the single-molecule assay to measure the effective incorporation rate $\overline{k}$ of *exo*$^-$-DNAP without considering DNAP translocation. If the state $E_p \cdot D_i$ and $E_p \cdot D_{i+1}$ in Fig.S21 can be properly identified, the following single-molecule experiment can be done to measure the effective incorporation rates.

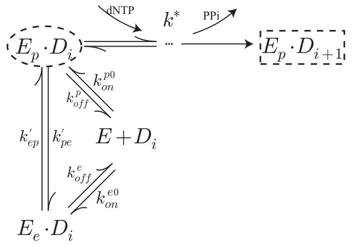


Fig.S21: The reaction scheme of the single-molecule assay to measure the effective incorporation rate of DNAP with deficient exonuclease domain. The dashed circle represents the starting point, and the dashed rectangle represents the ending point.

1. Initiate the nucleotide incorporation reaction by adding *exo*$^-$-DNAP and dNTP to the substrate $D_i$ and begin to record the state-switching trajectory of a single enzyme-DNA complex. Here dNTP can be dRTP or dWTP, and the primer terminal can be matched(R) or mismatched(W). When a single DNAP is captured by the substrate DNA, it can catalyze the incorporation of one or more nucleotides, depending on the template sequence context and the dNTP used. Then one can select a particular time window from the recorded trajectory, starting from the first-arrival at $E_p \cdot D_i$ (the starting point) and ending at the first-arrival at $E_p \cdot D_{i+1}$ (the ending point). In the following, either the starting point or the ending point means the first-arrival point.

2. In this time window, the system may make multiple visits to $E_p \cdot D_i$. Count the total time the system resides at $E_p \cdot D_i$. This so-called residence time may be clearly measured under low concentrations of dNTP.

3. Collect sufficient samples to get the averaged residence time $\Gamma_{p,i}$, which gives directly the required effective incorporation rate $k^*_{\alpha_i\alpha_{i+1}} = 1/\Gamma_{p,\alpha_i}$. Here $k^*_{\alpha_i\alpha_{i+1}} = k^{*0}_{\alpha_i\alpha_{i+1}}[d\alpha_{i+1}TP]$, $\alpha$ =A,T,G,C.

It is easy to prove $k^*_{\alpha_i\alpha_{i+1}} = 1/\Gamma_{p,\alpha_i}$. Since we choose the state $E_p \cdot D_i$ as the starting point and $E_p \cdot D_{i+1}$ as the ending point, the corresponding integrated master equations are,

$$
\begin{aligned}
-1 &= k_{on}^{p}\Gamma_0 + k_{ep}\Gamma_e - (k_1 + k_{pe} + k_{off}^{p})\Gamma_p \\
0 &= k_{off}^{p}\Gamma_p + k_{off}^{e}\Gamma_e - (k_{on}^{p} + k_{on}^{e})\Gamma_0 \\
0 &= k_{pe}\Gamma_p + k_{on}^{e}\Gamma_0 - (k_{ep} + k_{off}^{e})\Gamma_e \\
0 &= k_1\Gamma_1 + r_2\Gamma_3 - (k_2 + r_1)\Gamma_2 \\
&... \\
0 &= k_{j-1}\Gamma_{j-1} + r_j\Gamma_{j+1} - (k_j + r_{j-1})\Gamma_j \\
&... \\
0 &= k_{N-1}\Gamma_{N-1} - (k_N + r_{N-1})\Gamma_N
\end{aligned}
\tag{89}
$$

Here $\Gamma_2, \Gamma_3 ... \Gamma_N$ represent the intermediate states of the multi-step incorporation process (omitted in the figure). We choose the state $E_p \cdot D_i$ as the starting point in the data analysis, so we have $\int_0^\infty \frac{d}{dt} P_p dt = -1$ in the first equation. One can easily show $\Gamma_p = 1/k^*$, $k^*$ is the effective incorporation rate defined in Eq.(42).

The advantage of this single-molecule analysis is model-independence. Since $k_{i+1}^* = 1/\Gamma_{p,i}$ holds in general, the measurement of $\Gamma_{p,i}$ does not depend on any hypothesis about the details of the reaction scheme (in fact, steps after dNTP binding are often unclear).

#### *2. Measuring the effective excision rate of exo$^+$-DNAP*

The single-molecule assay of $\overline{r}$ follows the same logic, with the reaction schemes Fig.S22. The experiment is initiated by adding *exo*$^+$-DNAP to the substrate DNA. If the states $E_p \cdot D_i$, $E_e \cdot D_i$, $E + D_i$ and $E_p \cdot D_{i-1}$ in Fig.S22 can be well resolved, the state-switching trajectory between the starting point $E_p \cdot D_i$ and the ending point $E_p \cdot D_{i-1}$ can be recorded. Then the averaged residence time $\Gamma_{p,i}$ at $E_p \cdot D_i$ can be obtained, which gives $\overline{r}_i = 1/\Gamma_{p,i}$. Sometimes, however, the excision may occur without visiting $E_p \cdot D$. The trajectory recorded in such cases are not taken for the averaging.

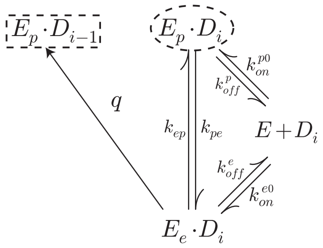


Fig.S22: The reaction scheme of the single-molecule assay to measure the effective excision rate of *exo*$^+$-DNAP. The dashed circle represents the starting point, and the dashed rectangle represents the ending point.

Since we choose the state $E_p \cdot D_i$ as the starting point and $E_p \cdot D_{i-11}$ as the ending point, the following integrated master equations are,

$$
\begin{aligned}
-1 &= k_{on}^{p}\Gamma_0 + k_{ep}\Gamma_e - (k_{pe} + k_{off}^{p})\Gamma_p \\
0 &= k_{off}^{p}\Gamma_p + k_{off}^{e}\Gamma_e - (k_{on}^{p} + k_{on}^{e})\Gamma_0 \\
0 &= k_{pe}\Gamma_p + k_{on}^{e}\Gamma_0 - (k_{ep} + k_{off}^{e} + q)\Gamma_e
\end{aligned}
\qquad (90)
$$

$\int_0^\infty \frac{d}{dt}P_p dt = -1$ in the first equation corresponds to the starting point $E_p \cdot D_i$. One can get $\Gamma_p = 1/\overline{r}$. Here $\overline{r}$ is the effective excision rate defined in Eq.(42). This analysis can also apply to more complex reaction schemes and one can always get $\overline{r}_i = 1/\Gamma_{p,i}$.

### B. The single-molecule assay with considering DNAP translocation

The single-molecule assay can still be applied when considering DNAP translocation. First, if the states *pre* and *post* cannot be distinguished in the experiment, indicating that the translocation is a fast process, the assays presented in the preceding section (Sec.V A) can be used. Second, if the translocation is a relatively slow process, either *pre* or *post* can be directly observed (*e.g.* for Dpo4 polymerase by smFRET [11]), then the single-molecule assay should be used with modification.

#### 1. The single-molecule assay to measure the effective incorporation rate of exo$^-$-DNAP

As shown by Eq.(76), the effective incorporation rate $\overline{k}$ is no longer $k^*$ but $k^*(1 - q\widetilde{k}_{pe}/\xi)$. It's hard to obtain this effective rate directly in a single measurement, since it consists of both the polymerase and the exonuclease contributions. Fortunately we can measure the factors $k^*$ and $1 - q\widetilde{k}_{pe}/\xi$ separately.

The measurement of $k^*$ is basically the same as that given in the preceding section (Sec.V A). The reaction scheme is shown in Fig.S23. The experiment is initiated by mixing DNAP and dNTP to the single molecule DNA. The time trajectory between the starting point $E_{post} \cdot D_{i-1}$ and the ending point $E_{pre} \cdot D_i$ is selected, if $E_{post} \cdot D_{i-1}$, $E_{pre} \cdot D_i$ and other states can be well distinguished. Then the average residence time at $E_{post} \cdot D_{i-1}$ gives $k_i^* = 1/\Gamma_{post,i-1}$ or $k_i^{*0} = 1/(\Gamma_{post,i-1}[d\alpha_i TP])$, with which one can calculate $F_{ini}$.

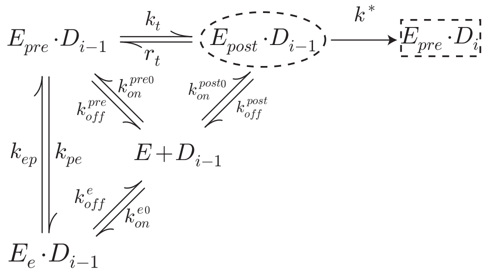

Fig.S23: The reaction scheme of the suggested single-molecule experiment to measure $k^*$. The dashed circle represents the starting point, and the dashed rectangle represents the ending point.

To prove the above relation, we write the corresponding integrated master equations as follows

$$\begin{aligned}
0 &= r_t\Gamma_{post} + k_{ep}\Gamma_e + k_{on}^{pre}\Gamma_0 - \left(k_t + k_{pe} + k_{off}^{pre}\right)\Gamma_{pre} \\
-1 &= k_t\Gamma_{pre} + k_{on}^{post}\Gamma_0 - \left(r_t + k^* + k_{off}^{post}\right)\Gamma_{post} \\
0 &= k_{pe}\Gamma_{pre} + k_{on}^{e}\Gamma_0 - \left(k_{ep} + k_{off}^{e}\right)\Gamma_e \\
0 &= k_{off}^{pre}\Gamma_{pre} + k_{off}^{post}\Gamma_{post} + k_{off}^{e}\Gamma_e - (k_{on}^{pre} + k_{on}^{post} + k_{on}^{e})\Gamma_0
\end{aligned} \tag{91}$$

Solving these equations, one can easily get $\Gamma_{post} = 1/k^*$.

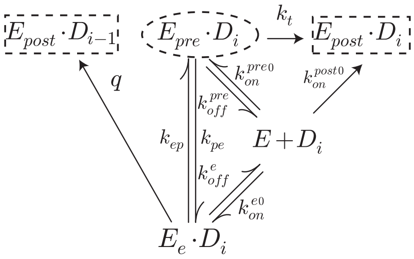


Fig.S24: The suggested reaction scheme to interpret the factor $q\widetilde{k}_{pe}/\xi$, based on FP analysis. The dashed circle represents the starting point, and the dashed rectangle represents the ending point.

The logic to measure $q\widetilde{k}_{pe}/\xi$ is given below, as shown in Fig.S24.

1. The experiment is initiated by mixing DNAP with DNA.

2. Record the state-switching trajectory of the complex. It may go directly to $E_{post} \cdot D_{i-1}$ without visiting $E_{pre} \cdot D_i$. Or it may arrive at $E_{pre} \cdot D_i$ *via* whatever pathway before the excision, and then go to $E_{post} \cdot D_{i-1}$ (with or without visiting $E_{post} \cdot D_i$). We collect trajectories of the latter case, and denote $E_{pre} \cdot D_i$ as the starting point (it may be visited repeatedly), $E_{post} \cdot D_i$ and $E_{post} \cdot D_{i-1}$ as the two ending points.

3. Select all the windows from the trajectories, which are between the starting point and either ending point. The windows are classified in two types, *i.e.* between $E_{pre} \cdot D_i$ and $E_{post} \cdot D_i$, or between $E_{pre} \cdot D_i$ and $E_{post} \cdot D_{i-1}$ without visiting $E_{post} \cdot D_i$.

4. Count the total number of either type of window $n_{post,i}$, $n_{post,i-1}$, and one gets $n_{post,i-1}\Big/\left(n_{post,i-1} + n_{post,i}\right) = (q\widetilde{k}_{pe}/\xi)_i$.

To prove the above relation, we write the integrated master equations as follows

$$\begin{aligned}
&-1 = k_{ep}\Gamma_e + k_{on}^{pre}\Gamma_0 - \left(k_t + k_{pe} + k_{off}^{pre}\right)\Gamma_{pre} \\
&0 = k_{pe}\Gamma_{pre} + k_{on}^{e}\Gamma_0 - \left(k_{off}^{e} + k_{ep} + q\right)\Gamma_e \\
&0 = k_{off}^{pre}\Gamma_{pre} + k_{off}^{e}\Gamma_e - (k_{on}^{pre} + k_{on}^{post} + k_{on}^{e})\Gamma_0 \\
&P_{post,i-1}(t \to \infty) = k_t\Gamma_{pre} + k_{on}^{post}\Gamma_0, \\
&P_{post,i}(t \to \infty) = q\Gamma_e
\end{aligned} \tag{92}$$

here $P_{post,i-1}(t \to \infty)$ and $P_{post,i}(t \to \infty)$ represent the probabilities of the first arrival at the two ending points

$E_{post} \cdot D_{i-1}$ and $E_{post} \cdot D_i$ respectively. From Eq.(92) one can get

$$\frac{P_{post,i-1}(t \to \infty)}{P_{post,i-1}(t \to \infty) + P_{post,i}(t \to \infty)} = \frac{q\widetilde{k}_{pe}}{\xi} \tag{93}$$

It should be noted that the reaction scheme Fig.S24 only offers the intuitive interpretation of $q\widetilde{k}_{pe}/\xi$ and can not be implemented in practice. Instead, the realistic reaction scheme is Fig.S25. Nevertheless, the above interpretation provides the method to analyze the experimental data.

#### 2. *The single-molecule assay to measure the effective excision rate of exo$^+$-DNAP*

The reaction scheme for the measurement of $\overline{r}$ is shown in Fig.S25. The experiment is initiated by adding DNAPs to the single molecule DNA. The time window selected from the trajectory is between the starting point $E_{post} \cdot D_i$ and the ending point $E_{post} \cdot D_{i-1}$, if $E_{post} \cdot D_i$, $E_{post} \cdot D_{i-1}$ and other states can be well distinguished. Then the average residence time at $E_{post} \cdot D_i$ gives $\overline{r}_i = 1/\Gamma_{post,i}$. Similarly, the trajectory recorded without visiting $E_{post} \cdot D_i$ are not taken for the averaging.

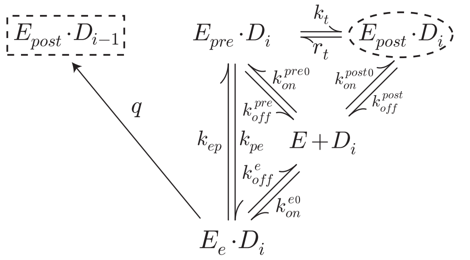


Fig.S25: The reaction scheme of the suggested single-molecule assay to measure $\overline{r}$. The dashed circle represents the starting point, and the dashed rectangle represents the ending point.

To prove the above relation, we solve the corresponding integrated master equations

$$\begin{aligned}
0 &= r_t\Gamma_{post} + k_{ep}\Gamma_e + k_{on}^{pre}\Gamma_0 - \left(k_t + k_{pe} + k_{off}^{pre}\right)\Gamma_{pre} \\
-1 &= k_t\Gamma_{pre} + k_{on}^{post}\Gamma_0 - \left(r_t + k_{off}^{post}\right)\Gamma_{post} \\
0 &= k_{pe}\Gamma_{pre} + k_{on}^{e}\Gamma_0 - \left(k_{ep} + k_{off}^{e} + q\right)\Gamma_e \\
0 &= k_{off}^{pre}\Gamma_{pre} + k_{off}^{post}\Gamma_{post} + k_{off}^{e}\Gamma_e - (k_{on}^{pre} + k_{on}^{post} + k_{on}^{e})\Gamma_0
\end{aligned} \tag{94}$$

and get $\Gamma_{post} = 1/(q\eta/\xi) = 1/\overline{r}$. Here $\overline{r}$, $\eta$, $\xi$ are defined in Eq.(76).


[1] Q. S. Li, P. D. Zheng, Y. G. Shu, Z. C. Ou-Yang, and M. Li, Physical Review E **100**, 012131 (2019).

[6] J. G. Bertram, K. Oertell, J. Petruska, and M. F. Goodman, Biochemistry **49**, 20 (2010).

[3] A. R. Fersht, *Enzyme Structure and Mechanism* (W.H.Freeman & Co Ltd., 1985), 2nd ed.

[4] B. M. Wingert, E. E. Parrott, and S. W. Nelson, Biochemistry **52**, 7723 (2013).

[5] A. K. Vashishtha and R. D. Kuchta, Biochemistry **54**, 240 (2015).

[6] J. G. Bertram, K. Oertell, J. Petruska, and M. F. Goodman, Biochemistry **49**, 20 (2010).

[7] Y. Santoso, C. M. Joyce, O. Potapova, L. Le Reste, J. Hohlbein, J. P. Torella, N. D. Grindley, and A. N. Kapanidis, Proc. Natl. Acad. Sci. U.S.A. **107**, 715 (2010).

[8] J. Hohlbein, L. Aigrain, T. D. Craggs, O. Bermek, O. Potapova, P. Shoolizadeh, N. D. Grindley, C. M. Joyce, and A. N. Kapanidis, Nat. Commun. **4**,1 (2013).

[9] R. P. Markiewicz, K. B. Vrtis, D. Rueda, and L. J. Romano, Nucleic Acids Res. **40**, 7975 (2012).

[10] R. Lamichhane, S. Y. Berezhna, J. P. Gill, E. Van Der Schans, and D. P. Millar, J. Am. Chem. Soc. **135**, 4735 (2013).

[11] A. Brenlla, R. P. Markiewicz, D. Rueda, and L. J. Romano, Nucleic Acids Res. **42**, 2555 (2014).
